# Recurrent Spiking Neural Networks for Continuous Motor Control: A Biologically Plausible Model of Motor Cortex

**DOI:** 10.64898/2026.10.26.756490

**Authors:** Hiroto Imamura, Fumiya Imamura, Jun Igarashi, Yoshikazu Isomura, Riichiro Hira

**Author notes:** **Corresponding authors:** Riichiro Hira, M.D., Ph.D., Hiroto Imamura, M.D.

## Abstract

Motor cortex acquires skilled movements through recurrent spiking dynamics and experience-dependent synaptic plasticity, yet how such circuits learn continuous motor control using biologically local signals remains unclear. Here, we developed a recurrent spiking neural network model of motor cortex that addresses both the computational and biological requirements of motor learning. We first analytically derived the recurrent firing-rate state directly from synaptic input and trained the network by implicit differentiation, thereby avoiding backpropagation through long spike trajectories. We then introduced State-Carry Initialization, which enables closed-loop motor control by continuously carrying the ongoing recurrent state across successive actions. Building on this formulation, we devised a learning rule termed Feedback-Spike Nudging (FSN), in which teaching signals are delivered as spikes through a feedback pathway. Synaptic changes depend only on presynaptic activity and the feedback-evoked change in postsynaptic firing, consistent with biological mechanisms of synaptic plasticity in motor cortex. The resulting networks learned continuous motor control through both supervised and reinforcement learning, which could be mapped respectively onto interactions of motor cortex with the cerebellum and basal ganglia. Moreover, the FSN learning rule aligned with experimental observations of structural spine plasticity in mouse motor cortex. Thus, our study lays a foundation for recurrent spiking models of motor cortex that are biologically grounded at both the functional and anatomical levels.

## Introduction

Motor cortex learns and executes skilled voluntary movements while maintaining an internal state in ongoing spiking activity [1–9]. Recurrent spiking neural networks (RSNNs) are therefore a useful model class [10]. Spikes provide the communication signal, recurrent activity maintains the internal state, and the circuit can transform ongoing sensory input into action. However, using RSNNs as models of motor cortex for movement generation presents two computational challenges. The first is the difficulty of learning to generate the desired movement. Surrogate-gradient backpropagation through time (BPTT) unrolls the membrane-voltage trajectory and spike events across the internal simulation, replaces the spike threshold with a smooth derivative, and propagates credit backward through that trajectory [11–15]. Its runtime and memory therefore grow with the number of internal spiking steps, and its gradient depends on a single finite spike realization. Exact spike-time gradients also depend on the simulated spike trajectory [16,17]. The second is the difficulty of continuous control. Because current behavior depends on earlier observations, the recurrent state must be carried from one action to the next. To overcome these two challenges, an RSNN framework must avoid training that relies on long internal spike simulations while preserving the recurrent state that links successive actions.

Of these two challenges, learning is partially addressed by implicit differentiation, which avoids backpropagating through every spike by differentiating the firing-rate state at which the network settles instead of unrolling the trajectory. IDE-FSNN and SPIDE introduced this approach for RSNNs [18,19]. Although highly promising, these methods still obtain the forward rates from explicit spike simulation, leaving a substantial computational burden in every forward pass. In addition, their reported evaluations concern static and event-based image classification, and the methods have not yet been applied to memory-dependent tasks, including continuous motor control. Extending implicit training to an efficient, sequential computation therefore requires both a firing-rate state that can be inferred directly, without spike simulation, and a mechanism that carries that state from one input to the next.

For a model of motor cortex to learn continuous motor control, it must address two further biological challenges, credit routing and local plasticity. Credit routing concerns how teaching information generated elsewhere in the brain is selectively delivered to the cortical neurons that causally contributed to the behavioral outcome. Exact gradient descent would deliver this information through backward pathways that precisely match the forward computation. This requirement, known as the weight-transport problem [20], has no established anatomical mechanism [21,22]. Feedback alignment showed that learning can proceed even when the backward weights are random and unmatched to the forward weights [21]. Fixed random feedback, however, aligns only approximately with the forward weights and performs poorly on difficult visual-recognition tasks [22]. Learning the feedback weights with a local rule that drives them toward the forward weights, as in the Kolen–Pollack algorithm, improves both alignment and performance [22,23]. In recurrent networks, credit must also be routed through the recurrent connections themselves, which exact gradients do by propagating backward through the transposed recurrent weights. Local rules such as RFLO and e-prop deliver credit only through the feedback pathway and do not address the non-local recurrent terms [24,25]. Biological motor learning further requires that teaching information be conveyed to target neurons through anatomically defined neural pathways. In the brain, teaching signals shaped by the cerebellum and basal ganglia reach motor cortex through the thalamus [6,26–29]. Top-down inputs can also evoke regenerative events in the apical dendrites of pyramidal neurons [30]. Such apical dendritic activity can carry teaching-related signals, including reward and error [31]. These long-range pathways therefore provide a plausible substrate for credit routing.

The second challenge, local plasticity, requires that once a teaching signal reaches its target neuron, each synapse determine its own modification using only locally available information. Reward-modulated spike-timing-dependent plasticity (STDP) partly satisfies this requirement by combining a synapse-specific eligibility trace derived from pre- and postsynaptic activity with a global reward signal [32–35]. However, because the reward signal itself is shared across many neurons, it does not directly provide distinct teaching information to individual target neurons. By contrast, backpropagation-inspired learning rules can deliver neuron-specific teaching signals through feedback pathways [21,22,24,25,36]. In these rules, however, the teaching signal usually enters the plasticity rule directly as a continuous-valued variable. It also leaves the activity of the network unchanged, and learning therefore remains detached from the dynamics that generate behavior. Methods in which feedback does act on network activity impose other constraints. Equilibrium propagation uses energy-based dynamics and weak nudging [37]. Its energy function requires symmetric connections. EqSpike implements this approach with spiking neurons connected by bidirectional synapses and inherits this requirement [38]. Generalized equilibrium propagation, a rate-based method, allows some asymmetry but penalizes it and drives the network back toward symmetry [39]. All three have been applied only to static classification tasks, in which each input is held until the network settles [37–39]. FOLLOW and Error Forcing instead inject prescribed continuous error currents into the network [40,41]. A rule in which teaching information arrives as spikes in an asymmetric recurrent network, such that the resulting changes in neuronal activity drive synaptic plasticity, would therefore provide a more direct biological implementation of local plasticity.

Here, we develop RSNNs that address both the computational demands of continuous motor control and the biological constraints governing learning in motor cortex. We first derive a formulation that eliminates explicit spike simulation from implicit differentiation. We then extend it across successive inputs through State-Carry Initialization (SCI), enabling memory-dependent computation during continuous control. Building on this framework, we introduce Feedback-Spike Nudging (FSN), which enables learning under the constraints of credit routing and local synaptic plasticity. We show that RSNNs trained with FSN learn continuous motor control in both supervised and reinforcement-learning settings. Moreover, the FSN plasticity rule is consistent with experimental observations of structural plasticity in mouse motor cortex during learning [4]. We further discuss how the computational components of FSN may correspond to specific circuit mechanisms in the brain. Together, these results establish a novel RSNN framework for motor cortex that integrates the computational requirements of continuous motor control with the biological organization of cortical learning circuits.

## Results

### A current-balance rate equation enables fast implicit training of RSNNs

We first sought to improve the computational efficiency of implicit differentiation before applying it to continuous learning in a model of motor cortex. Existing implicit-differentiation methods still simulate spikes to obtain the firing-rate state in every forward pass [18,19]. We therefore developed **Current-Balance Implicit Training (CBIT)**, which solves the recurrent firing-rate state directly from the mean synaptic input and differentiates this state implicitly. Because the solved rates are used directly for prediction and learning, they must match the absolute firing rates of the spiking network. To this end, we derived a leak-corrected current-balance rate equation for leaky integrate-and-fire (LIF) neurons (**Fig. 1a**; Methods; derivation in **Supplementary Information S1**), which gives the equilibrium firing rate as

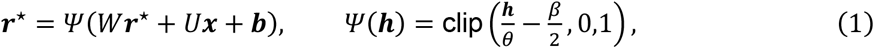

where *r*^*⋆*^ is the fixed-point firing-rate vector and *h* is the mean input current. *W, U*, and *b* are the recurrent weights, input weights, and bias, and *θ* and *β* are the threshold and membrane-leak coefficient. The transfer function *Ψ* maps *h* to firing rate. We solved Eq. (1) for *r*^*⋆*^ by damped fixed-point iteration (Methods). The solved *r*^*⋆*^ closely predicted the empirical firing rates of individual recurrent neurons (**Fig. 1b**). To compare Eq. (1) with alternatives, we embedded its transfer function and five comparator transfer functions in the same recurrent equation and solved each with the same iteration. Across 108 matched recurrent LIF conditions that varied membrane decay, reset mode, recurrent scale, input rate, and random seed, Eq. (1) had a lower mean absolute error (MAE) than every comparator [18,42–44] (Wilcoxon signed-rank test, n = 108 conditions, Holm-corrected p < 1.6 × 10^−4^; **Fig. 1c**). Thus, Eq. (1) accurately estimates the fixed-point firing rate *r*^*⋆*^ of the LIF network for a given input.

**Figure 1.**
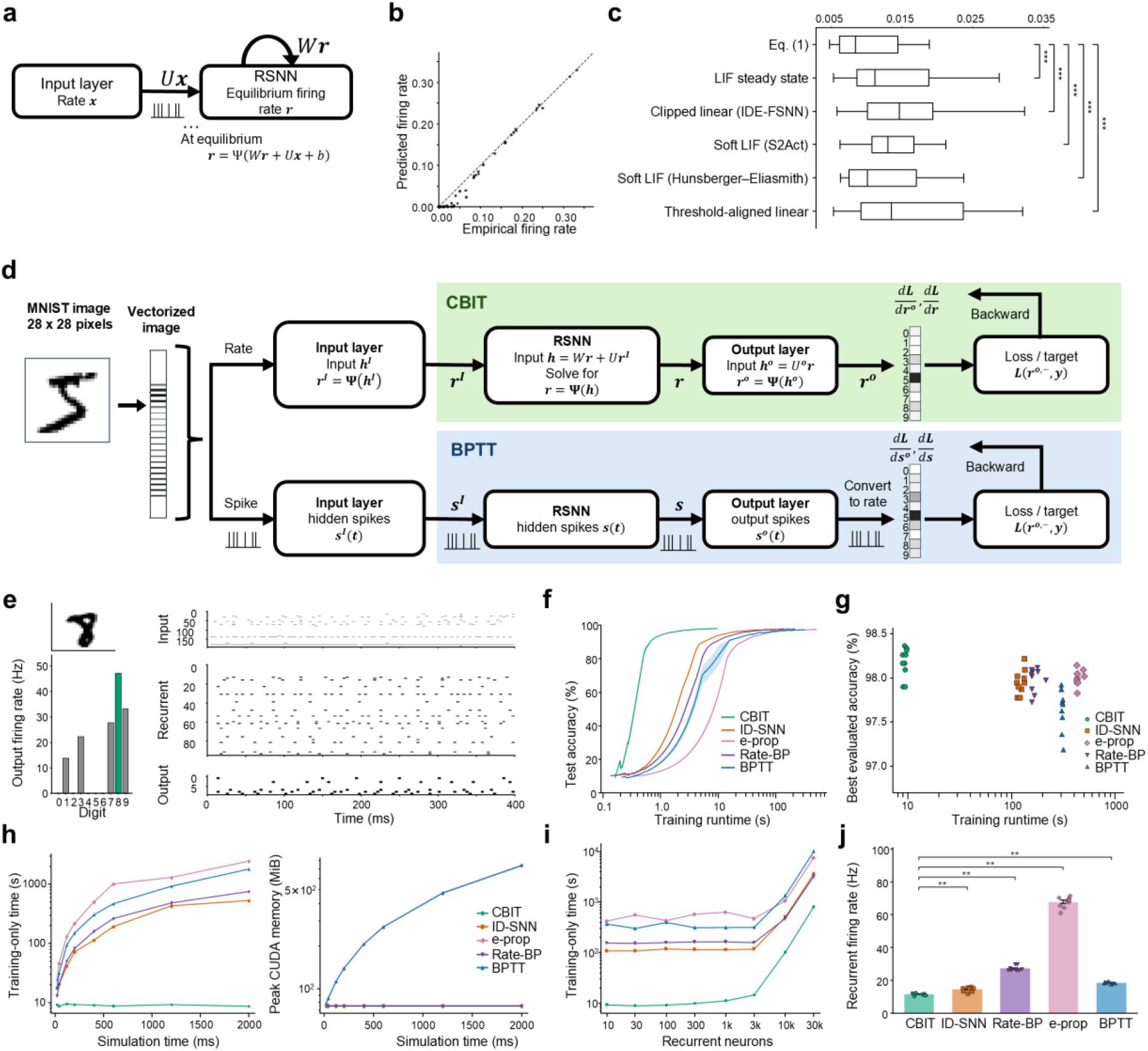
Current-balance equilibrium rates and fast implicit training of RSNNs. **a**, Rate-inference problem solved by Eq. (1). Mean input rates and recurrent synaptic drive determine the equilibrium firing-rate state of the RSNN. **b**, Empirical firing rates versus those predicted by Eq. (1) in one recurrent LIF simulation. Each point is one neuron, and the dashed line marks perfect agreement. **c**, MAE between empirical and predicted firing rates for Eq. (1) and five comparator transfer functions across n = 108 matched recurrent LIF conditions: LIF steady state [42], clipped linear (IDE-FSNN) [18], soft LIF (S2Act) [43], soft LIF (Hunsberger–Eliasmith) [44], and threshold-aligned linear. ***p < 0.001 versus Eq. (1) (Wilcoxon signed-rank test, n = 108, Holm-corrected). **d**, LIF MNIST classifier and its two training paths. BPTT differentiates through the full spike trajectory, whereas CBIT solves the rate state from Eq. (1) and differentiates the recurrent fixed point implicitly. **e**, Spiking evaluation of a CBIT-trained model: input image, input-, recurrent-, and output-layer rasters over the 400-ms stimulus, and the output firing rates used for classification. **f**, Spiking test accuracy versus training-only runtime for CBIT, ID-SNN, Rate-BP, e-prop, and BPTT (n = 10 seeds). **g**, Evaluated spiking accuracy versus training-only runtime (n = 10 seeds). CBIT versus each method: p = 0.0078 (ID-SNN, Rate-BP, and BPTT) and 0.012 (e-prop); Wilcoxon signed-rank test, n = 10, Holm-corrected. **h**, Training-only time (left) and peak CUDA memory (right) versus simulation length (single seed). **i**, Training-only time versus the number of recurrent neurons (single seed). **j**, Recurrent firing rate during spiking evaluation of the final checkpoints (n = 10 seeds). **p = 0.0078 for CBIT versus each method (Wilcoxon signed-rank test, n = 10, Holm-corrected).

We next used Eq. (1) to compute learning gradients. Let *L* denote the task loss and *W*_*ij*_ the recurrent weight from presynaptic neuron *j* to postsynaptic neuron *i* . Training requires *∂L*/*∂W*_*ij*_ for every synapse. BPTT obtains it by storing the entire spike trajectory and backpropagating through every timestep. Implicit differentiation avoids this. Because *r*^*⋆*^ satisfies the fixed-point equation above, differentiating both sides of that equation with respect to *W* gives the sensitivity of *r*^*⋆*^ to each weight in terms of the solved state alone, without any trajectory. Combining this sensitivity with *∂L*/*∂r*^*⋆*^, the derivative of the loss with respect to the solved rates, yields the recurrent weight gradient

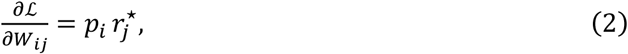

where 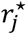 is the presynaptic firing rate and *p*_*i*_ is the postsynaptic credit. The presynaptic factor is available locally at the synapse. The credit *p*_*i*_, by contrast, is obtained from *∂L*/*∂r*^*⋆*^ by solving a linear system that propagates credit backward through the transposed recurrent matrix *W*^*⊤*^ and is therefore not local (derivation in **Supplementary Information S2**).

To evaluate how well CBIT learns, we used it to train a recurrent MNIST classifier [45], with Eq. (1) providing the forward-pass state and implicit differentiation providing the backward pass (**Fig. 1d**). We compared CBIT with four other methods. Spike-forward implicit differentiation (ID-SNN in figures) simulates the spikes to obtain the firing-rate state and uses the same implicit backward pass. It thereby reproduces, within our architecture, the approach shared by the earlier implicit-differentiation SNN methods IDE-FSNN [18], SPIDE [19], and MPIS [46]. The other comparators were rate-based backpropagation (Rate-BP) [47], surrogate-gradient BPTT [15], and e-prop [25]. CBIT reached the highest classification accuracy of the five methods (98.2%; Wilcoxon signed-rank test, n = 10 seeds, Holm-corrected p ≤ 0.012 for every comparison; **Fig. 1e–g**). It was also the fastest and most memory-efficient method at every simulation length tested (**Fig. 1h**). Its cost does not grow with simulation length because it replaces the simulated timesteps by a fixed, much smaller number of solver iterations (ten in this comparison; Methods, **Table 1**). Its speed advantage persisted as the recurrent population grew from 10 to 30,000 neurons, and at 30,000 neurons CBIT trained 4.0-to 12.6-fold faster than the other methods (**Fig. 1i**). In addition, its recurrent neurons fired most sparsely (11.7 Hz versus 14.7–67.8 Hz for the other methods; p = 0.0078 for each comparison; **Fig. 1j**). Thus, CBIT combined the highest accuracy with the highest efficiency in MNIST classification.

**Table 1.** Order of training cost and memory.

| Method | Arithmetic per update | Memory per sample |
|---|---|---|
| BPTT | O(TN<sup>2</sup>) | O(TN) |
| e-prop | O(TN<sup>2</sup>) | O(N<sup>2</sup>) (eligibility-trace form)<br>O(TN) (automatic-differentiation form) |
| Rate-BP | O(TN<sup>2</sup>) | O(N) |
| ID-SNN | O((T + J)N<sup>2</sup>) | O(N) |
| CBIT, CBIT+SCI | O((K + J)N<sup>2</sup>) | O(N) |
| FSN, cFSN | O((T<sub>free</sub> + T<sub>feedback</sub>)N<sup>2</sup>) | O(N) |

### State-Carry Initialization preserves trajectory-dependent memory

Although CBIT achieved high MNIST classification accuracy, MNIST involves only classification of discrete inputs. Extending CBIT to continuous control requires maintaining temporal continuity across successive state estimates. To retain context across successive solves, we introduced **State-Carry Initialization (SCI)**, in which the damped fixed-point iteration of Eq. (1) for each input phase *m* starts from the state solved for the preceding phase *m* − 1 rather than from zero (Methods; **Supplementary Information S3**). Because gradients were not propagated backward across phases, any memory of earlier inputs had to be carried by the state itself.

To verify that the carried state alone can retain such memory, we tested SCI on a sequential version of modular addition, a simple algorithmic task widely used to probe internal computation in neural networks [48,49]. In this task, the network is shown a first digit *x*, then an intervening cue, then a second digit *y*, and must report (*x* + *y*) mod 10 only after the second digit (**Extended Data Fig. 1a**). The first digit must be held in the recurrent state throughout the cue period. CBIT+SCI reached a spiking accuracy of 97.7% across ten seeds. Even the least accurate of the 100 ordered digit pairs reached 87.4% (**Extended Data Fig. 1b**).

The trained networks also showed why SCI is needed. In the space of the first two principal components of recurrent spiking activity, the first- and second-digit trajectories followed paths specific to each digit pair. The states solved with SCI lay at the endpoints of these paths (**Extended Data Fig. 1c**). Across the ten trained models, the accuracy computed from the solved rates (theoretical accuracy) was 98.9% with SCI, compared with 10.0% from zero initialization, 10.2% from random initialization, and 9.8% from a clipped closed-form solution (chance, 10%). In an example network, discarding the carried state also abolished the agreement between solved and empirical output rates after the second digit (*R*^2^ = 0.88 with SCI versus −0.32, −0.20, and −1.01 for zero, random, and closed-form initialization; **Extended Data Fig. 1d**). SCI therefore carried task-relevant history from one rate solve to the next.

**Extended Data Figure 1.**
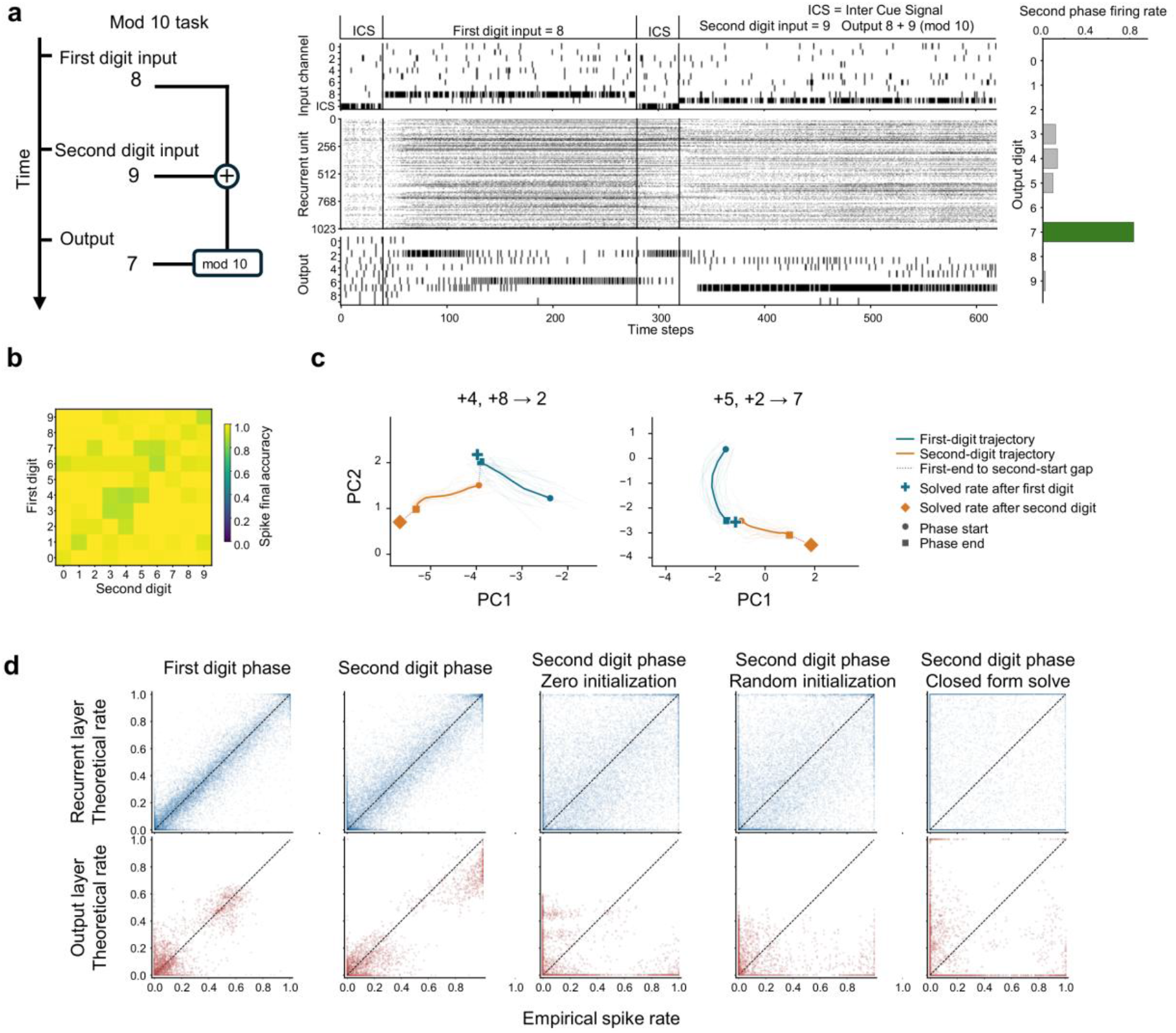
State-Carry Initialization enables memory-dependent modular addition in an RSNN. **a**, Sequential modular-addition task and representative spiking activity. Each trial presents an inter-cue signal (ICS), the first digit *x*, a second ICS, and the second digit *y*. The target, (*x* + *y*) mod 10, is reported after the second digit. The example shows *x* = 8 and *y* = 9, for which the correct output is 7. Rasters show input, recurrent, and output spikes across the trial. The output firing rates at the end of the trial select the correct digit. **b**, Spiking accuracy for each ordered pair of first and second digits, averaged over n = 10 seeds. **c**, Recurrent-population activity for two example digit pairs, projected onto the first two principal components (panel titles give the two inputs and the resulting output). Thin lines show individual spiking trajectories (20 trials per phase). Thick lines show the trial-averaged cumulative-rate trajectories for the first-digit (blue) and second-digit (orange) phases. Circles and squares mark the start and end of each phase, and the dotted segment links the two phases. The blue cross marks the state solved from Eq. (1) after the first digit, and the orange diamond marks the state solved with SCI after the second digit. **d**, Theoretical versus empirical firing rates for the recurrent (top) and output (bottom) populations of an example network. Columns show the first-digit phase, the second-digit phase solved with SCI, and the second-digit phase solved from zero, from a random state (16 restarts pooled), or by a clipped closed-form solution.

### SCI enables CBIT to train recurrent spiking controllers by reinforcement learning

We then tested CBIT+SCI in closed-loop continuous control, where observations, actions, and rewards unfold sequentially and the network must act at every step. We used the MuJoCo locomotion tasks Ant-v5 and Walker2d-v5 [50,51], because they require continuous, high-dimensional motor commands from a simulated body. Earlier versions of these continuous-control benchmarks have been used to evaluate spiking controllers [52,53]. At each environment step, the agent receives a proprioceptive observation (the height and orientation of the torso and the angles and velocities of its joints) and must output a torque for each joint (eight for Ant-v5 and six for Walker2d-v5).

At each step, the rates of a recurrent population of 128 LIF neurons were solved from Eq. (1) with SCI, that is, starting from the state solved at the preceding step. Readout populations decoded the mean of the action distribution and a value estimate from the solved rates. The agent was trained with proximal policy optimization (PPO) [54] with generalized advantage estimation (GAE) [55]. The PPO loss defined on the actions and values was differentiated through the implicit rate state (**Fig. 2a**, top). Performance was always evaluated with the spiking network, which received observations as spike trains and generated actions from its output spikes (**Fig. 2a**, bottom).

**Figure 2.**
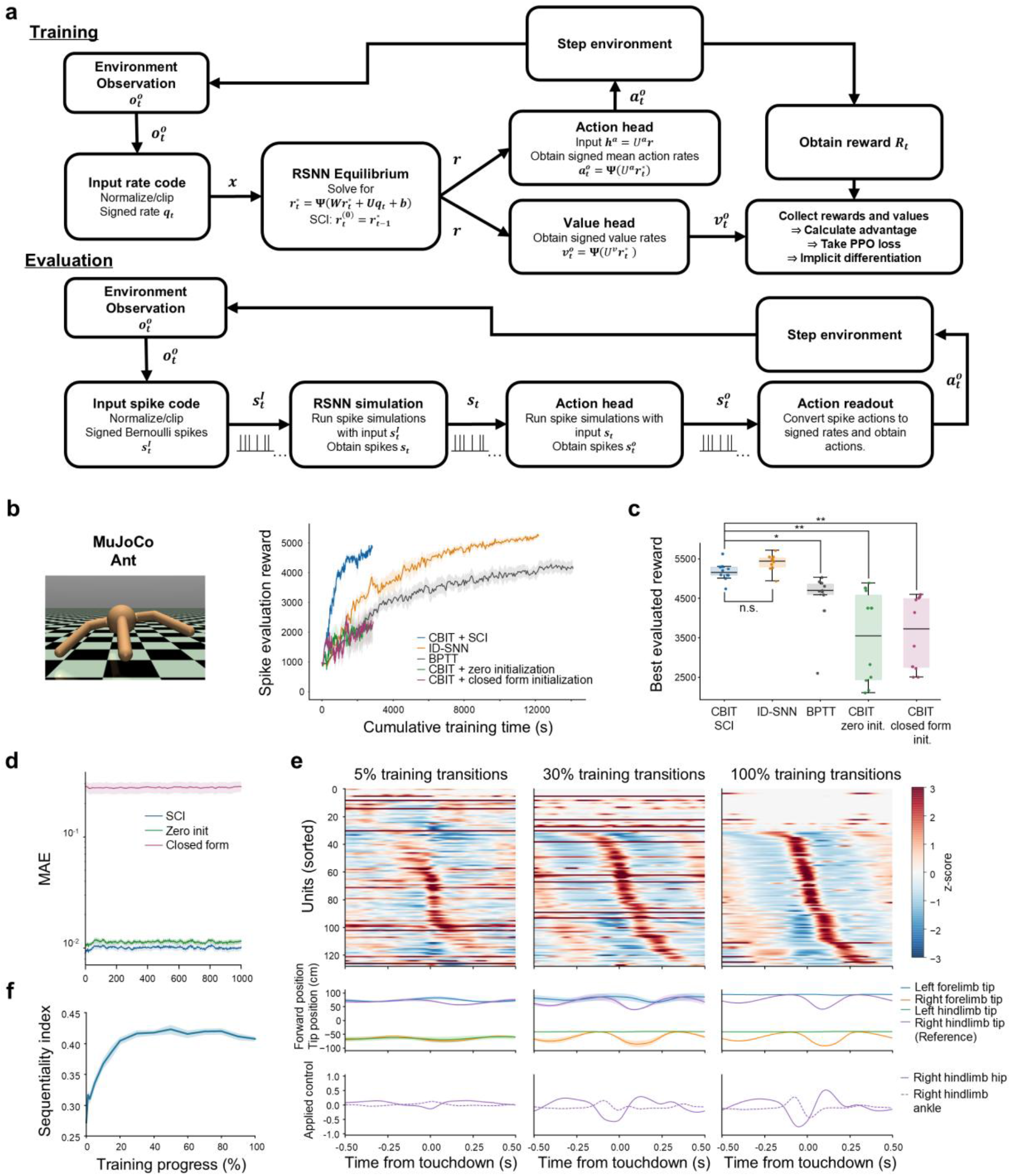
SCI enables CBIT to train recurrent spiking controllers by reinforcement learning. **a**, Training (top) and evaluation (bottom) loops. During training, the observation *o*_*t*_ is normalized and encoded as signed input rates *q*_*t*_. The recurrent LIF state is solved from Eq. (1), starting from the state solved at the previous step (SCI). Action and value heads read out the policy mean *a*_*t*_ and the value estimate *v*_*t*_ from the solved rates 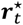. GAE combines rewards and values into advantages. The gradient of the PPO loss is obtained by implicit differentiation through the rate state. During evaluation, the same weights run as a spiking network that receives Bernoulli spike trains and produces actions from its output spikes. **b**, Ant-v5 learning curves. Evaluation reward versus training-only wall-clock time for CBIT+SCI, ID-SNN, BPTT, and the zero- and closed-form-initialization controls (n = 10 seeds). **c**, Ant-v5 best evaluation reward per seed (n = 10). *p = 0.020 (BPTT) and **p = 0.0078 (zero and closed-form initialization) versus CBIT+SCI (Wilcoxon signed-rank test, n = 10, Holm-corrected); n.s., not significant. **d**, Ant-v5 rate tracking: MAE between the recurrent rates solved from Eq. (1) and the recurrent spike rates of the trained spiking controller at each environment step, for solves started from the previous solved state (SCI), from zero, or by the closed-form solution (eight episodes per seed, n = 10 seeds). **e**, Touchdown-aligned recurrent activity and motor output at 5%, 30%, and 100% of training for an example Ant model (right hindlimb touchdown at time zero). Top, z-scored activity of the 128 recurrent neurons (n = 4 held-out episodes), ordered by peak time using four other episodes of the same checkpoint. Middle, forward positions of the four limb tips relative to the torso (mean ± SD, n = 4 episodes). Bottom, mean applied controls for the right hindlimb hip (solid) and ankle (dashed). **f**, Sequentiality index of touchdown-aligned firing rates [56] over training (n = 5, 9, and 10 seeds at 0%, 1%, and later checkpoints).

We compared CBIT+SCI with spike-forward implicit differentiation (ID-SNN) and BPTT in the same architecture. The evaluation reward of CBIT+SCI, the total reward per episode, rose fastest and reached its plateau within a fraction of the training time that ID-SNN and BPTT needed to approach the same level (**Fig. 2b**; **Extended Data Fig. 2a**; **Supplementary Videos 1 and 2**). CBIT+SCI completed training 4.3- and 5.0-fold faster than ID-SNN and BPTT in Ant-v5 and 5.6- and 6.8-fold faster in Walker2d-v5. Its best evaluation reward did not differ significantly from that of ID-SNN and was higher than that of BPTT (Wilcoxon signed-rank test, n = 10 seeds; Holm-corrected p = 0.020 in Ant-v5 and 0.0078 in Walker2d-v5; **Fig. 2c**; **Extended Data Fig. 2b**). In particular, in Walker2d-v5, BPTT reached far lower rewards than either implicit method (mean best reward 1,106, versus 4,798 for CBIT+SCI and 4,728 for ID-SNN). CBIT+SCI was therefore the most efficient method and matched the best performance of any method.

We next asked whether the gradients themselves could contribute to the higher rewards of the two implicit methods. Holding the initial weights, PPO data, and incoming states fixed, we resampled only the input spikes (32 realizations for each of ten initial networks per task). Relative to BPTT, ID-SNN reduced the relative variance of policy gradients by 62.6% in Ant-v5 and 88.8% in Walker2d-v5 (**Extended Data Fig. 2f**). CBIT+SCI uses expected input rates and therefore produced the same gradient in every realization. Both implicit methods had lower variance than BPTT in every seed, for both policy-only and full PPO gradients (Wilcoxon signed-rank test, Holm-corrected p = 0.016 for each comparison). Lower sensitivity to input-spike variability may thus contribute to their better control performance.

To test whether SCI was necessary, we trained two controls in which each solve restarted either from zero or from a closed-form estimate. Both controls lowered the best and final rewards in Ant-v5 and Walker2d-v5 (all Holm-corrected p ≤ 0.016; **Fig. 2b,c**; **Extended Data Fig. 2a,b**). We then replayed the trained CBIT+SCI controllers as spiking networks and compared their recurrent spike rates with the rates solved at each environment step. Rates solved with SCI closely tracked the spike rates throughout 1,000-step episodes. The same solver restarted from zero tracked them less closely, and the closed-form solution failed to track them (**Fig. 2d**; **Extended Data Fig. 2c**). This agreement is expected because the spiking controller, like SCI, carries its state from one step to the next. Together, these comparisons indicate that carrying the preceding state is needed both for accurate rate prediction and for control.

To examine how locomotor coordination developed, we replayed CBIT+SCI checkpoints throughout training and aligned recurrent activity, limb movements, and motor commands to ground contact. At 5% of training, a touchdown-aligned sequence was already present in Ant but was broad and noisy, and the limb tips barely moved relative to the torso. By 30% and 100% of training, the sequence had become sharper and more robust and was accompanied by periodic limb-tip trajectories and by changes in hip and ankle control before and after contact (**Fig. 2e**). Walker2d showed a corresponding development of recurrent activity, foot trajectories, and leg controls (**Extended Data Fig. 2d**). We quantified this sequential firing with the sequentiality index [56]. Between 10% and 50% of training, the index increased from 0.37 to 0.42 in Ant and from 0.30 to 0.35 in Walker2d, and it declined modestly thereafter (**Fig. 2f**; **Extended Data Fig. 2e**). Thus, learning was accompanied by the emergence of robust, contact-aligned neural sequences and coordinated motor output. This temporal organization is qualitatively consistent with motor-cortical recordings during locomotion, in which different neurons are preferentially active at successive phases of the step cycle [57,58].

These experiments established an efficient way to train recurrent LIF controllers for closed-loop continuous control while retaining trajectory-dependent state. The exact CBIT update, however, still requires each neuron to receive the mathematically computed credit *p*, a quantity that no local mechanism in the circuit could supply. We therefore next asked whether that non-local quantity could be replaced by a physical response to feedback spikes within the recurrent circuit.

**Extended Data Figure 2.**
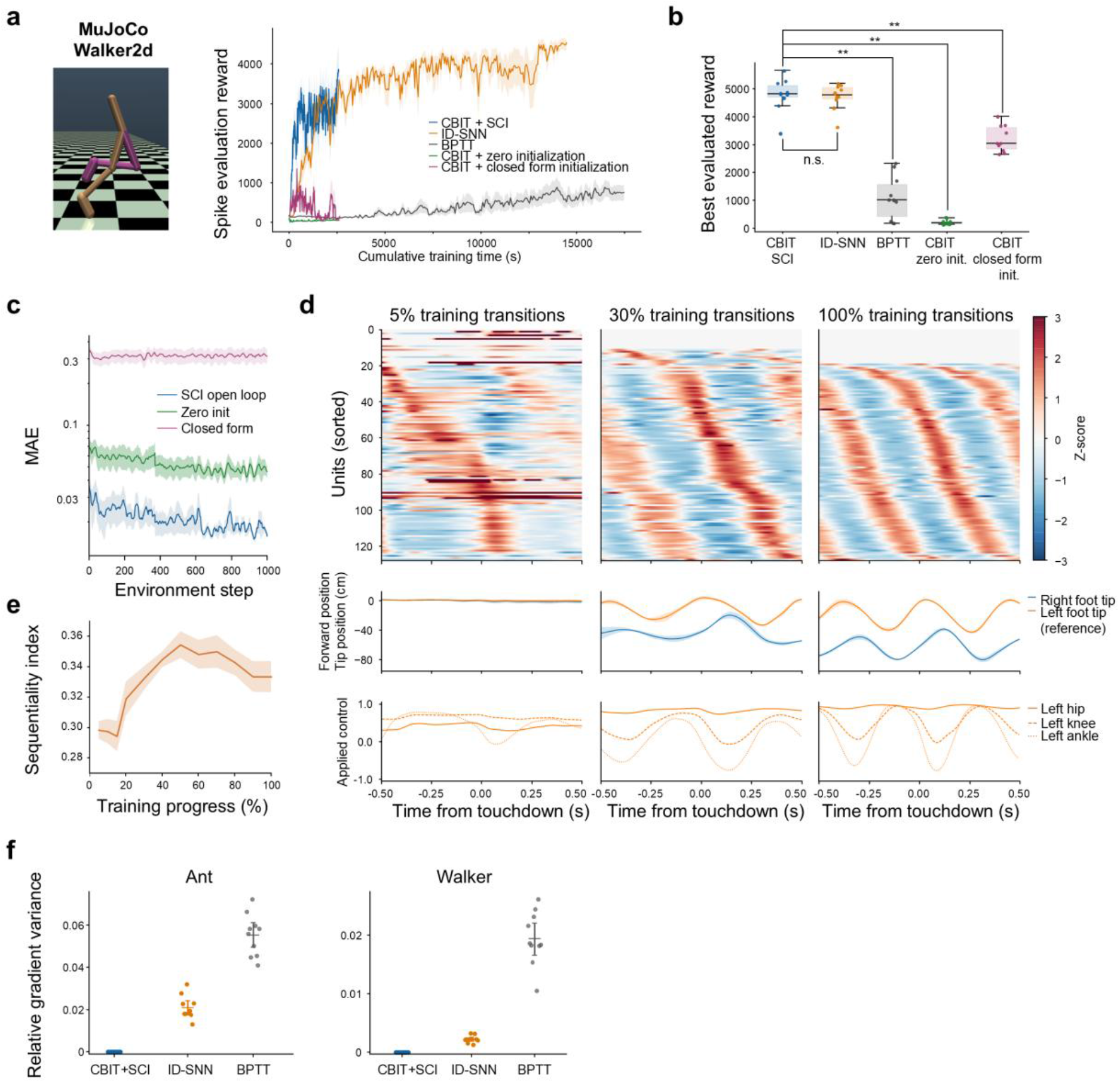
Walker2d control and gradient variability in recurrent spiking policies. **a**, Walker2d-v5 learning curves for the five training methods, as in **Fig. 2b** (n = 10 seeds). **b**, Walker2d-v5 best evaluation reward per seed, as in **Fig. 2c**. **p = 0.0078 versus CBIT+SCI (Wilcoxon signed-rank test, n = 10, Holm-corrected). **c**, Walker2d-v5 rate tracking, as in **Fig. 2d** (n = 10 seeds, of which nine remained at step 1,000). **d**, Touchdown-aligned recurrent activity and motor output at 5%, 30%, and 100% of training for an example Walker2d model (left foot touchdown at time zero), shown as in **Fig. 2e**. Middle, forward positions of the right (blue) and left (orange) feet relative to the torso (mean ± SD, n = 4 held-out episodes). Bottom, mean applied controls for the left hip (solid), knee (dashed), and ankle (dotted). **e**, Walker2d-v5 sequentiality index, as in **Fig. 2f** (n = 7–10 seeds per checkpoint). No seed met the inclusion criteria before 5% of training, when the agents had not yet produced regular step cycles. **f**, Relative variance of policy-only gradients across 32 input-spike realizations in Ant (left) and Walker2d (right), with weights and PPO data held fixed (n = 10 initial networks per task; mean and 95% bootstrap confidence interval). CBIT+SCI uses expected input rates and therefore has zero variance. Both implicit methods had lower variance than BPTT (Wilcoxon signed-rank test, n = 10, Holm-corrected p = 0.016).

### Feedback-Spike Nudging turns the non-local credit into a local, spike-evoked plasticity signal

CBIT trains RSNNs efficiently. As a model of learning in motor cortex, however, it leaves two requirements unmet, both concerning the postsynaptic credit *p*_*i*_ of Eq. (2). The first is credit routing. The error defined at the output must reach the recurrent neurons. Exact gradient descent would send it backward through a copy of the readout weights, for which no biological mechanism is known (the weight-transport problem [21,22]). In cortex, teaching signals instead arrive from other regions, for example through thalamic pathways proposed to convey basal-ganglia and cerebellar teaching signals [26,28]. The second is local plasticity. Each synapse must derive its update from the activity of its own pre- and postsynaptic neurons rather than from a computed variable. In cortical pyramidal neurons, top-down input acts in this way, by changing postsynaptic activity, for example through dendritic spikes that gate synaptic change [30,31,59]. **Feedback-Spike Nudging (FSN)** meets both requirements within the same recurrent spiking circuit (**Fig. 3a**). A learned feedback pathway routes the error as spikes, and the resulting change in each neuron’s firing serves as its postsynaptic credit.

**Figure 3.**
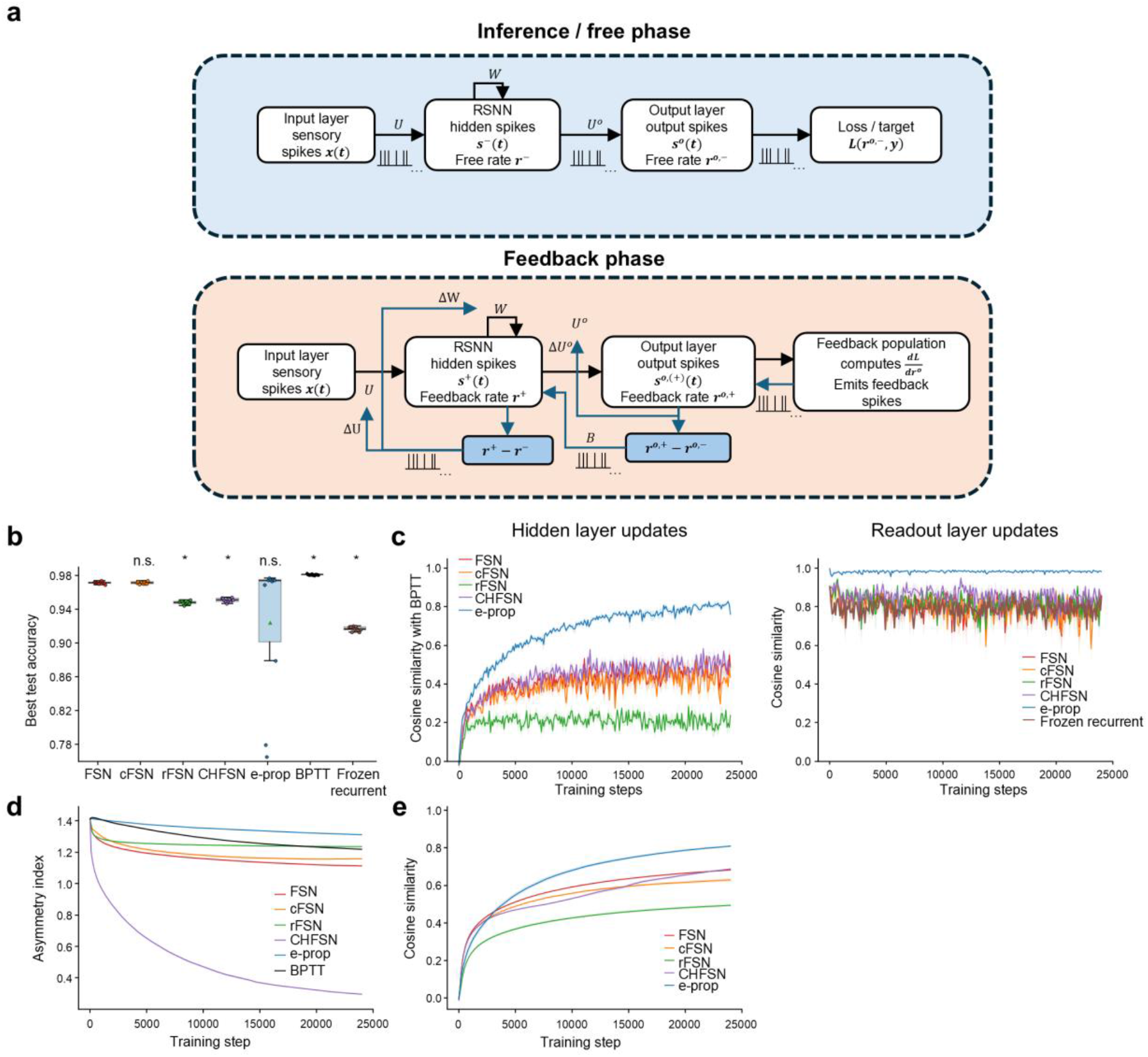
Feedback-Spike Nudging trains recurrent spiking MNIST classifiers. **a**, Two-phase learning mechanism. In the free phase (top), sensory spikes *x*(*t*) drive the recurrent population (recurrent weights *W*) through the input weights *U*. The recurrent population drives the LIF readout through *U*^*o*^. This produces the free rates 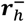 and 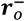 and the loss 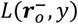. In the feedback phase (bottom), a feedback population converts the output error *∂L*/*∂r*_*o*_ into signed feedback spikes. These spikes perturb the output population and, through the feedback matrix *B*_*out*_, the recurrent population. The resulting rate displacements 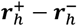 and 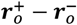 drive the local updates *ΔU, ΔW*, and *ΔU*^*o*^. **b**, Best test accuracy for FSN, cFSN, rFSN, CHFSN, e-prop, BPTT, and the frozen-recurrent control. *p = 0.012 versus FSN (Wilcoxon signed-rank test, n = 10, Holm-corrected). **c**, Left, cosine similarity between the hidden-layer parameter update (input weights, recurrent weights, and recurrent biases) and the negative surrogate-BPTT gradient over training. Right, the same for readout weights and biases (n = 10 seeds). **d**, Asymmetry index of the recurrent matrix *W* over training (lower values indicate greater symmetry; n = 10 seeds). **e**, Cosine similarity between the learned feedback matrix *B*_out_ and the readout *U*^*o*^ over training (n = 10 seeds).

FSN follows from the CBIT gradient. The credit *p*_*i*_ is the sensitivity of the loss to a small current injected into neuron *i*, which the exact gradient computes by propagating the output error backward through the transposed recurrent matrix *W*^*⊤*^. Suppose instead that the error is injected as an actual current into the circuit. The network first runs a free phase, in which sensory input drives the network without feedback, and settles to the free-phase rates *r*^−^. In the feedback phase, signed excitatory and inhibitory feedback spikes encoding the output error enter the output population and, through a feedback pathway *B*, the recurrent population. The network then resettles to the feedback-phase rates *r*^+^ . The resulting displacement *δr* = *r*^+^ − *r*^−^ is the circuit’s own response to the injected error. It coincides with the exact credit −*p* when the recurrent connectivity is symmetric. For an asymmetric *W*, the displacement propagates through *W* itself rather than *W*^*⊤*^ and is therefore a same-circuit approximation to −*p* (Methods; **Supplementary Information S4**). Substituting it into Eq. (2) gives the FSN update

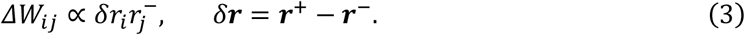

Equation (3) is the product of the presynaptic free-phase rate and the postsynaptic, feedback-evoked change in firing. Both factors are carried by the two neurons that the synapse connects. The rule therefore requires no derivative variable, no copied weight, and no imposed symmetry (derivation in **Supplementary Information S4–S6**). The comparison of a free and a feedback phase is shared with contrastive Hebbian learning [60,61] and equilibrium propagation [37]. However, both methods use a symmetric update that treats the pre- and postsynaptic neurons alike. The FSN product is instead directional, a form first introduced by the generalized recirculation rule for rate-based networks [62]. The presynaptic neuron contributes its free-phase rate, and the postsynaptic neuron contributes its feedback-evoked displacement.

The remaining element is the feedback pathway *B* that carries the spikes. The exact gradient would require *B* to equal the transposed readout *U*^*o⊤*^. The form of Eq. (3) resolves this. Because the update depends only on the firing rates of the two neurons that a synapse connects, and on no variable specific to the synapse, the same local rule can be applied to a readout synapse and to the feedback synapse that connects the same pair of neurons in the opposite direction. With ordinary weight decay, the two weights then converge toward each other without either synapse reading the other’s weight (Kolen–Pollack mirroring [22,23]). FSN therefore denotes the local update of Eq. (3) at every synapse together with this feedback pathway learned through Kolen–Pollack mirroring.

#### Centered FSN (cFSN)

subtracts each neuron’s minibatch mean from both activity factors of the Kolen–Pollack update. The feedback weights then follow the covariance between the free-phase rates of recurrent neurons and the feedback-evoked displacements of output neurons, rather than their raw product. The input, recurrent, and readout updates are unchanged.

### FSN trains recurrent spiking MNIST classifiers

We tested FSN and cFSN on LIF recurrent MNIST classifiers (n = 10 seeds). Two controls kept every other element of FSN and changed one. Random-feedback FSN (rFSN) fixed the feedback pathway at its random initial weights. Contrastive-Hebbian FSN (CHFSN) replaced the FSN product with the full contrastive-Hebbian recurrent update. A frozen-recurrent control (labeled “frozen recurrent” in the figures) fixed the input and recurrent weights and trained only the output weights and bias with the FSN rule. BPTT and e-prop served as non-local comparators under the same spiking evaluation.

We first compared classification accuracy after training. FSN and cFSN reached final spiking accuracies of 97.0% and 96.9%, respectively. FSN exceeded rFSN (94.6%) and CHFSN (93.9%; Wilcoxon signed-rank test, n = 10 seeds, Holm-corrected p = 0.012) and did not differ significantly from cFSN or e-prop (p = 0.74 and 0.85). Only BPTT was more accurate (98.0%; p = 0.012). The best test accuracy reached during training showed the same ordering (**Fig. 3b**). We next assessed learning efficiency from the number of steps needed to reach accuracy thresholds and from the runtime of a fixed training budget. FSN reached 80% and 90% test accuracy in fewer steps than e-prop (p = 0.031 and 0.023), as did cFSN (median 400 and 1,500 steps versus 4,050 and 11,700 for e-prop; **Extended Data Fig. 3c**). They also completed 24,000 training steps in 1,256 s and 1,287 s, approximately 19% faster than BPTT and 2.4-fold faster than e-prop (p = 0.012 for both comparisons; **Extended Data Fig. 3a–c**). FSN and cFSN therefore combined high accuracy with efficient learning.

To determine how much of this performance could be obtained by learning only the readout, we compared FSN with the frozen-recurrent control. The control reached a final accuracy of 91.6%, 5.4 percentage points below FSN (Holm-corrected p = 0.012; **Fig. 3b**). FSN therefore learned more than a readout of fixed recurrent dynamics.

We next asked whether the feedback-induced displacement carries useful gradient information. Although FSN never explicitly computes a gradient, its hidden-layer updates (input weights, recurrent weights, and recurrent biases) aligned with the negative surrogate-BPTT gradient (mean cosine 0.41). This alignment was stronger than with fixed random feedback (0.21; Holm-corrected p = 0.0098; **Fig. 3c**, left). Readout-layer alignment was high for all methods (**Fig. 3c**, right).

The displacement coincides with the negative of the exact credit when the recurrent connectivity is symmetric. We therefore tracked a normalized asymmetry index of *W* over training, with lower values indicating greater symmetry (**Fig. 3d**). The index declined modestly under FSN and cFSN but remained far from zero. CHFSN, by contrast, strongly symmetrized *W* yet reached lower final accuracy than FSN. Despite this asymmetry, the hidden-layer updates of FSN were positively aligned with the negative surrogate-BPTT gradient (**Fig. 3c**). Accurate learning therefore did not require recurrent symmetry.

Finally, we asked whether Kolen–Pollack mirroring aligns the feedback pathway with the readout without weight transport. Kolen–Pollack mirroring increased the alignment between the learned feedback matrix and the readout relative to fixed random feedback (cosine 0.68 for FSN, 0.63 for cFSN, and 0.50 for rFSN; Holm-corrected p = 0.0098 for FSN versus rFSN; **Fig. 3e**).

Together, these results show that the non-local credit exposed by CBIT can be replaced by a spike-driven, same-circuit displacement that supports accurate learning in an asymmetric RSNN.

**Extended Data Figure 3.**
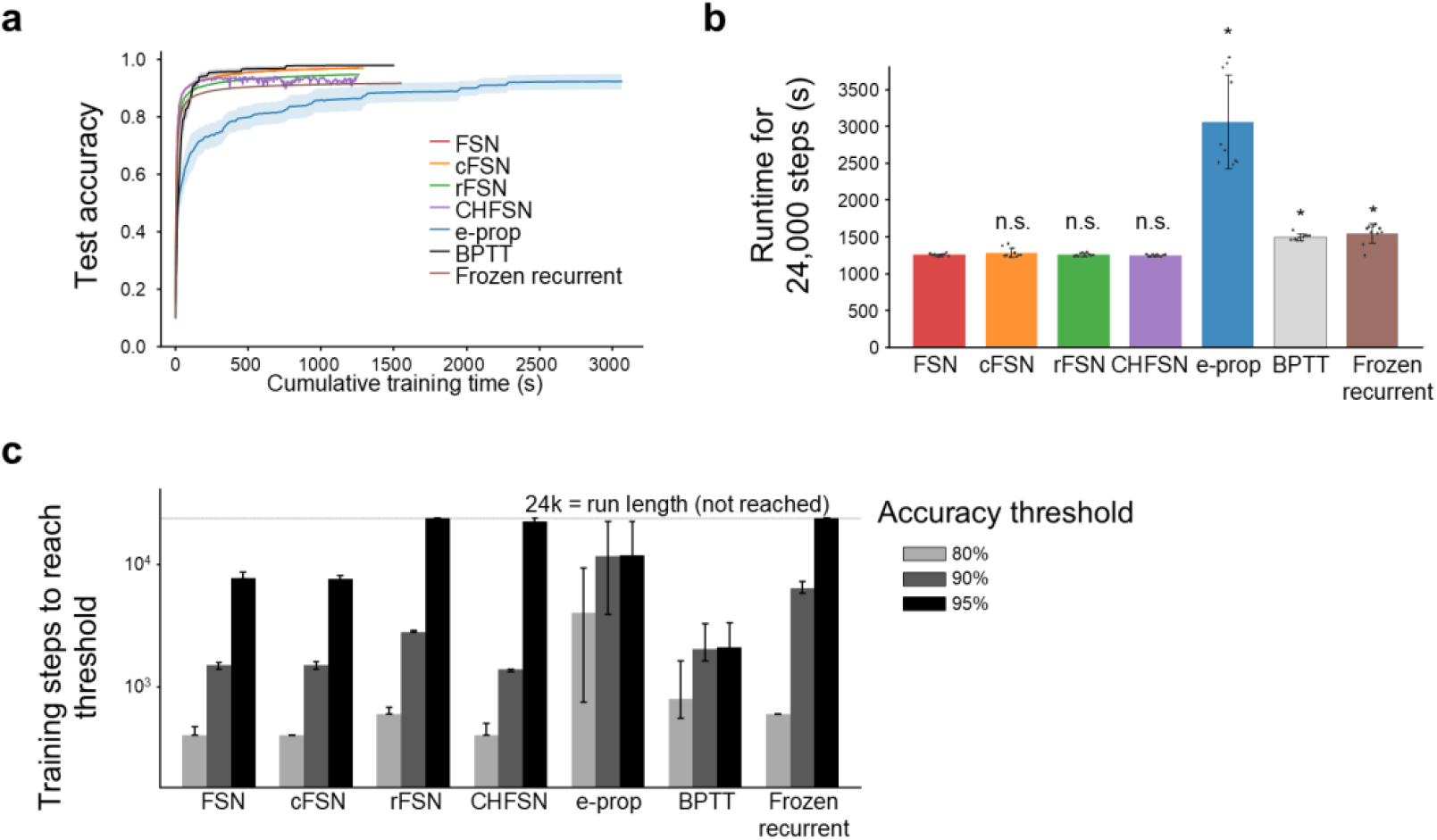
Training performance and efficiency for the MNIST comparison. **a**, Spiking test accuracy versus cumulative training time for the seven methods; each curve ends at the runtime of that method (n = 10 seeds). **b**, Training time for 24,000 updates (mean ± SD, n = 10). *p = 0.012 (BPTT and e-prop) and 0.016 (frozen-recurrent control) versus FSN (Wilcoxon signed-rank test, n = 10, Holm-corrected); n.s., not significant. **c**, Training steps needed to reach 80%, 90%, and 95% test accuracy (median and interquartile range, n = 10). Runs that never reached a threshold were assigned the run length of 24,000 steps (dotted line).

### FSN learns continuous Reacher control through supervised learning

Inspired by models in which cerebellar feedback supports learning and shapes cortical recurrent dynamics [63,64], we next asked whether FSN could support supervised motor learning in a recurrent spiking controller. We used Reacher-v5, in which a two-joint planar arm must bring its fingertip to a target [50,51] (**Fig. 4a**). At each step, the controller receives ten variables (the joint angles and velocities, the target position, and the fingertip-to-target vector) and outputs a torque for each of the two joints. A spiking recurrent student with 256 recurrent LIF neurons learned from a non-spiking teacher network trained with soft actor– critic (SAC) [65]. The student generated each action from its free-phase output. The teacher supplied a target action for the same observation. The difference between the two actions was encoded as signed feedback spikes that perturbed the recurrent state. During training, the final feedback-phase state initialized the next action-generating phase (feedback-state carry). After training, evaluation used the actions and carried recurrent states of the free phase alone, without feedback. In this setting, the teacher-derived action error plays the role of the corrective information that cerebellar feedback supplies in these models. FSN then converts the resulting change in cortical-like recurrent activity into local synaptic updates.

**Figure 4.**
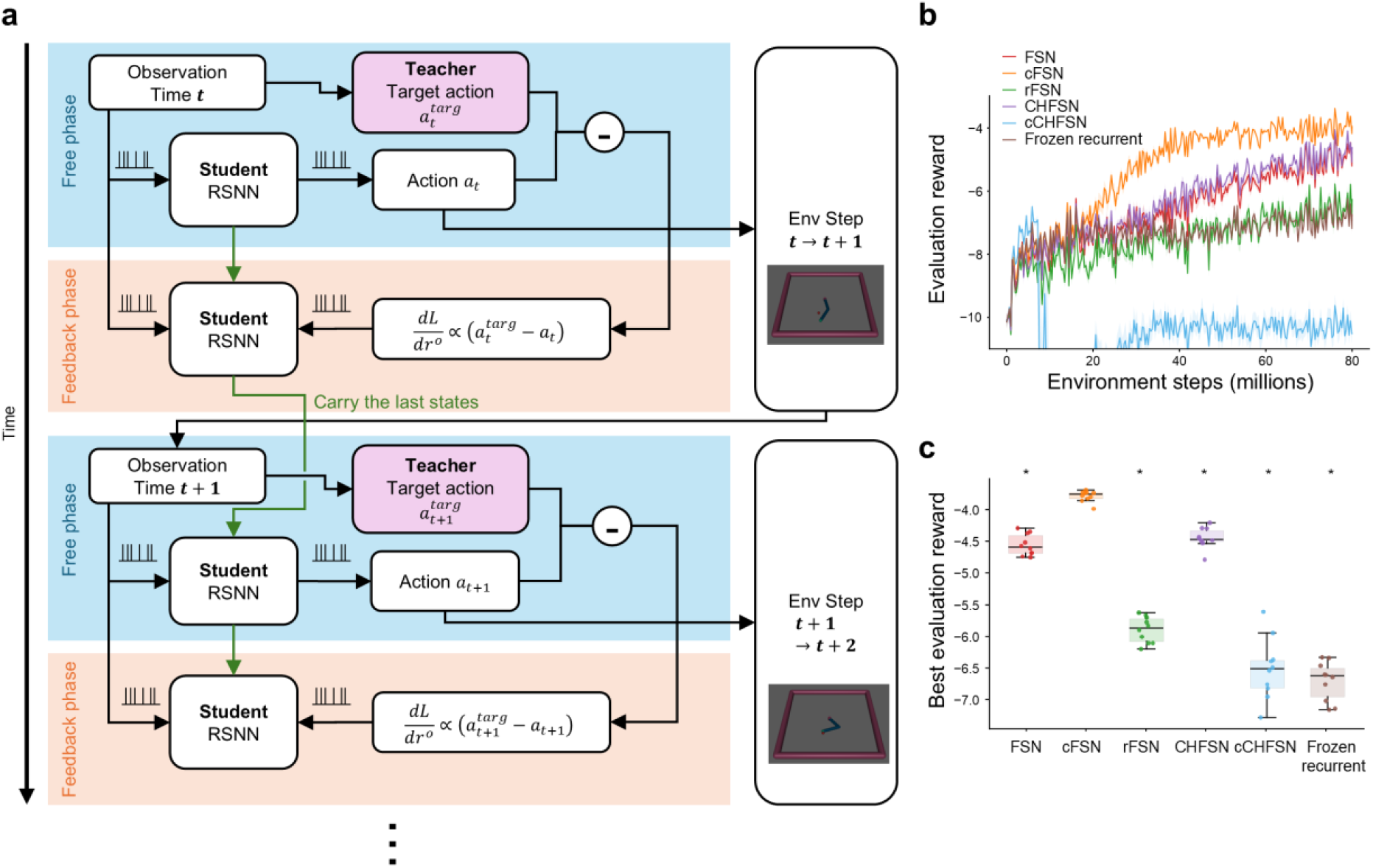
FSN learns continuous Reacher control from a motor teacher. **a**, Supervised training with FSN. At each environment step, the student RSNN runs a free phase, decodes an action from its free-phase output rates, and steps the environment. The pretrained non-spiking teacher supplies only a target action. The action error is converted into signed feedback spikes that perturb the output population and, through the learned feedback pathway *B*_out_, the recurrent population. The perturbed feedback-phase state is carried to the next step. Local updates multiply presynaptic free-phase rates by the postsynaptic displacement between the free and feedback phases. **b**, Reward during training, measured every 20 updates over 64 episodes with teaching feedback (n = 10 seeds). **c**, Reward of the best checkpoint of each run, re-evaluated without teaching feedback on 512 fresh episodes. *p = 0.029 for cFSN versus each other method (Wilcoxon signed-rank test, n = 10, Holm-corrected).

cFSN learned the best policy (**Fig. 4b**; **Supplementary Video 3**), with a mean reward of −3.78 at the best checkpoints evaluated without teaching feedback, compared with −2.88 for the teacher. It exceeded uncentered FSN, CHFSN, rFSN, a frozen-recurrent control, and centered contrastive-Hebbian FSN (cCHFSN; Wilcoxon signed-rank test, n = 10 seeds, all Holm-corrected p = 0.029; **Fig. 4c**). Performance was similar when the same checkpoints were evaluated with feedback (−3.79 versus −3.78). The control comparisons supported contributions from centered Kolen–Pollack mirroring, plasticity of the input and recurrent connections, and the directed update. At the end of training, cFSN also showed the strongest alignment of input and recurrent updates with the gradient (cosine 0.46; **Extended Data Fig. 4a**). Feedback–readout alignment, by contrast, did not predict reward, because FSN and CHFSN aligned more strongly than cFSN but performed worse (**Extended Data Fig. 4b**).

**Extended Data Figure 4.**
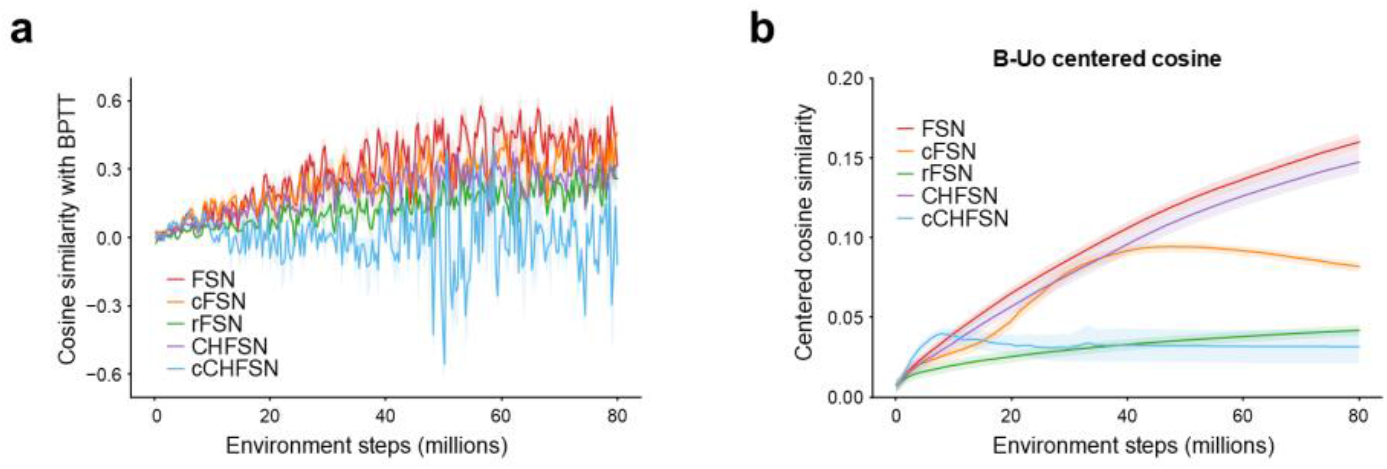
Gradient-alignment diagnostics for FSN on Reacher. **a**, Cosine similarity between each method’s update of the input weights, recurrent weights, and recurrent biases and a surrogate-BPTT reference gradient over training (n = 10 seeds). **b**, Centered cosine similarity between the feedback matrix *B*_out_ and the readout *U*^*o*^ over training (n = 10 seeds).

### Feedback-state-carry cFSN learns Ant locomotion from reward alone

Having tested FSN in supervised motor learning inspired by cerebro-cerebellar interactions, we next asked whether it could also support reinforcement learning. In reinforcement learning, no target action is available. Behavior must instead improve from rewarding outcomes, and the teaching signal therefore has to be inferred from reward. Because this signal depends on sampled actions, subsequent rewards, and estimates of future outcomes, it is noisier and less direct than a supervised target. To test whether FSN could learn from such a signal, we trained a spiking recurrent actor–critic on the Ant-v5 locomotion task. PPO with GAE generated reward-dependent teaching signals at the output, which were encoded as feedback spikes. FSN converted the resulting firing-rate displacements into local synaptic updates while learning from the collected trajectories (**Fig. 5a**). In the brain, basal-ganglia signals relayed through the thalamus to motor cortex could provide a reward-dependent teaching signal analogous to the one defined here by the PPO objective, consistent with the thalamocortical credit-routing architecture proposed by Tanno et al. [28].

**Figure 5.**
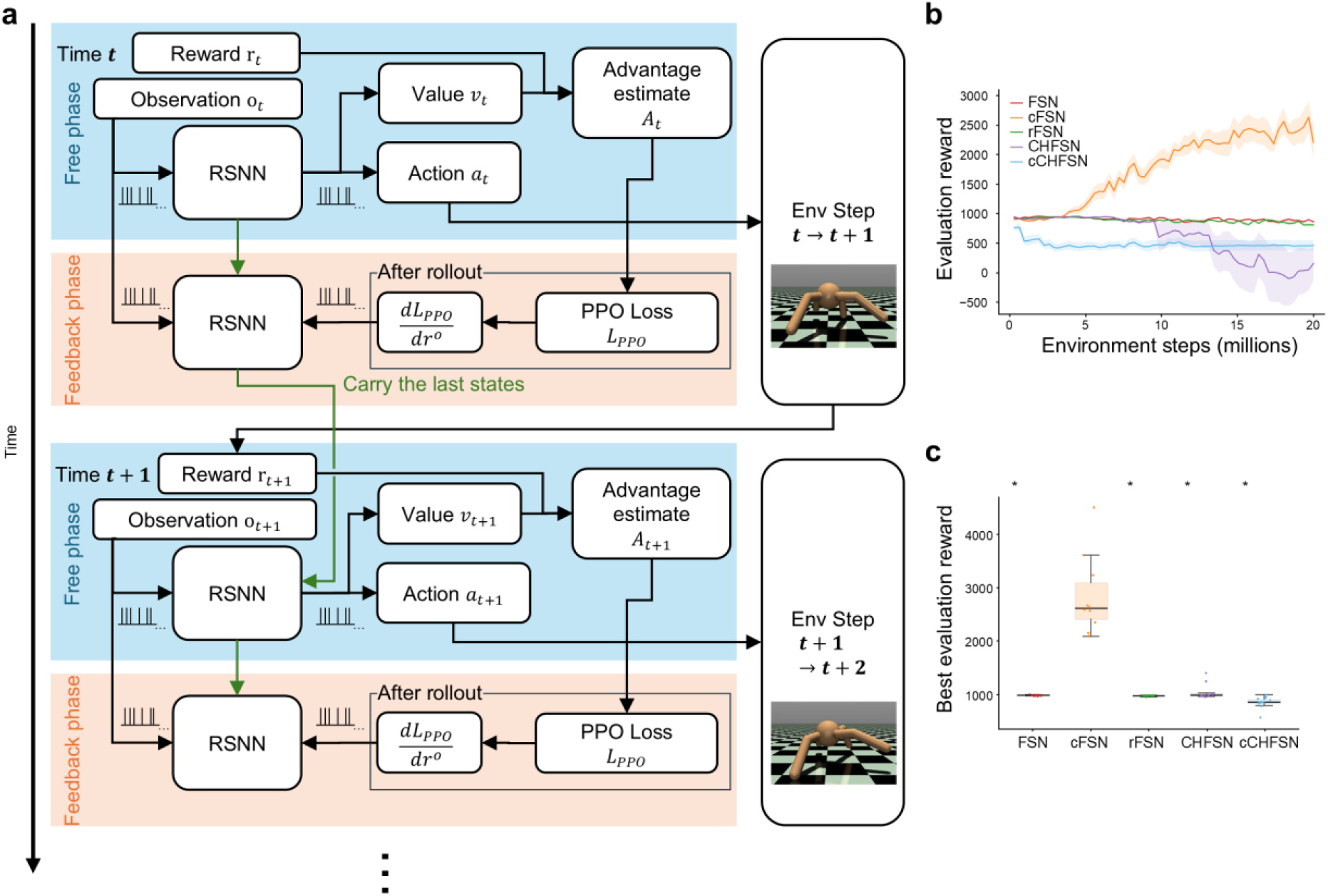
Feedback-state-carry cFSN trains a spiking Ant controller from reward. **a**, Reinforcement learning with FSN. During rollout collection, the actor–critic generates actions and value estimates with fixed weights. After each action, an additional feedback-state window produces the state carried to the next step. After the rollout, GAE computes advantages and value targets. PPO minibatches then replay the stored inputs and initial states. The derivative of the PPO loss with respect to the output rates is encoded as feedback spikes. The resulting rate displacement drives local synaptic updates. The updated weights are used for the next rollout. **b**, Deterministic evaluation reward versus environment steps (n = 10 seeds). **c**, Best evaluation reward per seed. *p = 0.020 for cFSN versus each control (Wilcoxon signed-rank test, n = 10, Holm-corrected).

cFSN was the only local variant that learned sustained forward locomotion (**Fig. 5b**; **Supplementary Video 4**). It reached a mean best evaluation reward of 2,844, whereas all controls remained near the survival-level reward earned by staying upright without moving forward. cFSN exceeded FSN, rFSN, CHFSN, and cCHFSN in best reward in every seed (Wilcoxon signed-rank test, n = 10 seeds, all Holm-corrected p = 0.020; **Fig. 5c**). Only cFSN also accumulated substantial forward reward (1,961, versus 14–127 for the controls). Its total reward remained below that of the CBIT+SCI agent trained with implicit gradients on the same task, mainly because it moved forward more slowly (forward reward 1,961 versus 4,441). Together with the Reacher results, these results indicate that FSN in its centered form supports both supervised and reinforcement learning.

### Longitudinal motor-cortex spine data corroborate the FSN local-product prediction

FSN predicts that synaptic change depends on the product of presynaptic input and a postsynaptic displacement evoked by feedback or reward-related signals. To test this prediction in real synapses, we analyzed the open longitudinal spine-imaging dataset of Hedrick et al. [4], obtained from the apical dendrites of layer 2/3 pyramidal neurons in primary motor cortex during motor learning. For each tracked spine, the dataset contains spine-local glutamatergic input measured with the glutamate sensor iGluSnFR3 [66], calcium activity of the parent dendritic shaft measured with the calcium sensor RCaMP2 [67], and the spine area at each imaging session. We used the glutamate signal as a proxy for presynaptic activity, the shaft calcium signal as a proxy for postsynaptic activity [4], and spine area as a structural proxy for synaptic strength [68].

To map the learning rule onto these signals, we aligned them to reward. In the model, the free phase reflects activity before error or reward feedback arrives, and the feedback phase reflects activity after it arrives. We therefore treated the period after reward delivery as the analog of the feedback phase. For each spine and reward, we measured the event rates of glutamate transients and shaft calcium transients in 2.5-s windows before and after reward. Let the standardized pre-reward rates *a* (presynaptic) and *x* (postsynaptic) stand for free-phase activity, and the post-reward rates *b* and *y* stand for feedback-phase activity. The **FSN product** is then *a*(*y* − *x*), the pre-reward glutamatergic input multiplied by the reward-aligned change in shaft calcium event rate (**Fig. 6a**). It corresponds to the presynaptic free-phase rate multiplied by the postsynaptic feedback-evoked displacement in Eq. (3).

**Figure 6.**
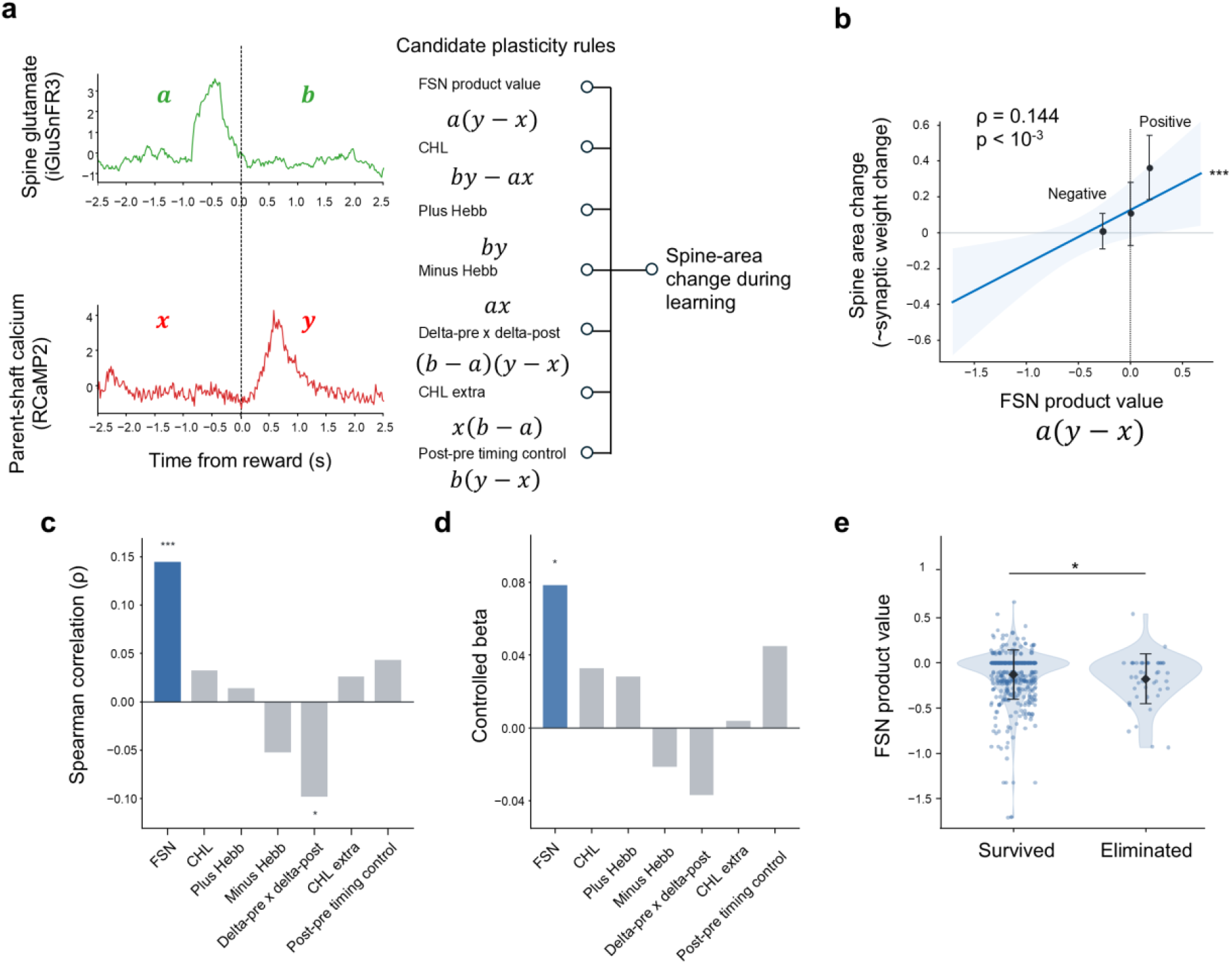
Biological spine-data validation of the reward-modulated local-product rule. **a**, Example spine. Spine-local iGluSnFR3 and parent-shaft RCaMP2 traces aligned to reward delivery (dashed line, *t* = 0). Labels *a* and *b* denote the standardized pre- and post-reward iGluSnFR3 event rates, and *x* and *y* the corresponding RCaMP2 event rates. These rates define the candidate plasticity rules, including the FSN product *a*(*y* − *x*). **b**, Future spine-area change versus the FSN product for n = 517 surviving spine transitions from seven mice. Blue line and shading, linear fit and its 95% confidence band (mouse-clustered covariance); black points and error bars, mean area change and its 95% interval (10,000 mouse-cluster bootstrap resamples) for negative, zero, and positive products, placed at the median product of each group. Spearman ρ = 0.14, ***p = 9.9 × 10^−4^. **c**, Spearman correlation between each candidate rule and future spine-area change (n = 517 transitions pooled from seven mice; FSN product in blue). Rule labels correspond to the formulas in **a**; CHL, contrastive Hebbian learning. *p < 0.05, ***p < 0.001 (p values without correction for multiple comparisons). **d**, Linear-model coefficient of each rule per standard deviation, after adjustment for current log spine area, nearby coactivity within 10 μm, movement-relatedness, and learning phase (n = 517 transitions from seven mice). *p < 0.05 (mouse-clustered Wald t test, df = 6, without correction for multiple comparisons). **e**, FSN products for transitions in which the spine survived to the next session (n = 517) or was eliminated (n = 46), from seven mice. Points, individual transitions; violins, distributions; diamonds and error bars, mean ± SD. *p = 0.028 (one-sided Wilcoxon rank-sum test for higher products in surviving spines).

We first asked whether the FSN product predicted the change in spine area at the next imaging session, which served as a proxy for the change in synaptic strength. Among 517 surviving spine transitions from seven mice, the product was positively associated with future spine-area change (Spearman ρ = 0.14, p = 9.9 × 10^−4^; **Fig. 6b**). To test whether this relationship reflected the specific structure of the rule rather than a generic Hebbian effect, we compared the FSN product with six Hebbian-like and contrastive rules built from the same event rates (**Fig. 6a**). Of the seven rules, the FSN product had the largest positive correlation with future spine-area change (**Fig. 6c**). The product of the glutamate and calcium rate changes (delta-pre × delta-post) was negatively associated (ρ = −0.098, p = 0.026), and no other rule, including contrastive Hebbian learning (CHL; ρ = 0.032, p = 0.46), was associated with the outcome.

We then asked whether the FSN product predicted future spine-area change beyond known covariates. Linear models of the 517 transitions included current spine area, the coactivity of nearby spines within 10 μm, movement-relatedness, and learning phase, with standard errors clustered by mouse. The coefficient of the FSN product remained positive (0.078 per standard deviation; 95% CI, 0.016 to 0.14; p = 0.022; **Fig. 6d**), whereas the CHL coefficient did not differ from zero (0.033, p = 0.39). The relationship therefore held beyond baseline spine size, local coactivity, movement-relatedness, and learning phase.

The same relationship extended to spine survival. Across 563 spine transitions, the FSN product was higher when the spine survived to the next session (n = 517; mean −0.13) than when it was eliminated (n = 46; mean −0.18; one-sided Wilcoxon rank-sum test, p = 0.028; **Fig. 6e**).

Together, these analyses provide independent biological support for the FSN update rule. Presynaptic activity before reward multiplied by the reward-aligned change in postsynaptic activity predicted later spine growth, whereas the Hebbian-like alternatives built from the same signals did not.

## Discussion

We set out to build a model of motor cortex that satisfies three conditions, namely that it is a recurrent network of spiking units, that its plasticity is local, and that plasticity is triggered by the change in postsynaptic activity that a learning signal evokes. To this end, we developed three new methods, CBIT, SCI, and FSN. CBIT enables fast training of RSNNs. Combining CBIT with SCI further allows RSNNs to learn motor control efficiently. Because an equilibrium firing-rate state does not by itself retain how it was reached, SCI starts each equilibrium from the preceding one, so that the network carries its internal state from one moment of behavior to the next, as motor cortex does. By further introducing a local learning rule mediated by changes in postsynaptic activity, we obtained an RSNN that learns through FSN. This network learned continuous motor control, a central function of motor cortex, through both supervised and reinforcement learning. Furthermore, in synaptic input and postsynaptic activity imaged in motor cortex during motor learning, the local learning rule used in FSN predicted which spines would subsequently enlarge and survive. The network thus satisfies physiological and anatomical constraints at the synaptic and cellular levels while performing one of the most fundamental functions of motor cortex, continuous motor control. At the macroscopic anatomical level, the model could also be mapped onto cerebro-cerebellar and cortico-basal ganglia circuits. The RSNN trained with FSN therefore provides an integrated and biologically plausible model of motor cortex.

Because each factor of the FSN update is an activity physically present at the synapse, the rule can be compared directly with cortical physiology. In FSN, each synapse changes in proportion to its presynaptic activity multiplied by the change in postsynaptic firing that feedback spikes evoke (**Extended Data Fig. 5a**). The rule therefore needs no continuous teaching variable as a plasticity factor (RFLO and e-prop [24,25]), no surrogate spike derivative (surrogate-gradient learning [15]), and no imposed recurrent symmetry (equilibrium propagation [37]). At the cellular level, it depends on two operations, each with an experimental counterpart. First, a teaching signal must act by changing the activity of the postsynaptic neuron, and this change must drive plasticity. Top-down feedback to cortical pyramidal neurons evokes calcium events in their apical dendrites [30], which drive burst firing at the soma [69] and induce lasting potentiation of the synapses active at the time [59]. Such instructive input thus promotes plasticity through the activity it evokes in the postsynaptic neuron and changes that neuron’s firing, which is the quantity that FSN uses as credit. Because this instructive input can differ between neurons and even between dendritic branches [31,59], it can also deliver neuron-specific credit directly, rather than leaving each synapse to infer its credit from a global neuromodulatory signal [34,35]. Second, a trace of recent presynaptic activity must persist until the feedback-evoked response arrives. Behavioral time-scale synaptic plasticity (BTSP) has this structure. In adult mouse V1, presynaptic inputs active in the seconds preceding a plateau-like instructive depolarization are potentiated [70], much as FSN pairs the free-phase presynaptic rate with the subsequent feedback-evoked change in postsynaptic firing. Eligibility traces converted into lasting plasticity by delayed signals have also been demonstrated in cortex [71]. Together, these two operations give the presynaptic and postsynaptic neurons distinct roles, the former supplying eligible input and the latter the instructive response. This asymmetry is also what distinguishes the FSN product from the contrastive-Hebbian update, which treats the two neurons alike [61]. The same directional form was proposed earlier as the generalized recirculation rule for rate-based networks [62]. In that work, however, the directional rule learned less reliably than its symmetric variant, which is equivalent to contrastive Hebbian learning and relies on symmetric weights. FSN reverses this result. With feedback weights learned by Kolen–Pollack mirroring and an asymmetric recurrent matrix, the directional update learned continuous motor control, and it also outperformed the contrastive-Hebbian variant (**Fig. 3b–d**). Consistent with the directional form, in the motor-cortex spine data the FSN product predicted later spine enlargement, whereas the contrastive-Hebbian product did not (**Fig. 6c,d**).

At the circuit level, FSN requires two conditions. First, the teaching signal must be routed from other brain regions to motor cortex. Such signals could be supplied by the basal ganglia and cerebellum, which have been proposed to support reinforcement and supervised learning, respectively [72]. Their outputs reach motor cortex and secondary motor cortex (M2) through the motor thalamus [26,28,29,73]. Axons from M2 and the ventromedial (VM) nucleus of the thalamus terminate in layer 1 of motor cortex [28,74,75], on the apical dendrites of intratelencephalic (IT) and pyramidal-tract (PT) neurons (**Extended Data Fig. 5b**). The reward-derived teaching signal in our Ant task is weighted by advantages built from temporal-difference errors [54,55], paralleling computations ascribed to the basal ganglia. The action error of our Reacher task, in turn, parallels the corrective feedback used in a cerebro-cerebellar learning model [63]. Second, within motor cortex, credit must be assigned to each neuron in proportion to its contribution to the motor output. FSN learns the feedback pathway with the local Kolen–Pollack mirroring rule [22,23]. This rule applies the same local plasticity to each readout synapse and to the feedback synapse connecting the same two neurons in the opposite direction, so that the two weights converge. This suggests that Kolen–Pollack mirroring could operate at the corresponding synapses in motor cortex, with equivalent plasticity at the teaching synapses onto a neuron and at that neuron’s own output synapses. Such pairings could involve the input and output synapses of IT neurons or those of PT neurons (**Extended Data Fig. 5b**). The latter pairing is consistent with the selective strengthening of thalamocortical inputs onto the corticospinal neurons engaged in a learned task [76] and with pathway-specific presynaptic supervision of motor-cortical spine dynamics [3]. The requirements of FSN therefore lead to experimentally testable hypotheses about plasticity in motor cortex.

From a computational standpoint, learning through the equilibrium firing-rate state rather than through individual spikes suggests two ways to use event-driven neuromorphic hardware [77,78]. In the first, the gradient is computed on conventional digital hardware, either before deployment or alongside a deployed network. CBIT solves and differentiates the firing-rate state at which the spiking dynamics settle, and its training cost is therefore largely independent of the length of the spike simulation. The resulting implicit gradients are also less sensitive to input-spike sampling than BPTT gradients. SCI extends this implicit training, previously applied only to static inputs [18,19], to memory-dependent and closed-loop tasks. These properties allowed spiking controllers to learn difficult reinforcement-learning tasks, including Walker2d locomotion, on which BPTT reached far lower rewards. Because every evaluation in this study ran the trained weights as a spiking network, a CBIT-trained network can be transferred unchanged to event-driven hardware, without retraining on the device. The same gradient could support online learning. An implicit gradient obtained by a few iterations costs about an order of magnitude less than integrating the spikes over one decision (Methods, **Table 1**). A programmable digital processor could therefore compute this gradient alongside a running spiking controller and update the weights without interrupting control. CBIT-trained networks also fired the most sparsely of all methods compared in MNIST classification, making them even better suited to energy-efficient neuromorphic implementation. In the second, the spiking network learns on the hardware itself. FSN learns from forward dynamics and local products. Its teaching signal can also be delivered while the controller is acting, a form suited to on-chip and online learning in neuromorphic and robotic systems [77]. Moreover, because the FSN update uses only the activity of the two neurons that a synapse connects, it is not restricted to a single recurrent population. Feedback spikes routed to additional layers would apply the same local rule at every synapse, opening a path toward large multilayer spiking models of cortical circuits. Together, our results indicate that credit computed at the equilibrium firing-rate state can both train RSNNs efficiently and, when supplied physically by feedback spikes, act as a local plasticity rule consistent with learning-related changes at motor-cortex synapses.

**Extended Data Figure 5.**
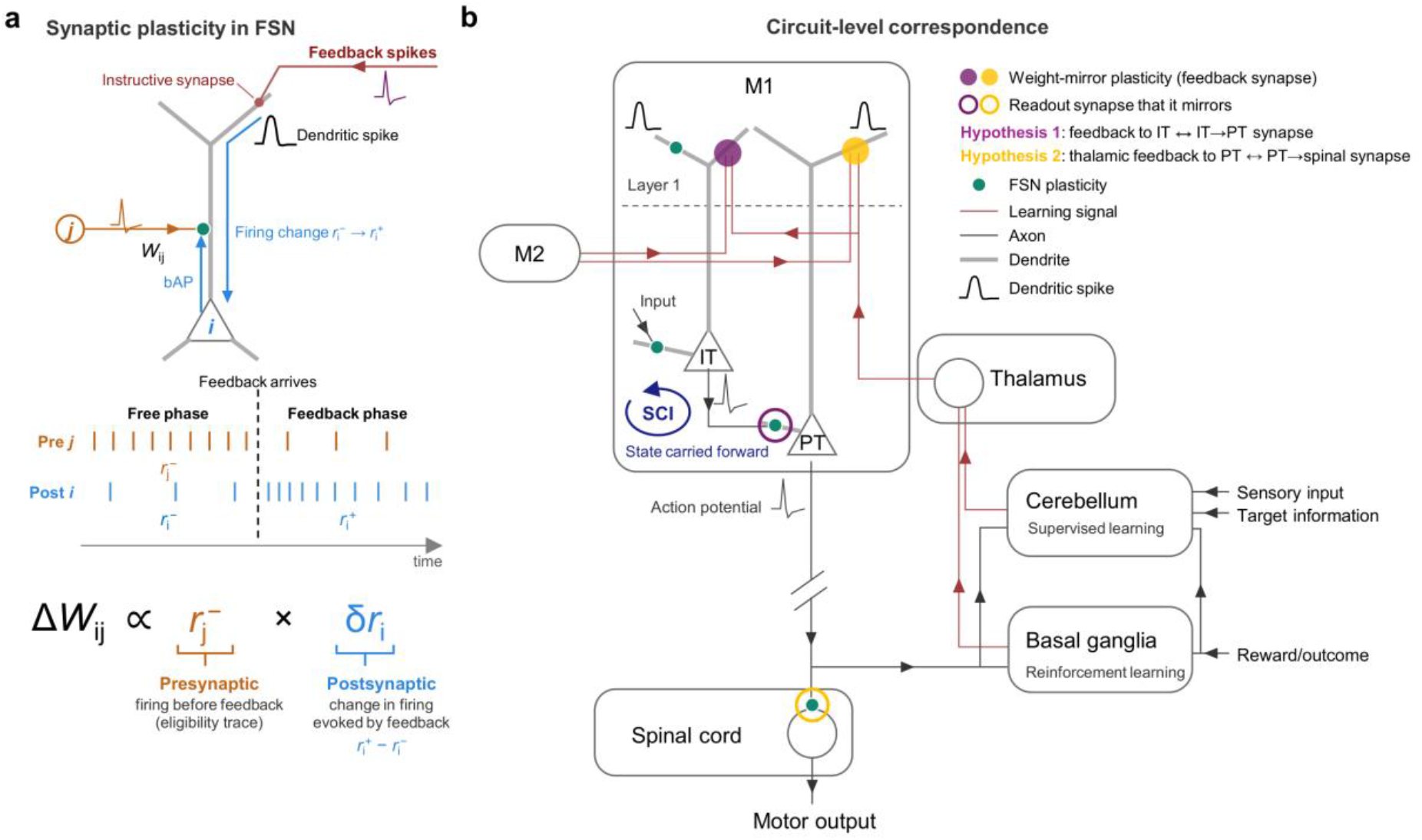
Proposed correspondence between the RSNN trained with FSN and motor-cortical circuits. **a**, Neuron-level correspondence. 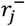 denotes the firing rate of presynaptic neuron *j* before the arrival of feedback. 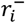 and 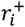 denote the firing rates of postsynaptic neuron *i* before and after feedback, respectively. The feedback signal provides synaptic input to layer 1, and the resulting postsynaptic depolarization, potentially involving a dendritic spike, promotes neuronal firing. The feedback-induced change in firing rate is given by 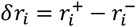. This signal is conveyed to the synapse between neurons *i* and *j* through backpropagating action potentials, thereby promoting synaptic plasticity. **b**, Circuit-level correspondence. Primary motor cortex (M1) has two major classes of excitatory neurons, intratelencephalic (IT) neurons and pyramidal-tract (PT) neurons. Learning signals (red) reach the apical dendrites in layer 1 from the motor thalamus and/or secondary motor cortex (M2). They induce postsynaptic depolarization, potentially involving dendritic spikes. The thalamus relays signals from the cerebellum (supervised learning) and basal ganglia (reinforcement learning) [72]. The sites at which plasticity occurs through the mechanism described in panel **a** are indicated by green dots. We propose that Kolen–Pollack mirroring occurs either between the input and output synapses of IT neurons (hypothesis 1, yellow) or between the input and output synapses of PT neurons (hypothesis 2, purple). SCI corresponds to the recurrent M1 state carried from one moment to the next. Collaterals of PT axons send a copy of the motor command to the cerebellum and basal ganglia.

## Methods

### Discrete-time LIF dynamics and firing-rate readout

For membrane voltage *v*_*t*_, synaptic current *h*_*t*_, threshold *θ*, membrane decay *α*, and spike *s*_*t*_ *∈* {0,1}, the pre-reset voltage and spike were

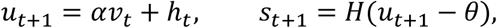

where *H* is the Heaviside function and the per-step leak amount is *β* = 1 − *α*. Subtractive-reset neurons used

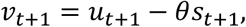

whereas zero-reset neurons used

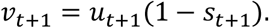

For a simulation of *T* internal timesteps with burn-in *T*_burn_, the firing rate of neuron *i* was

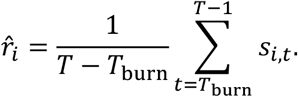

### Current-balance rate equation

Consider a subtractive-reset LIF neuron *i* driven by a stationary input current with mean *h*_*i*_. When the voltage update is averaged over a long window, the time average of the spike train *s*_*t,i*_ converges to the firing rate *r*_*i*_, the time average of the membrane voltage *v*_*t,i*_ converges to its mean *⟨v*_*i*_*⟩*, and the average change of the voltage from one step to the next vanishes at stationarity. The mean input current must therefore balance the current removed by the spike resets and by the leak, which gives the current-balance identity (derivation in **Supplementary Information S1**)

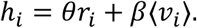

Approximating the mean subthreshold voltage by the midpoint of the integration interval, *⟨v*_*i*_*⟩ ≃ θ*/2, gives the leak-corrected current-to-rate map of Eq. (1) (transfer function *Ψ*)

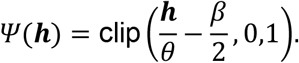

*Ψ* is applied elementwise to the current vector *h*. For recurrent weights *W*, input weights *U*, input-rate vector *x*, and bias *b*, the theoretical recurrent state *r*^*⋆*^ was the fixed point

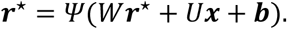

The fixed point was obtained by damped fixed-point iteration with damping factor *η*,

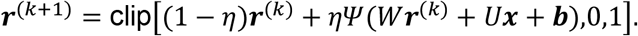

During training, the iteration ran for a fixed number of steps.

### Firing-rate benchmark

The benchmark crossed three random seeds, three membrane decay factors (0.85, 0.90, and 0.95), two reset modes (zero and subtractive), three recurrent-weight scales (0, 0.25, and 0.55), and two input rates (0.05 and 0.15 spikes per step), giving 108 matched conditions. Each condition contained 64 recurrent LIF neurons with threshold 1 and 16 Bernoulli input neurons. Recurrent and input weights were Gaussian, with variance inversely proportional to the number of presynaptic neurons, and self-connections were removed. Empirical rates were measured over the last 6,500 timesteps of an 8,000-step spiking simulation. Equation (1) was compared with five transfer functions embedded in the same recurrent equation, named by their functional form with the source in parentheses: LIF steady state [42], clipped linear (IDE-FSNN) [18], soft LIF (S2Act) [43], soft LIF (Hunsberger–Eliasmith) [44], and threshold-aligned linear, a control whose linear onset is aligned to the threshold current. Each fixed point was solved from zero by damped iteration. The final fixed point was compared with the empirical rates. Benchmark settings are listed in **Supplementary Information S9**.

### Current-Balance Implicit Training

CBIT trains the recurrent rate state of Eq. (1) without differentiating through all internal spike timesteps. The forward loss was evaluated from the solved rate state. Let

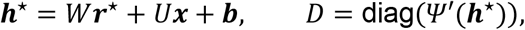

where *D* is the gain of the transfer function, *θ*^−1^ on its linear branch and zero where *Ψ* is clipped. Differentiating the fixed-point condition *r*^*⋆*^ = *Ψ*(*h*^*⋆*^) at fixed input gives

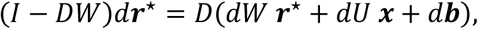

which shows that a weight change alters the solved state through the recurrent loop and that the sensitivity of the state to the weights involves the inverse of *I* − *DW*. Let *λ* be the derivative of the loss ℒ with respect to the solved rates *r*^*⋆*^. Substituting the solved *dr*^*⋆*^ into the loss differential *d*ℒ = *λ*^*⊤*^*dr*^*⋆*^ gives

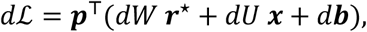

where the current-level credit

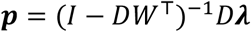

is the loss derivative propagated backward through the transposed recurrent matrix *W*^*⊤*^ and scaled by the gain. Reading off the coefficient of each weight change yields the local outer-product gradients

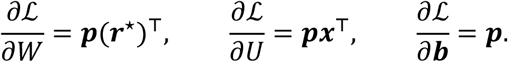

Each synaptic gradient thus combines the postsynaptic current-level credit with the presynaptic activity. Computing *p* is the non-local step, because the inverse matrix above propagates the error through *W*^*⊤*^. We did not form this inverse explicitly but approximated its action by a fixed number of Neumann iterations, each of which propagates the running estimate once more through *W*^*⊤*^ (derivation in **Supplementary Information S2**; convergence conditions in **S8.2**). The number of iterations is given for each task below.

The spike-forward implicit-differentiation comparator (ID-SNN) used the spiking forward output with the implicit backward path.

### State-Carry Initialization

For sequential inputs, solving each rate fixed point from zero discards the recurrent state built up by earlier inputs. SCI instead initialized the solve at step or phase *m* from the previous solved state,

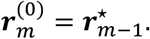

Because the initializer was detached, no gradient flowed into preceding phases. Memory therefore depended only on the carried state and the finite solver dynamics. Control solvers restarted each phase or step from zero, from a random vector, or from a clipped closed-form solution, as specified for each task below.

### Feedback-Spike Nudging

FSN converted the local outer-product structure of the implicit gradient into a forward-running, two-phase spiking process. In the free phase, sensory spikes drove the recurrent and output populations without feedback. The resulting recurrent and output rates are denoted *r*^*h*,−^ and *r*^*o*,−^. The task loss was differentiated only with respect to the free output rate,

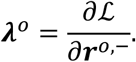

In the feedback phase, signed Bernoulli events encoded this output teaching signal as oppositely directed feedback currents. These events first perturbed output neurons. Output-displacement events then reached recurrent neurons through a learned output-to-hidden pathway. Task-specific gains, probability caps, and feedback timing are given in **Supplementary Information S13–S15**.

Let *r*^*h*,+^ and *r*^*o*,+^ denote the rates measured during the feedback phase. The local displacements were

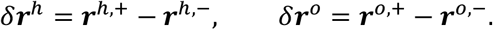

The measured displacements are locally available approximations to the current-level credit. For a fixed set of unclipped neurons, the displacement is proportional to the negative of the exact credit when the recurrent connectivity is symmetric (derivation in **Supplementary Information S4**). Finite spiking windows, feedback gains, and clipping make this correspondence approximate.

The FSN update was

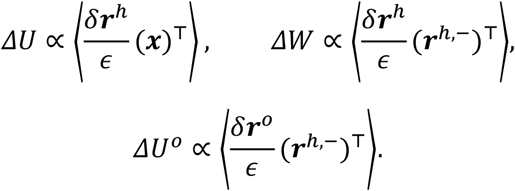

Here, *U*^*o*^ denotes the readout weights from recurrent to output neurons, the denominator in each update is the nudge strength, and angle brackets denote averages over the minibatch or rollout. Bias updates used the displacement alone.

The feedback matrix *B*, which carries output displacements to the recurrent neurons, was learned by Kolen–Pollack mirroring [22,23]. Each feedback synapse, from output neuron *c* to recurrent neuron *i*, received the same activity product as the readout synapse from *i* to *c*, taken in the opposite direction:

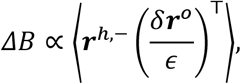

which is the transpose of the readout product in *ΔU*^*o*^. Neither synapse reads the other synapse’s weight. Both updates also included a weight decay. With shared learning and decay rates, the mismatch *B* − *U*^*o⊤*^ shrinks at every update, and the feedback pathway therefore comes to mirror the readout without weight transport (derivation in **Supplementary Information S5**). Centering of the feedback product (cFSN, defined below), unequal learning rates, and optimizer preconditioning break this exact relation.

Centered FSN (cFSN) changed only this feedback-weight product. cFSN subtracts each neuron’s minibatch mean from its free rate and output displacement, which leaves the covariance of the two quantities:

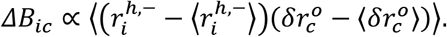

Random-feedback FSN (rFSN) fixed the feedback matrix. Contrastive-Hebbian FSN (CHFSN) replaced the FSN recurrent update by the contrastive-Hebbian update

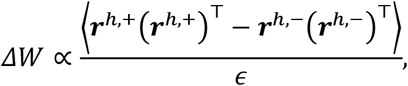

and centered contrastive-Hebbian FSN (cCHFSN) is its centered version, in which each neuron’s minibatch mean is subtracted from its rates before the products are formed. In the Ant task, both controls also replaced the readout update with its contrastive-Hebbian counterpart. Details are given in **Supplementary Information S13–S15**.

### Computational complexity

Let *N* be the number of recurrent neurons and *T* the number of internal timesteps per input or decision. *K* and *J* denote the numbers of fixed-point iterations and Neumann updates in CBIT. Each simulated timestep and each solver iteration requires one pass over the recurrent weights, which costs *O*(*N*^2^) operations. **Table 1** lists the arithmetic per update and the memory per sample beyond the weights. BPTT requires *O*(*TN*^2^) arithmetic and *O*(*TN*) memory for the stored trajectory. E-prop requires the same *O*(*TN*^2^) arithmetic. Its memory is *O*(*N*^2^) in the eligibility-trace form, which keeps one trace per synapse, and *O*(*TN*) in the automatic-differentiation form used in this study. ID-SNN and Rate-BP simulate the spikes forward with *O*(*TN*^2^) arithmetic but differentiate at the rate level, which requires *O*(*N*) memory. ID-SNN additionally performs *J* Neumann updates. CBIT replaces the simulation by *K* iterations and the backward pass by *J* Neumann updates. Its arithmetic is therefore *O*((*K* + *J*)*N*^2^). In the MNIST and continuous-control comparisons, *K* = 10 and *J* = 5 to 12, whereas *T* = 100 to 200, and *K* + *J ≪ T*. Its memory is the solved state, which requires *O*(*N*). SCI changes only the initial state of the iteration. FSN runs the free and feedback phases forward with *O*((*T*_free_ + *T*_feedback_)*N*^2^) arithmetic. It applies one product per synapse and holds only the rates and displacements, which requires *O*(*N*) memory.

### MNIST classification with CBIT

The MNIST [45] classifiers comprised 192 input, 96 recurrent, and 10 output LIF neurons with threshold 1.1 and subtractive reset. Pixel intensities were converted into Bernoulli spikes, which drove the input population through a trainable linear projection. Each image was presented for 100 timesteps, and rates were computed after a 10-step burn-in. The leak amount was 0.1813, corresponding to approximately 4 ms per step for a 20-ms membrane time constant. Class logits were 20 times the output firing rates. The loss was the cross-entropy plus L1 penalties (coefficient 0.1) on the recurrent and output rates. Ten matched seeds were trained for 3,000 updates on minibatches of 256 images drawn from the 60,000 training images and were evaluated by spiking simulation on a fixed 4,096-image test subset. All methods used Adam [79] with weight decay 10^−^5 and gradient clipping at norm 1. The learning rates were 2.5 × 10^−3^ for CBIT and ID-SNN, 2 × 10^−3^ for Rate-BP and e-prop, and 5 × 10^−3^ for BPTT.

CBIT estimated the input rates from two sampled input frames, solved the recurrent state with ten damped iterations (damping factor 0.5), and approximated the credit with twelve Neumann updates. BPTT used a fast-sigmoid surrogate derivative. The e-prop gradients [25] were computed by automatic differentiation with the recurrent spike input detached from the backward pass, the formulation used by its authors for GPU training. Rate-BP replaced the spiking backward pass with a rate-level graph [47]. Training time excluded evaluation, and peak CUDA memory was recorded. Two single-seed sweeps varied the simulation length from 5 to 500 timesteps and the number of recurrent neurons from 10 to 30,000, with all other settings fixed, and recorded training-only time. Implementation details are given in **Supplementary Information S10**.

### Modular addition

We adapted the modular-addition task [48,49] into a sequential, four-phase task to test whether SCI preserves information that is not available from the current input. Each trial contained an inter-cue signal (ICS), a first digit, a second ICS, and a second digit. Both ICS phases signaled addition through a single cue channel, and the target was (*x* + *y*) mod 10. Inputs were rate vectors over one cue channel and ten digit channels and were converted into Bernoulli spikes for spiking evaluation. The network contained 1,024 recurrent and 10 output LIF neurons with threshold 1, subtractive reset, and leak amounts of 0.005 (recurrent) and 0.02 (output). It was trained by CBIT with SCI. The loss, applied after the second digit, combined the cross-entropy of output-rate logits with output-rate and margin terms. Ten seeds were trained with the Adam optimizer [79] for 900 steps on minibatches of 512 trials.

Spiking evaluation used 8,192 trials per model with 40-timestep ICS phases. The theoretical-accuracy comparison used all ten trained models, the 100 ordered digit pairs, and six trials per pair with independent input-rate noise, pooling 16 restarts within each seed for random initialization. The firing-rate scatterplots used one example model and six spiking trials per pair, simulated with the same 40-timestep ICS phases. The PCA visualization used 20 trials per displayed pair on common axes. Phase durations and solver settings are given in **Supplementary Information S11**.

### Continuous-control reinforcement learning

Agents were trained with proximal policy optimization (PPO) [54] and generalized advantage estimation (GAE) [55] on the MuJoCo [50] Ant-v5 and Walker2d-v5 tasks of Gymnasium [51]. Observations were normalized online, split into positive and negative channels, and represented either by expected rates or by Bernoulli spike trains. The policy and value function shared 128 recurrent LIF neurons with subtractive reset and a 20-ms membrane time constant. Each action mean and the value were decoded from the rate difference between 16 positive and 16 negative LIF readout neurons, and actions were drawn from a tanh-squashed Gaussian policy.

The internal simulation of each decision approximately matched the MuJoCo decision interval, with 156 timesteps of 0.32 ms for Ant and 200 timesteps of 0.08 ms for Walker2d. In CBIT+SCI, the recurrent state was solved with ten damped iterations (damping factor 0.25) starting from the previous solved state, and the credit was approximated with five Neumann updates. The two initialization controls started each solve from zero or from a clipped closed-form solution before the same iterations. BPTT differentiated through the spike trajectory with a surrogate derivative. Each method was trained on ten matched seeds for 10 million environment steps. Each update collected 64 steps from each of 64 environments. Policies were evaluated with the spiking network every ten updates. The evaluation reward was the total reward per episode, averaged over 64 deterministic episodes. Implementation details of these experiments and of the analyses below are given in **Supplementary Information S12**.

For the rate-tracking analysis (**Fig. 2d** and **Extended Data Fig. 2c**), the final CBIT+SCI checkpoint of each seed was run as a spiking controller for eight episodes of up to 1,000 steps. At each step, the same ten-iteration solver received the expected input rates and was initialized from its previous solution (SCI) or from zero, or was replaced by the clipped closed-form solution. Recurrent MAE was averaged over the episodes still running at each step and smoothed with a 21-step centered running mean.

For the gradient-variance analysis (**Extended Data Fig. 2f**), each of the ten initial networks of each task received one fixed PPO minibatch of 1,024 transitions and 32 Bernoulli input-spike realizations, which were shared by BPTT and ID-SNN. CBIT+SCI used the expected input rates. Relative variance was the summed coordinate-wise variance of the gradient across realizations divided by the squared norm of the mean gradient, giving one value per network.

### Development of touchdown-aligned activity

To follow the development of locomotor activity (**Fig. 2e,f** and **Extended Data Fig. 2d,e**), the CBIT+SCI checkpoints of all ten seeds were replayed at 15 points during training, with eight spiking episodes per checkpoint. Recurrent firing rates, limb-tip positions relative to the torso, and applied controls were aligned to the touchdowns of a reference limb, detected from MuJoCo ground contacts, and smoothed with a Gaussian kernel (SD, 25 ms). At each checkpoint, four episodes were used to order the neurons by the time of their peak rate and to z-score their rates, and the other four episodes provided the displayed averages. The sequentiality index [56] was computed from the held-out rates of each seed and checkpoint before averaging across seeds.

### MNIST classification with FSN

The local MNIST experiments used 784 Bernoulli sensory channels, 256 recurrent LIF neurons, and 10 output LIF neurons with leak amount 0.05, threshold 1, and subtractive reset. Each training update comprised a 100-timestep free phase and a 100-timestep feedback phase, with a 10-timestep burn-in. Because the same sensory spike sequence was replayed in both phases, the rate difference reflected the feedback rather than input resampling. The output teaching signal was the derivative of the cross-entropy loss, computed from logits equal to 20 times the free output rates, with respect to those rates. This signal was not backpropagated through the recurrent dynamics. It was delivered as signed Bernoulli feedback events with nudge strength 0.15.

FSN and cFSN were compared with rFSN, CHFSN, BPTT, e-prop with an adaptive feedback matrix, and a frozen-recurrent control, using ten matched seeds, minibatches of 128 images, and 24,000 updates. The local rules applied plain parameter updates without an adaptive optimizer or gradient clipping. The frozen-recurrent control trained only the readout weights and output bias, keeping the input and recurrent weights, recurrent bias, and feedback matrix at their initial values. Accuracy was evaluated on the full 10,000-image test set. Alignment diagnostics compared each local update with the surrogate-BPTT gradient computed on the same minibatch and spike realization. Implementation details are given in **Supplementary Information S13**.

### Supervised Reacher imitation with FSN

A frozen SAC teacher [65] with two hidden layers of 256 ReLU units supplied target actions for Reacher-v5, in which the initial joint angles were randomized. The student generated every environment action. The ten observation variables were split into positive and negative channels, each represented by 96 redundant Bernoulli sensory axons projecting to 256 recurrent LIF neurons. Each of the two actions was decoded from the rate difference between 16 positive and 16 negative output LIF neurons through a hyperbolic tangent function. The loss was the squared error between the student and teacher actions plus a weak output-rate regularizer. Its derivative with respect to the free output rates was delivered as signed feedback spikes with nudge strength 0.04.

During training, each environment step comprised 150 free and 150 feedback timesteps with a 15-step burn-in and zero reset, and the feedback-phase state was carried to the next step. Local updates were accumulated over 64 steps of 256 environments and applied with Adam. Six configurations, including a control with frozen input and recurrent weights, were trained on ten seeds for 80 million environment steps. Training-time evaluations used fixed weights and feedback-state carry over 64 episodes every 20 updates. The checkpoint with the highest logged reward and the final checkpoint were then evaluated on 512 fresh episodes without teaching feedback, and once more with feedback-state carry as a control. In the evaluation without teaching feedback, actions and the carried state came only from the 150-timestep free phase. Weights were fixed, and recurrent states were reset at episode boundaries. The diagnostics in **Extended Data Fig. 4** compared the combined input and recurrent updates with a surrogate-BPTT gradient and measured the centered feedback– readout alignment. Implementation details are given in **Supplementary Information S14**.

### Ant reinforcement learning with FSN

The Ant-v5 experiment trained a spiking recurrent actor–critic with PPO-derived teaching signals. The 105 normalized observation variables were split into positive and negative channels, each represented by 32 redundant Bernoulli sensory axons. The network contained 256 recurrent and 288 output LIF neurons with leak amount 0.05 and zero reset, with 16 positive and 16 negative output neurons for each of the eight actions and the value. At each environment step, a 225-timestep free phase generated the action and was followed by a 225-timestep feedback-state window. The final feedback state was then carried to the next step.

Training used 256 environments, 64 steps per rollout, and 20 million environment steps. During rollout and evaluation, the feedback-state window received no teaching signal. During PPO optimization, the stored input spikes and initial states of each transition were replayed. The PPO loss was then differentiated with respect to the free output rates. This teaching signal was normalized, clipped, and delivered as signed feedback spikes with nudge strength 0.04. The resulting local updates were applied with Adam. cFSN, FSN, rFSN, CHFSN, and cCHFSN were trained on ten seeds and evaluated every 20 updates over 64 deterministic spiking episodes. Implementation details are given in **Supplementary Information S15**.

### Biological spine-data analysis

We analyzed the iGluSnFR3–RCaMP2 longitudinal motor-learning dataset of Hedrick et al. [4]. Imaging accompanied daily rewarded lever-press training on days 1–3, 6–8, and 11–13. Each field was revisited five days later. Each spine transition paired an early with a middle session or a middle with a late session. Spine-local iGluSnFR3 activity was paired with RCaMP2 activity pooled from shaft regions of interest along the parent branch, with the same calcium trace assigned to spines on that branch. The source study estimated apparent spine area from integrated spine fluorescence normalized by nearby shaft fluorescence and corrected for pixel size. We used next-session minus current-session area as the signed structural outcome, and retention or elimination as a separate outcome.

Active periods were the binary classifications published with the source study [4]. We counted inactive-to-active transitions and divided their number by window duration, giving equal weight to each detected event. Event rates used matched 2.5-s pre-reward and post-reward windows, separated from reward by 0.1-s guards. Event rates were standardized within each branch and learning stage. With standardized pre- and post-reward iGluSnFR3 rates *a* and *b* and RCaMP2 rates *x* and *y*, the FSN product was

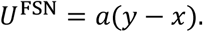

Products were computed for each rewarded trial and averaged within each spine transition. Six comparator rules were built from the same variables. Event selection, scaling, the comparator formulas, and the reconstruction of spine area from the source data are described in **Supplementary Information S16**.

Spine-area analyses used the 517 surviving transitions from seven mice, and spine fate was analyzed across 563 transitions, including 46 eliminations. Spearman and Pearson correlations pooled all transitions. Unadjusted and covariate-adjusted ordinary-least-squares models used a finite-sample-corrected covariance clustered by mouse, with t tests on six degrees of freedom. The covariates were current log spine area, nearby coactivity, movement-relatedness, and learning phase. Confidence intervals for grouped means came from 10,000 bootstrap resamples of mice.

### Computational resources

Network training and runtime benchmarks were performed on the TSUBAME4.0 supercomputer at the Institute of Science Tokyo, whose HPE Cray XD665 nodes are equipped with AMD EPYC 9654 CPUs and NVIDIA H100 GPUs. Each job used one H100 GPU, and all methods of a runtime or memory comparison were run on the same node type.

### Statistics and reproducibility

Simulation experiments used ten random seeds unless stated otherwise. Wilcoxon signed-rank tests with Holm correction [80] were used for paired comparisons. Surviving and eliminated spines were compared using a one-sided Wilcoxon rank-sum test for higher products in surviving spines.

Unless stated otherwise in a legend, lines and shaded bands show the mean ± SEM, and box plots show the median (center line), interquartile range (box), and the most extreme values within 1.5 times the interquartile range (whiskers), with points for individual seeds. Asterisks in figures mark Holm-corrected p < 0.05 (*) and p < 0.01 (**).

## Acknowledgements

We thank T. Harumoto for helpful discussions. This work was supported by MEXT/JSPS KAKENHI JP24H02156 (R.H.), JP24KK0186 (R.H.), JP26H01161 (R.H.), and JP26K23436 (R.H.); AMED JP26wm0625405 (R.H. and J.I.); JST FOREST JPMJFR231X (R.H.); the Nakatani Foundation (R.H.); the Takeda Science Foundation (R.H.); the Precise Measurement Technology Promotion Foundation (R.H.); the Tateishi Science and Technology Foundation (R.H.); collaborative research by young researchers through GX Frontier Visionary Initiative, Science Tokyo (R.H.). H.I. was supported by the Toyota Riken Overseas Scholarship. F.I. was supported by the Funai Overseas Scholarship of the Funai Foundation for Information Technology. This work utilized the TSUBAME4.0 supercomputer at the Institute of Science Tokyo for analysis. We used ChatGPT-5 (OpenAI) and Claude (Anthropic) for English and Japanese translation during manuscript preparation.

## Author Contributions

H.I. and F.I. performed the numerical simulations and analyses. H.I. developed the computational methods. Y.I. and J.I. supervised the project. R.H. and H.I. wrote the manuscript with input from all authors.

## Competing Interests

The authors declare no competing interests.

## Data Availability

Data will be available upon acceptance.

## Code Availability

Code will be available upon acceptance.

## Supplementary Information

The **Supplementary Information** provides the mathematical derivations and pseudocode for the current-balance rate equation, CBIT, SCI, and FSN (S1–S8) and the detailed implementations, hyperparameters, and analysis procedures for all experiments (S9–S16).

## Supplementary Videos

**Supplementary Video 1**. Ant locomotion by an RSNN trained with CBIT+SCI. Evaluation episode of the final checkpoint of training seed 3 (total reward 5,402). Left, the Ant, the applied actions, and the actions of the last 2 s. Right, spikes of the recurrent neurons and of the positive and negative actor and critic output populations in the three most recent action windows.

**Supplementary Video 2**. Walker2d locomotion by an RSNN trained with CBIT+SCI. Evaluation episode of the final checkpoint of training seed 4 (total reward 4,880). Layout as in **Supplementary Video 1**, with six actions and 200 internal timesteps per action window.

**Supplementary Video 3**. Reacher control by an RSNN trained with cFSN to imitate a SAC teacher. Six illustrative rollouts of the final checkpoint of training seed 3, with fixed weights and without teacher-derived feedback. Quantitative evaluation without teaching feedback is shown in **Fig. 4c**. Left, the arm (target, red; fingertip, green), the fingertip-to-target distance, and the applied joint torques. Right, spikes of 100 example recurrent neurons and of the positive and negative output populations for each joint in the three most recent action windows (150 internal timesteps per window).

**Supplementary Video 4**. Ant locomotion by an RSNN trained with cFSN from reward. Evaluation episode of the final checkpoint of training seed 4 (total reward 4,485). Layout as in **Supplementary Video 1**, with 225 internal timesteps per action window.

## Notes

### Competing Interest Statement

The authors have declared no competing interest.

