## Supplementary information for "Recurrent Spiking Neural Networks for Continuous Motor Control: A Biologically Plausible Model of Motor Cortex"

This Supplementary Information has two parts. Part I (Sections S1–S8) gives the mathematical derivations of the current-balance rate equation, Current-Balance Implicit Training (CBIT), State-Carry Initialization (SCI) and Feedback-Spike Nudging (FSN), together with pseudocode. Part II (Sections S9–S16) gives the experiment-specific implementations, hyperparameters, analysis procedures and statistical tests. Supplementary Tables 1–10 appear in the corresponding sections of Part II. Figure numbers (Fig. and Extended Data Fig.) refer to the main text.

#### Contents

**Part I. Mathematical methods** **2**

**S1 The rate equation and its numerical solution** **2**

**S2 Implicit differentiation and CBIT** **3**

**S3 State-Carry Initialization** **5**

**S4 From feedback spikes to the FSN update** **5**

**S5 Kolen–Pollack mirroring** **8**

**S6 Contrastive controls** **9**

**S7 Algorithms and update schedules** **9**

**S8 Conditions and additional identities** **11**

**Part II. Experimental methods** **13**

**S9 Benchmarking theoretical equilibrium firing-rate estimates** **13**

**S10 MNIST classification with Current-Balance Implicit Training** **16**

**S11 Modular addition with State-Carry Initialization** **19**

**S12 Continuous-control reinforcement learning with CBIT** **22**

**S13 Biologically local MNIST learning with Feedback-Spike Nudging** **27**

**S14 Supervised Reacher imitation with feedback-state carry** **31**

**S15 Biologically local Ant reinforcement learning** **34**

**S16 Biological spine-data validation of the local-product rule** **37**

**Supplementary References** **39**

### Part I. Mathematical methods

We first derive the current-balance rate equation and explain how we solve it. We then differentiate the solved state to obtain the CBIT weight updates. This identifies the postsynaptic credit that FSN replaces with a neuron's response to feedback spikes. The FSN derivation follows the signal through the output population, the feedback pathway and the recurrent population, before giving the local plasticity rule. Sections S9–S16 provide the experiment-specific parameters and statistical procedures.

#### S1 The rate equation and its numerical solution

##### S1.1 Network variables and the current-balance transfer function

We consider input, recurrent and output populations with rates $\boldsymbol{x} \in \mathbb{R}^M$, $\boldsymbol{r} \in \mathbb{R}^N$ and $\boldsymbol{r}^o \in \mathbb{R}^C$, respectively. Rates are measured in spikes per internal timestep; dividing by the timestep duration in seconds converts them to hertz. Input weights $U \in \mathbb{R}^{N\times M}$, recurrent weights $W \in \mathbb{R}^{N\times N}$ and readout weights $U^o \in \mathbb{R}^{C\times N}$ determine the mean synaptic currents:

$$\boldsymbol{h} = W\boldsymbol{r} + U\boldsymbol{x} + \boldsymbol{b}, \qquad \boldsymbol{h}^o = U^o\boldsymbol{r} + \boldsymbol{b}^o. \tag{S1.1}$$

We use column vectors throughout. In Part II, the readout matrix $U^o$ is also written $V$.

To relate current to firing rate, consider a subtractive-reset LIF neuron:

$$v_{t+1,i} = (1-\beta_i)v_{t,i} + h_{t,i} - \theta_i s_{t+1,i}, \tag{S1.2}$$

where $v$ is membrane voltage, $\beta_i$ is the leak amount per step, $\theta_i > 0$ is the threshold and $s_{t+1,i} \in \{0,1\}$ is the emitted spike. Consider a long window of $T$ timesteps during which the input is stationary with mean current $\bar{h}_i$. Averaging Eq. (S1.2) over the window, the time average of the spike train $s_{t+1,i}$ converges to the firing rate $r_i$, the time average of the voltage $v_{t,i}$ converges to its mean $\langle v_i\rangle$, and the average of the increment $v_{t+1,i} - v_{t,i}$ vanishes as $T$ grows because the voltage is bounded. The averaged update therefore reads $0 = -\beta_i\langle v_i\rangle + \bar{h}_i - \theta_i r_i$, that is,

$$\bar{h}_i = \theta_i r_i + \beta_i\langle v_i\rangle. \tag{S1.3}$$

Mean input current therefore compensates for both spike resets and membrane leak; the finite-window form, including the boundary term, is given in Section S8.1. Approximating the mean subthreshold voltage by the midpoint $\langle v_i\rangle \approx \theta_i/2$ gives $r_i \approx \bar{h}_i/\theta_i - \beta_i/2$. Because each neuron emits at most one spike per timestep, we clip this estimate to the allowed range:

$$\Psi_i(h_i) = \operatorname{clip}\left(\frac{h_i}{\theta_i} - \frac{\beta_i}{2}, 0, 1\right). \tag{S1.4}$$

This is the current-balance transfer function used in the rate model. The effect of zero reset is given in Section S8.1.

##### S1.2 Solving the self-consistent rate equation

The transfer function $\Psi_i$ is defined for each neuron $i$. In all experiments, the neurons of a population share the same threshold $\theta$ and leak amount $\beta$ (values are given in Part II), and we write $\Psi(\boldsymbol{h})$ for the vector whose $i$th element is $\Psi_i(h_i)$; vectors are set in bold throughout. Recurrent firing contributes to its own input current. Substituting Eq. (S1.1) into the transfer function therefore gives the self-consistent equation

$$\boldsymbol{r}^\star = \Psi(W\boldsymbol{r}^\star + U\boldsymbol{x} + \boldsymbol{b}). \tag{S1.5}$$

We solve this equation with damped fixed-point iterations. Starting from $\boldsymbol{r}^{(0)}$, each iteration combines the current estimate with the rate predicted from its synaptic current:

$$\boldsymbol{r}^{(k+1)} = \mathrm{clip}\Big[(1-\eta)\boldsymbol{r}^{(k)} + \eta\Psi(W\boldsymbol{r}^{(k)} + U\boldsymbol{x} + \boldsymbol{b}), 0, 1\Big], \qquad 0 < \eta \le 1. \tag{S1.6}$$

The relaxation factor $\eta$ controls how far the estimate moves at each iteration. After the specified number $K$ of iterations, the resulting recurrent rate supplies the output layer:

$$\boldsymbol{r}^o = \Psi^o(U^o\boldsymbol{r} + \boldsymbol{b}^o), \qquad \boldsymbol{r} = \boldsymbol{r}^{(K)}. \tag{S1.7}$$

Training uses fixed iteration budgets. In the derivation below, $\boldsymbol{r}$ denotes an equilibrium when deriving an exact implicit gradient; the implementation evaluates that formula at the finite solved state. The iteration counts, relaxation factors and initial states are specified in Sections S10–S12. SCI changes the initializer, as explained in Section S3.

### S2 Implicit differentiation and CBIT

To train the weights, we need to know how a weight change affects the loss through the solved rates. The transfer function is linear within each of its three branches. Away from the clipping boundaries, its gain is

$$D = \mathrm{diag}(D_1, \ldots, D_N), \qquad D_i = \begin{cases} 1/\theta_i, & 0 < h_i/\theta_i - \beta_i/2 < 1, \\ 0, & h_i/\theta_i - \beta_i/2 < 0 \text{ or } h_i/\theta_i - \beta_i/2 > 1. \end{cases} \tag{S2.1}$$

The output gain $D^o$ is defined in the same way. Thus a small current change produces $d\boldsymbol{r} = D\,d\boldsymbol{h}$ or $d\boldsymbol{r}^o = D^o\,d\boldsymbol{h}^o$ while the branch remains unchanged.

#### S2.1 The readout gradient

We begin with the nonrecurrent output layer. Let $\mathcal{L}$ be the task loss, and let $\boldsymbol{\lambda}^o = \partial\mathcal{L}/\partial\boldsymbol{r}^o$ be its derivative with respect to the output firing rates. The total differential of the readout is

$$d\boldsymbol{h}^o = dU^o\,\boldsymbol{r} + U^o\,d\boldsymbol{r} + d\boldsymbol{b}^o, \tag{S2.2}$$
$$d\boldsymbol{r}^o = D^o(dU^o\,\boldsymbol{r} + U^o\,d\boldsymbol{r} + d\boldsymbol{b}^o). \tag{S2.3}$$

For the derivative with respect to $U^o$ and $\boldsymbol{b}^o$, we hold the recurrent state fixed, which gives $d\boldsymbol{r} = 0$. Applying the chain rule to the loss then gives

$$\begin{aligned} d\mathcal{L} &= (\boldsymbol{\lambda}^o)^\top d\boldsymbol{r}^o \\ &= (\boldsymbol{\lambda}^o)^\top D^o(dU^o\,\boldsymbol{r} + d\boldsymbol{b}^o) \\ &= (\boldsymbol{p}^o)^\top(dU^o\,\boldsymbol{r} + d\boldsymbol{b}^o), \qquad \boldsymbol{p}^o = D^o\boldsymbol{\lambda}^o. \end{aligned} \tag{S2.4}$$

Here $\boldsymbol{p}^o$ is the loss sensitivity to an output current, which we call *current-level credit.* Reading off the coefficient of each changed weight or bias yields

$$\frac{\partial\mathcal{L}}{\partial U^o} = \boldsymbol{p}^o\boldsymbol{r}^\top, \qquad \frac{\partial\mathcal{L}}{\partial\boldsymbol{b}^o} = \boldsymbol{p}^o. \tag{S2.5}$$

In particular, the gradient of the synapse from recurrent neuron $j$ to output neuron $k$ is $p_k^o r_j$. This separates a postsynaptic credit from the presynaptic firing rate.

#### S2.2 The recurrent and input-weight gradients

Changing a recurrent weight also changes the recurrent activity that returns to the neuron. We retain this dependence by differentiating the self-consistent equation. At fixed input $\boldsymbol{x}$,

$$
\begin{aligned}
d\boldsymbol{h} &= dW\,\boldsymbol{r} + W\,d\boldsymbol{r} + dU\,\boldsymbol{x} + d\boldsymbol{b}, && \text{(S2.6)}\\
d\boldsymbol{r} &= D(dW\,\boldsymbol{r} + W\,d\boldsymbol{r} + dU\,\boldsymbol{x} + d\boldsymbol{b}), && \text{(S2.7)}\\
(I - DW)d\boldsymbol{r} &= D(dW\,\boldsymbol{r} + dU\,\boldsymbol{x} + d\boldsymbol{b}). && \text{(S2.8)}
\end{aligned}
$$

If $I - DW$ is invertible, this determines how the equilibrium changes:

$$d\boldsymbol{r} = (I - DW)^{-1}D(dW\,\boldsymbol{r} + dU\,\boldsymbol{x} + d\boldsymbol{b}). \tag{S2.9}$$

Let $\boldsymbol{\lambda}^h = \partial\mathcal{L}/\partial\boldsymbol{r}$ be the rate-level loss derivative entering the recurrent population. For a loss defined only on the output, the readout chain rule gives $\boldsymbol{\lambda}^h = (U^o)^\top\boldsymbol{p}^o$. A penalty defined directly on recurrent rates adds its derivative here. Substituting Eq. (S2.9) into the loss differential gives

$$
\begin{aligned}
d\mathcal{L} &= (\boldsymbol{\lambda}^h)^\top d\boldsymbol{r}\\
&= (\boldsymbol{\lambda}^h)^\top (I - DW)^{-1}D(dW\,\boldsymbol{r} + dU\,\boldsymbol{x} + d\boldsymbol{b})\\
&= (\boldsymbol{p}^h)^\top (dW\,\boldsymbol{r} + dU\,\boldsymbol{x} + d\boldsymbol{b}), && \text{(S2.10)}
\end{aligned}
$$

where the recurrent current-level credit is

$$\boldsymbol{p}^h = D(I - W^\top D)^{-1}\boldsymbol{\lambda}^h = (I - DW^\top)^{-1}D\boldsymbol{\lambda}^h. \tag{S2.11}$$

The two forms are equivalent even when some gains are zero (Section S8.3). Comparing coefficients in Eq. (S2.10) now gives

$$\frac{\partial\mathcal{L}}{\partial W} = \boldsymbol{p}^h\boldsymbol{r}^\top, \qquad \frac{\partial\mathcal{L}}{\partial U} = \boldsymbol{p}^h\boldsymbol{x}^\top, \qquad \frac{\partial\mathcal{L}}{\partial \boldsymbol{b}} = \boldsymbol{p}^h. \tag{S2.12}$$

These products have the same structure as the readout gradient; the extra computation lies in obtaining $\boldsymbol{p}^h$.

#### S2.3 Computing the credit with Neumann iterations

We approximate the action of the inverse in Eq. (S2.11) with Neumann iterations, rather than constructing a matrix inverse. Define an auxiliary vector $\boldsymbol{a}$ by

$$(I - W^\top D)\boldsymbol{a} = \boldsymbol{\lambda}^h, \qquad \boldsymbol{p}^h = D\boldsymbol{a}. \tag{S2.13}$$

Rearranging gives $\boldsymbol{a} = \boldsymbol{\lambda}^h + W^\top D\boldsymbol{a}$. Starting from $\boldsymbol{a}^{(0)} = \boldsymbol{\lambda}^h$, we apply this relation repeatedly:

$$\boldsymbol{a}^{(j+1)} = \boldsymbol{\lambda}^h + W^\top D\boldsymbol{a}^{(j)}, \qquad \boldsymbol{p}^h \approx D\boldsymbol{a}^{(J)}. \tag{S2.14}$$

After $J$ recurrence updates, $\boldsymbol{a}^{(J)} = \sum_{j=0}^{J}(W^\top D)^j\boldsymbol{\lambda}^h$: the estimate contains the initial term and $J$ recurrently propagated terms. Each term propagates the loss derivative once more through the transposed recurrent matrix $W^\top$. This repeated propagation through $W^\top$ is the non-local computation that distinguishes the recurrent credit from the readout gradient, and it is the step that FSN replaces by the circuit's own response (Section S4). This is the backward calculation used by CBIT. It follows the forward rate solve in Eq. (S1.6); it does not differentiate through that sequence of forward iterates. Convergence conditions and the finite-iteration approximation are stated in Section S8.2.

### S3 State-Carry Initialization

For a sequence of inputs, restarting every solve from zero can discard the network state formed by earlier inputs. We instead initialize the next solve with the rate reached in the preceding phase or environment step:

$$\boldsymbol{r}_m^{(0)} = \mathrm{stopgrad}(\boldsymbol{r}_{m-1}), \qquad \boldsymbol{r}_m = F_m^{K_m}(\boldsymbol{r}_m^{(0)}), \tag{S3.1}$$

where $F_m$ is one damped iteration of Eq. (S1.6) for input $\boldsymbol{x}_m$, and $K_m$ is the number of iterations. The first phase uses the task's specified initial state. The stop-gradient operation stopgrad prevents backpropagation through the preceding phase while preserving its value as the next initializer.

With a finite number of iterations, the new rate can depend on both the current input and the carried state. If multiple fixed points exist, initialization can also influence which one is reached. Thus the memory retained by SCI does not require every phase to converge to a distinct equilibrium. During explicit-spike evaluation, the network carries its own membrane and spike states through the input sequence. The rate-state calculation predicts those firing rates; it does not initialize the spiking network with the solved rate vector.

### S4 From feedback spikes to the FSN update

The implicit gradient has identified the quantity each synapse needs: presynaptic activity multiplied by postsynaptic current-level credit. We now ask whether the postsynaptic neuron can measure an approximation to that credit through its own response to feedback. We first explain this for the output layer, then trace how the output response is sent back to the recurrent population.

#### S4.1 Free activity and the output teaching signal

During the *free phase*, sensory spikes drive the network without teaching feedback. We record input rates $\boldsymbol{x}^-$, recurrent rates $\boldsymbol{r}^-$ and output rates $\boldsymbol{r}^{o,-}$, together with the final spiking state. Superscript $-$ denotes the free phase; $+$ will denote the subsequent feedback phase.

The task supplies an output rate-level teaching signal

$$\boldsymbol{\lambda}^o = \frac{\partial \mathcal{J}}{\partial \boldsymbol{r}^{o,-}}, \tag{S4.1}$$

where $\mathcal{J}$ is the applicable output objective. It compares the output with a class or teacher action in supervised learning, or uses the PPO objective in Ant. FSN uses this signal to generate feedback spikes. Positive and negative feedback events deliver currents of opposite sign, which can be represented by the difference of two nonnegative spike populations. We write their mean current into the output neurons as

$$\bar{\boldsymbol{f}}^o = -\varepsilon_o \boldsymbol{\lambda}^o. \tag{S4.2}$$

Here $\bar{\boldsymbol{f}}^o$ is the additional synaptic current produced by feedback, and $\varepsilon_o > 0$ includes the output nudge strength and gain. The negative sign makes the perturbation oppose an increase in the loss. The signed-event implementation is specified below in Section S4.6.

#### S4.2 How output displacement supplies the readout update

To see what the output neurons contribute, first hold their recurrent input at the free rate $\boldsymbol{r}^-$. Their free and feedback rate equations are then

$$\boldsymbol{r}^{o,-} = \Psi^o(U^o\boldsymbol{r}^- + \boldsymbol{b}^o), \tag{S4.3}$$
$$\boldsymbol{r}^{o,+} = \Psi^o(U^o\boldsymbol{r}^- + \boldsymbol{b}^o + \bar{\boldsymbol{f}}^o). \tag{S4.4}$$

Within unchanged transfer-function branches, subtracting the two equations multiplies the added current by the output gain:

$$\begin{aligned}\delta\boldsymbol{r}^o &\equiv \boldsymbol{r}^{o,+} - \boldsymbol{r}^{o,-}\\ &= D^o\bar{\boldsymbol{f}}^o\\ &= -\varepsilon_o D^o\boldsymbol{\lambda}^o = -\varepsilon_o\boldsymbol{p}^o.\end{aligned} \tag{S4.5}$$

Thus the output firing-rate change measures the negative current-level credit, up to the feedback scale. The output neuron's response supplies $D^o$; the feedback population need not transmit that derivative separately.

Gradient descent would change the readout by $-\eta_o\boldsymbol{p}^o(\boldsymbol{r}^-)^\top$. Substituting $\boldsymbol{p}^o = -\delta\boldsymbol{r}^o/\varepsilon_o$ gives the delta-readout rule

$$\Delta U^o = \eta_o\left\langle\frac{\delta\boldsymbol{r}^o}{\varepsilon_o}(\boldsymbol{r}^-)^\top\right\rangle_{\mathcal{B}}, \qquad \Delta\boldsymbol{b}^o = \eta_o\left\langle\frac{\delta\boldsymbol{r}^o}{\varepsilon_o}\right\rangle_{\mathcal{B}}. \tag{S4.6}$$

The brackets denote averaging over the training batch, and $\Delta$ denotes an increment added to a parameter. A readout synapse therefore uses the presynaptic free-phase rate and the postsynaptic change caused by feedback. We held the hidden input fixed to derive this correspondence; the experimental feedback phase couples both populations, as described in Section S4.5.

#### S4.3 Routing the output response to recurrent neurons

The recurrent neurons also need teaching input. We use the measured output response to form

$$\widehat{\boldsymbol{p}}^o = -\frac{\delta\boldsymbol{r}^o}{\varepsilon_o}. \tag{S4.7}$$

A feedback population encodes this estimate as spikes and projects to the recurrent population through its own synaptic matrix $B \in \mathbb{R}^{N\times C}$. The mean teaching vector delivered through that projection, and the resulting additional current, are

$$\boldsymbol{\lambda}^h_{\text{fb}} = B\widehat{\boldsymbol{p}}^o, \qquad \bar{\boldsymbol{f}}^h = -\varepsilon_h\boldsymbol{\lambda}^h_{\text{fb}}. \tag{S4.8}$$

The entry $B_{ik}$ weights the output-related feedback sent to recurrent neuron $i$ from feedback channel $k$. The positive scale $\varepsilon_h$ sets the recurrent nudge strength. These feedback synapses are independent of the readout synapses; their Kolen–Pollack learning rule is given in Section S5.

This defines $\boldsymbol{\lambda}^h_{\text{fb}}$ as a signal delivered by feedback, whereas $\boldsymbol{\lambda}^h = (U^o)^\top\boldsymbol{p}^o$ in Section S2 is the exact rate-level loss derivative. The two agree in the ideal case that the measured output credit equals $\boldsymbol{p}^o$ and $B = (U^o)^\top$. The implemented circuit measures the output response and uses its own feedback weights.

#### S4.4 Deriving the recurrent displacement and local update

To derive the response to this teaching current, first hold its mean $\bar{\boldsymbol{f}}^h$ fixed while the recurrent population settles. Sensory input and synaptic weights are the same in the two phases:

$$\boldsymbol{r}^- = \Psi(W\boldsymbol{r}^- + U\boldsymbol{x} + \boldsymbol{b}), \tag{S4.9}$$

$$\boldsymbol{r}^+ = \Psi(W\boldsymbol{r}^+ + U\boldsymbol{x} + \boldsymbol{b} + \bar{\boldsymbol{f}}^h). \tag{S4.10}$$

The difference between the two input currents is $W(\boldsymbol{r}^+ - \boldsymbol{r}^-) + \bar{\boldsymbol{f}}^h$. If each neuron remains in the same branch of the piecewise-linear transfer, its rate difference is its gain times this current difference. Writing $\delta\boldsymbol{r} = \boldsymbol{r}^+ - \boldsymbol{r}^-$ therefore gives, step by step,

$$\begin{aligned}\delta\boldsymbol{r} &= \Psi(W\boldsymbol{r}^+ + U\boldsymbol{x} + \boldsymbol{b} + \bar{\boldsymbol{f}}^h) - \Psi(W\boldsymbol{r}^- + U\boldsymbol{x} + \boldsymbol{b}) \\ &= D\big[W(\boldsymbol{r}^+ - \boldsymbol{r}^-) + \bar{\boldsymbol{f}}^h\big] \\ &= DW\delta\boldsymbol{r} - \varepsilon_h D\boldsymbol{\lambda}^h_{\text{fb}}.\end{aligned} \tag{S4.11}$$

Moving the recurrent term to the left gives

$$(I - DW)\delta\boldsymbol{r} = -\varepsilon_h D\boldsymbol{\lambda}^h_{\text{fb}}. \tag{S4.12}$$

When the inverse exists, the measured recurrent credit is consequently

$$\widehat{\boldsymbol{p}}^h \equiv -\frac{\delta\boldsymbol{r}}{\varepsilon_h} = (I - DW)^{-1}D\boldsymbol{\lambda}^h_{\text{fb}}. \tag{S4.13}$$

Compare this with the exact credit $\boldsymbol{p}^h = (I - DW^\top)^{-1}D\boldsymbol{\lambda}^h$ in Eq. (S2.11). Feedback spreads through the circuit's actual $W$, while the exact gradient uses $W^\top$. For symmetric recurrence and matching teaching vectors the two expressions coincide. For an asymmetric circuit, the measured displacement provides an approximation whose alignment can be evaluated experimentally.

Substituting this measured credit into the negative-gradient outer product gives the FSN recurrent update:

$$\Delta W = -\eta_h \left\langle \widehat{\boldsymbol{p}}^h (\boldsymbol{r}^-)^\top \right\rangle_{\mathcal{B}} = \eta_h \left\langle \frac{\delta\boldsymbol{r}}{\varepsilon_h} (\boldsymbol{r}^-)^\top \right\rangle_{\mathcal{B}}. \tag{S4.14}$$

The same substitution gives input and bias increments,

$$\Delta U = \eta_h \left\langle \frac{\delta\boldsymbol{r}}{\varepsilon_h} (\boldsymbol{x}^-)^\top \right\rangle_{\mathcal{B}}, \qquad \Delta\boldsymbol{b} = \eta_h \left\langle \frac{\delta\boldsymbol{r}}{\varepsilon_h} \right\rangle_{\mathcal{B}}. \tag{S4.15}$$

For an individual recurrent synapse, the rule is $\Delta W_{ij} = \eta_h \delta r_i r_j^- / \varepsilon_h$, before batch averaging. Neuron $j$ supplies its free-phase activity and neuron $i$ supplies its own feedback-induced change. No synapse needs the analytically calculated vector $\boldsymbol{p}^h$. These equations specify the local increments before the task's optimizer, decay and parameter constraints.

#### S4.5 The coupled feedback episode used in the experiments

The preceding derivation separated the populations to explain the source of each update. In the experiments, output and recurrent neurons continue evolving during the same feedback episode. The feedback population reads a running output rate after burn-in:

$$\widehat{\boldsymbol{p}}^o_t = -\frac{\bar{\boldsymbol{r}}^{o,+}_{1:t} - \boldsymbol{r}^{o,-}}{\varepsilon_o}. \tag{S4.16}$$

It converts the processed estimate into signed events $\boldsymbol{q}_t^o$, whose mean represents that estimate when probability caps are inactive. These events produce recurrent feedback on the next timestep:

$$\boldsymbol{f}_{t+1}^h = -\varepsilon_h B \boldsymbol{q}_t^o. \tag{S4.17}$$

The feedback current is initially zero. The one-step delay lets the output response at time $t$ influence subsequent recurrent activity without an instantaneous loop.

As recurrent activity changes, it also changes the drive to the output layer. Within a fixed rate-model branch, the coupled output displacement becomes

$$\delta \boldsymbol{r}^o = D^o(U^o \delta \boldsymbol{r} - \varepsilon_o \boldsymbol{\lambda}^o). \tag{S4.18}$$

The first term is the response to changed recurrent activity; the second is the response to direct output feedback. Thus $-\delta \boldsymbol{r}^o/\varepsilon_o$ contains both contributions. The implemented local rules use these measured coupled displacements. The isolated derivation explains their connection to the implicit gradient, while the coupled finite-spike implementation is assessed by the reported learning and update-alignment measurements. Section S8.4 gives the stationary coupled equations.

##### S4.6 Signed feedback events

For a desired signed mean signal $z$, event amplitude $a_f > 0$ and probability cap $p_{\max} \le 1$, the signed-Bernoulli encoder is

$$\mathcal{S}(z) = a_f \operatorname{sign}(z)\xi, \qquad \xi \sim \text{Bernoulli}\left(\min\{|z|/a_f, p_{\max}\}\right). \tag{S4.19}$$

When the cap is inactive, $\mathbb{E}[\mathcal{S}(z)] = z$. With capping,

$$\mathbb{E}[\mathcal{S}(z)] = a_f \operatorname{sign}(z) \min(|z|/a_f, p_{\max}).$$

Output events encode the negative output teaching signal, and the second feedback population encodes the measured output displacement as described above. Signal normalization, clipping, event amplitudes and burn-in depend on the task and are specified in Sections S13–S15.

#### S5 Kolen–Pollack mirroring

The remaining question is how to learn the feedback matrix $B$ without copying the readout weights. We use a local coactivity rule, termed *Kolen–Pollack mirroring*, that pairs the same phase measurements in the two directions. Define the scaled output displacement $\boldsymbol{\zeta}^o = \delta \boldsymbol{r}^o/\varepsilon_o$. The readout and feedback increments, including their respective decay terms, are

$$\Delta U^o = \eta_o \left\langle \boldsymbol{\zeta}^o (\boldsymbol{r}^-)^\top \right\rangle_{\mathcal{B}} - \eta_o \kappa_o U^o, \tag{S5.1}$$

$$\Delta B = \eta_B \left\langle \boldsymbol{r}^- (\boldsymbol{\zeta}^o)^\top \right\rangle_{\mathcal{B}} - \eta_B \kappa_B B. \tag{S5.2}$$

The activity products are transposes of each other, but each pathway changes its own weights using the activities of the connected neurons. Neither synapse reads the other synapse's weight. Part II writes the feedback matrix as $B_{\text{out}} = B^\top$, which has the same shape as $V = U^o$.

With matched scalar learning rates and decay, the mismatch $E = B - (U^o)^\top$ follows $E_{n+1} = (1 - \eta\kappa)E_n$ because the two activity increments cancel in the difference. When $|1 - \eta\kappa| < 1$, this mismatch contracts with successive updates. Centering, unequal learning rates and separate optimizer preconditioning change this relation, and the experiment-specific feedback alignment is therefore measured rather than assumed.

Centered FSN (cFSN) changes only the feedback-weight product. We subtract the minibatch mean of each neuron's free rate and output displacement:

$$\widetilde{\boldsymbol{r}}^{-} = \boldsymbol{r}^{-} - \left\langle \boldsymbol{r}^{-} \right\rangle_{\mathcal{B}}, \qquad \widetilde{\boldsymbol{\zeta}}^{o} = \boldsymbol{\zeta}^{o} - \left\langle \boldsymbol{\zeta}^{o} \right\rangle_{\mathcal{B}}. \tag{S5.3}$$

The centered Kolen–Pollack rule is

$$\Delta B = \eta_B \left\langle \widetilde{\boldsymbol{r}}^{-} (\widetilde{\boldsymbol{\zeta}}^{o})^{\top} \right\rangle_{\mathcal{B}} - \eta_B \kappa_B B. \tag{S5.4}$$

Its activity term is the within-batch cross-covariance. The means are taken across examples for each neuron, not across neurons. The directed input, recurrent and readout products in FSN are unchanged by this centering. Random-feedback FSN (rFSN) instead keeps $B$ fixed.

### S6 Contrastive controls

To test the role of the directed FSN product, the contrastive-Hebbian control uses the difference between recurrent coactivity in the two phases:

$$\Delta W_{\mathrm{CH}} = \frac{\eta_h}{\varepsilon_h} \left\langle \boldsymbol{r}^{+} (\boldsymbol{r}^{+})^{\top} - \boldsymbol{r}^{-} (\boldsymbol{r}^{-})^{\top} \right\rangle_{\mathcal{B}}. \tag{S6.1}$$

Substituting $\boldsymbol{r}^{+} = \boldsymbol{r}^{-} + \delta\boldsymbol{r}$ shows how this differs from FSN:

$$\boldsymbol{r}^{+} (\boldsymbol{r}^{+})^{\top} - \boldsymbol{r}^{-} (\boldsymbol{r}^{-})^{\top} = \delta\boldsymbol{r} (\boldsymbol{r}^{-})^{\top} + \boldsymbol{r}^{-} (\delta\boldsymbol{r})^{\top} + \delta\boldsymbol{r} (\delta\boldsymbol{r})^{\top}. \tag{S6.2}$$

The first term is the FSN product. The complete contrast is symmetric, whereas the FSN product distinguishes presynaptic free activity from postsynaptic displacement.

The MNIST and Reacher contrastive controls retain the delta-readout rule in Eq. (S4.6). The contrastive controls of the Ant experiment also replace the readout update by

$$\Delta U^{o}_{\mathrm{CH}} = \frac{\eta_o}{\varepsilon_o} \left\langle \boldsymbol{r}^{o,+} (\boldsymbol{r}^{+})^{\top} - \boldsymbol{r}^{o,-} (\boldsymbol{r}^{-})^{\top} \right\rangle_{\mathcal{B}}. \tag{S6.3}$$

The uncentered Kolen–Pollack increment uses the transposed activity product. For centered contrastive feedback, each population is centered separately within each phase:

$$\Delta B_{\mathrm{cCH}} = \frac{\eta_B}{\varepsilon_o} \left[ \left\langle \widetilde{\boldsymbol{r}}^{+} (\widetilde{\boldsymbol{r}}^{o,+})^{\top} \right\rangle_{\mathcal{B}} - \left\langle \widetilde{\boldsymbol{r}}^{-} (\widetilde{\boldsymbol{r}}^{o,-})^{\top} \right\rangle_{\mathcal{B}} \right] - \eta_B \kappa_B B. \tag{S6.4}$$

Reacher cCHFSN centers the recurrent contrastive product while retaining a delta readout and centered Kolen–Pollack mirroring. Ant cCHFSN centers both the recurrent and contrastive-readout products. Input-weight updates retain the directed displacement product. These task-specific controls are detailed in Sections S13–S15.

### S7 Algorithms and update schedules

Algorithm S1 collects the forward solve and implicit backward calculation. Algorithm S2 follows the two physical spiking phases and their local updates. The symbols and operations are defined in the preceding derivations; task-specific scaling, optimization and update schedules are given in Part II.

**Algorithm S1** CBIT with optional State-Carry Initialization

**Require:** A training batch, weights, input phases and iteration counts $K_m, J$

**Forward rate calculation**

**for** each input phase $m$ **do**

  Initialize $\boldsymbol{r}^{(0)}$ from the preceding phase for SCI, or the specified initial state.

  Detach the initializer from preceding phases.

  Apply $K_m$ damped fixed-point iterations using Eq. (S1.6).

  Store the resulting rate $\boldsymbol{r}_m$ for the readout and the next phase.

**end for**

Compute output rates and the task loss, including applicable rate penalties.

**Implicit credit calculation**

Compute output credit $\boldsymbol{p}^o = D^o \boldsymbol{\lambda}^o$ and the recurrent loss derivative $\boldsymbol{\lambda}^h$.

Initialize $\boldsymbol{a} \leftarrow \boldsymbol{\lambda}^h$.

Repeat $J$ times: $\boldsymbol{a} \leftarrow \boldsymbol{\lambda}^h + W^\top D \boldsymbol{a}$.

Set recurrent current-level credit $\boldsymbol{p}^h \leftarrow D\boldsymbol{a}$.

**Parameter update**

Form weight gradients from credit times presynaptic rate, Eqs. (S2.5), (S2.12).

Apply the configured optimizer, regularization and parameter constraints.

**Algorithm S2** Feedback-Spike Nudging with Kolen–Pollack mirroring

**Require:** Input spikes, an initial neural state, weights and an output teaching objective

**Free phase**

Present the input spikes and run the network without teaching feedback.

Record free rates $\boldsymbol{x}^-$, $\boldsymbol{r}^-$, $\boldsymbol{r}^{o,-}$ and the final membrane and spike states.

**Output teaching signal**

Obtain $\boldsymbol{\lambda}^o = \partial \mathcal{J} / \partial \boldsymbol{r}^{o,-}$ from the task objective.

Encode $-\varepsilon_o \boldsymbol{\lambda}^o$ as signed output-feedback events.

**Coupled feedback phase**

Continue from the free-phase final state; initialize recurrent teaching current to zero.

**for** each feedback timestep **do**

  Update recurrent neurons with sensory spikes and the available feedback current.

  Update output neurons with recurrent spikes and output-feedback events.

  **if** past the feedback burn-in **then**

    Update the running output rate and estimate $\widehat{\boldsymbol{p}}^o_t = -(\bar{\boldsymbol{r}}^{o,+}_{1:t} - \boldsymbol{r}^{o,-})/\varepsilon_o$.

    Encode the processed estimate as signed events $\boldsymbol{q}^o_t$.

    Send these through $B$: use $-\varepsilon_h B \boldsymbol{q}^o_t$ on the next timestep.

  **end if**

**end for**

Record feedback-phase rates and compute $\delta \boldsymbol{r} = \boldsymbol{r}^+ - \boldsymbol{r}^-$, $\delta \boldsymbol{r}^o = \boldsymbol{r}^{o,+} - \boldsymbol{r}^{o,-}$.

**Local plasticity**

Scale the measured displacements: $\boldsymbol{\zeta}^h \leftarrow \delta \boldsymbol{r}/\varepsilon_h$, $\boldsymbol{\zeta}^o \leftarrow \delta \boldsymbol{r}^o/\varepsilon_o$.

Input and recurrent increments: $\Delta U \leftarrow \eta_h \left\langle \boldsymbol{\zeta}^h (\boldsymbol{x}^-)^\top \right\rangle_{\mathcal{B}}$, $\Delta W \leftarrow \eta_h \left\langle \boldsymbol{\zeta}^h (\boldsymbol{r}^-)^\top \right\rangle_{\mathcal{B}}$.

Readout increment: $\Delta U^o \leftarrow \eta_o \left\langle \boldsymbol{\zeta}^o (\boldsymbol{r}^-)^\top \right\rangle_{\mathcal{B}}$.

Kolen–Pollack increment: $\Delta B \leftarrow \eta_B \left\langle \boldsymbol{r}^- (\boldsymbol{\zeta}^o)^\top \right\rangle_{\mathcal{B}}$.

For cFSN, center both factors in the $B$ product across the batch.

Add bias increments from the mean displacement and the configured decay terms.

Accumulate or apply these increments according to the task schedule (Part II).

### S8 Conditions and additional identities

#### S8.1 Finite-window balance and zero reset

Summing Eq. (S1.2) over a finite window of $T$ timesteps gives

$$\bar{h}_i = \theta_i \hat{r}_i + \beta_i \bar{v}_i + \frac{v_{T,i} - v_{0,i}}{T}, \qquad \hat{r}_i = \frac{1}{T} \sum_{t=0}^{T-1} s_{t+1,i}. \tag{S8.1}$$

The stationary identity drops the boundary term. The midpoint approximation to the remaining voltage average produces the transfer function; it is not an exact identity for arbitrary finite spike trains.

With zero reset, write $u_{t+1,i} = (1 - \beta_i) v_{t,i} + h_{t,i}$ and $v_{t+1,i} = u_{t+1,i}(1 - s_{t+1,i})$. The same summation gives

$$\bar{h}_i = \beta_i \bar{v}_i + \theta_i \hat{r}_i + \overline{(u_{t+1,i} - \theta_i) s_{t+1,i}} + \frac{v_{T,i} - v_{0,i}}{T}. \tag{S8.2}$$

The extra term measures threshold overshoot. The rate transfer remains an approximation for this reset rule. The reset used in each experiment is specified in Sections S9–S16.

##### S8.2 Finite forward and backward approximations

For an initializer in $[0,1]^N$ and $0 < \eta \leq 1$, the outer clipping in Eq. (S1.6) is redundant in exact arithmetic. Within a fixed branch, the iteration Jacobian is $J_\eta = (1-\eta)I + \eta DW$. Local convergence requires $\rho(J_\eta) < 1$, where $\rho$ denotes spectral radius. Invertibility of $I - DW$ by itself does not guarantee convergence of this iteration.

The infinite Neumann series converges if $\rho(W^\top D) < 1$. If $\boldsymbol{a}$ is the exact solution of Eq. (S2.13), the truncation error is

$$\boldsymbol{a} - \boldsymbol{a}^{(J)} = (W^\top D)^{J+1}\boldsymbol{a}. \tag{S8.3}$$

The implicit gradient is exact for a differentiable equilibrium branch. Evaluating it at a finite forward iterate approximates that equilibrium derivative; it generally differs from differentiating the finite iteration sequence itself.

##### S8.3 Gain and feedback-response identities

The gain can vanish for neurons clipped at zero or one. The identity

$$(I - DW^\top)D = D(I - W^\top D) \tag{S8.4}$$

establishes Eq. (S2.11) without requiring $D^{-1}$. At clipping boundaries the ordinary derivative is undefined; the chosen implementation convention is separate from the fixed-branch derivation.

For fixed-branch responses define $S = (I - DW)^{-1}D$ and $T = (I - DW^\top)^{-1}D$. Their difference satisfies

$$S - T = S(W - W^\top)T. \tag{S8.5}$$

For an output-driven loss and an exact isolated output credit, the feedback response differs from the implicit recurrent credit by

$$SB\boldsymbol{p}^o - T(U^o)^\top \boldsymbol{p}^o = (S - T)(U^o)^\top \boldsymbol{p}^o + S\big[B - (U^o)^\top\big]\boldsymbol{p}^o. \tag{S8.6}$$

The two terms separate recurrent asymmetry from mismatch in the feedback projection. These identities describe rate-model responses under the stated conditions; they do not guarantee descent for an arbitrary asymmetric, coupled or clipped finite-spike response.

##### S8.4 Stationary coupled feedback

For the ideal stationary version of the coupled episode, without signal clipping or normalization, substitute $\widehat{\boldsymbol{p}}^o = -\delta\boldsymbol{r}^o/\varepsilon_o$ into the recurrent feedback current. The recurrent phase relation and Eq. (S4.18) then give

$$(I - DW)\delta\boldsymbol{r} = (\varepsilon_h/\varepsilon_o)DB\delta\boldsymbol{r}^o, \tag{S8.7}$$

$$\delta\boldsymbol{r}^o = D^oU^o\delta\boldsymbol{r} - \varepsilon_o D^o\boldsymbol{\lambda}^o. \tag{S8.8}$$

Eliminating $\delta\boldsymbol{r}^o$ makes the additional feedback loop explicit:

$$\left[I - DW - \frac{\varepsilon_h}{\varepsilon_o}DBD^oU^o\right]\delta\boldsymbol{r} = -\varepsilon_h DBD^o\boldsymbol{\lambda}^o. \tag{S8.9}$$

This stationary calculation explains why the coupled displacement can differ from the isolated response. The experiments use finite spiking windows, running averages and the causal delay in Eq. (S4.17).

### Part II. Experimental methods

Sections S9–S16 describe the experimental protocols, architectures, parameters, comparator implementations, optimization schedules, evaluation procedures and statistical tests used in the manuscript. Supplementary Tables 1–10 are embedded in the corresponding sections. Unless stated otherwise, rates are expressed as spikes per internal simulation timestep, vectors are set in bold, every spike evaluation uses the trained parameters of the corresponding training condition, and random seeds are the experimental replicates. Paired comparisons use Wilcoxon signed-rank tests across matched seeds or conditions, with Holm correction [1] within the family stated for each analysis; the one-sided test of spine survival (Section S16) is the only directional test.

#### S9 Benchmarking theoretical equilibrium firing-rate estimates

##### S9.1 Benchmark design

Each candidate method was expressed as a scalar transfer function $\psi(h)$ from mean input current $h$ to firing rate. For recurrent weights $W$, input weights $W_{\text{in}}$, input rates $\boldsymbol{r}_{\text{in}}$, tonic current $b$ and recurrent rate vector $\boldsymbol{r}$, the theoretical equilibrium of each method was the self-consistent fixed point

$$\boldsymbol{r}^* = \psi\left(W\boldsymbol{r}^* + W_{\text{in}}\boldsymbol{r}_{\text{in}} + b\right),$$

with $\psi$ applied to each element. The fixed point $\boldsymbol{r}^*$ was compared with the empirical firing rate $\hat{\boldsymbol{r}}$ measured from a discrete-time recurrent LIF simulation (Methods) that used the same weights, input spikes and neuron parameters. Both zero-reset and subtractive-reset dynamics were evaluated. Empirical rates were computed after burn-in as

$$\hat{r}_i = \frac{1}{T - T_{\text{burn}}} \sum_{t=T_{\text{burn}}}^{T-1} s_{i,t},$$

with $T = 8000$ timesteps and $T_{\text{burn}} = 1500$.

##### S9.2 Benchmark parameter grid

The benchmark used $N_{\text{rec}} = 64$ recurrent neurons, $N_{\text{in}} = 16$ input neurons and threshold $\theta = 1.0$. The parameter grid crossed three random seeds, three membrane decay factors $\alpha$ (leak amount $\beta = 1 - \alpha$), two reset modes, three recurrent-weight scales and two input firing rates, giving 108 matched conditions:

**Supplementary Table 1 | Parameter grid for the recurrent LIF firing-rate benchmark.**

| Parameter | Values |
|---|---|
| Random seeds | 0, 1, 2 |
| Recurrent neurons | 64 |
| Input neurons | 16 |
| Simulation timesteps / burn-in | 8,000 / 1,500 |
| Membrane decay $\alpha$ | 0.85, 0.90, 0.95 |
| Leak amount $\beta = 1 - \alpha$ | 0.15, 0.10, 0.05 |
| Reset modes | zero, subtractive |
| Recurrent-weight scale | 0, 0.25, 0.55 |
| Input firing rate | 0.05, 0.15 |
| Input-weight scale | 0.85 |
| Threshold / tonic current | 1.0 / 0 |
| Number of matched conditions | 108 |

Input spikes were sampled independently as $s^{\text{in}}_{k,t} \sim \text{Bernoulli}(r_{\text{in},k})$, with the input firing probability held constant within a condition. Recurrent weights were sampled as

$$W = \frac{g_{\text{rec}}}{\sqrt{N_{\text{rec}}}} Z, \qquad Z_{ij} \sim \mathcal{N}(0,1),$$

where $g_{\text{rec}}$ is the recurrent-weight scale, with the diagonal set to zero. Input weights were sampled as

$$W_{\text{in}} = \frac{g_{\text{in}}}{\sqrt{N_{\text{in}}}} Z_{\text{in}}, \qquad Z_{\text{in},ik} \sim \mathcal{N}(0,1),$$

with $g_{\text{in}} = 0.85$. The tonic current was $b = 0$.

#### S9.3 Transfer functions

The current-balance rate equation (Section S1) was used in the form

$$\psi_{\text{CB}}(h) = \text{clip}\left(\frac{h}{\theta} - \frac{\beta}{2}, 0, 1\right).$$

Each comparator was evaluated by substituting its own transfer function $\psi_m(h)$ into the same recurrent fixed-point equation. The comparators are named by their functional form, with the source of that form in parentheses, as in Fig. 1c.

##### LIF steady state

With threshold current $q = \beta\theta$ and discrete-time membrane constant $\tau_m = -1/\log\alpha$, the deterministic LIF steady-state rate under constant drive was

$$\psi_{\text{LIF}}(h) = \frac{-\log\alpha}{-\log\left(1 - \frac{q}{h}\right)} \quad (h > q), \qquad \psi_{\text{LIF}}(h) = 0 \quad (h \le q).$$

Equivalent forms are given by Hunsberger and Eliasmith and by S2Act [2,3].

**Clipped linear (IDE-FSNN)**

$$\psi_{\text{IDE}}(h) = \text{clip}\left(\frac{h}{\theta}, 0, 1\right).$$

This is the clipped linear fixed-point rate used in implicit differentiation on the equilibrium state for feedback SNNs (IDE-FSNN) [4]; it omits the leak-offset term of the rate equation.

**Soft LIF (S2Act)**

The soft LIF rate of S2Act [3], which trains rate-based LIF actor–critic networks for spiking deployment, was mapped to the same discrete-current domain using $\Theta_\gamma(x) = \gamma \log\left(1 + e^{x/\gamma}\right)$ and

$$\psi_{\text{S2Act}}(h) = \left[\tau_{\text{ref}} + \tau_m \log\left(1 + \frac{q}{\Theta_\gamma(h - q)}\right)\right]^{-1},$$

with refractory period $\tau_{\text{ref}} = 0$ timesteps and $\gamma = 0.03$.

**Soft LIF (Hunsberger–Eliasmith)**

The softened LIF response of Hunsberger and Eliasmith [2] used the normalized current $J = h/q$, the soft threshold offset $z = \sigma \log\left(1 + e^{(J-1)/\sigma}\right)$ and

$$\psi_{\text{SoftLIF}}(h) = \left[\tau_{\text{ref}} + \tau_m \log\left(1 + \frac{1}{z}\right)\right]^{-1},$$

with $\tau_{\text{ref}} = 0$ timesteps and $\sigma = 0.03$.

**Threshold-aligned linear**

A control whose linear onset is aligned to the LIF threshold current, without the midpoint leak correction:

$$\psi_{\text{TA}}(h) = \text{clip}\left(\frac{h - q}{\theta - q}, 0, 1\right).$$

#### S9.4 Fixed-point solution

For each condition and transfer function $\psi_m$, the solver started from $\boldsymbol{r}^{(0)} = 0$ and applied damped iterations

$$\boldsymbol{r}^{(k+1)} = \text{clip}\left((1-\eta)\boldsymbol{r}^{(k)} + \eta\, \psi_m\Big(W\boldsymbol{r}^{(k)} + W_{\text{in}}\boldsymbol{r}_{\text{in}} + b\Big), 0, 1\right), \qquad \eta = 0.05,$$

until $\max_i |r_i^{(k+1)} - r_i^{(k)}| < 10^{-8}$ or for at most 2,500 iterations.

##### S9.5 Accuracy metric and statistical analysis

For each condition and method, the fixed point was compared with the empirical rate by the mean absolute error across the recurrent neurons,

$$\mathrm{MAE} = \frac{1}{N_{\mathrm{rec}}} \sum_i |r_i^* - \hat{r}_i|.$$

The statistical unit was the parameter condition ($n = 108$), not the neuron. For each comparator, the condition-level MAE of the rate equation was compared with that of the comparator by a Wilcoxon signed-rank test; negative paired differences favor the rate equation. Holm correction was applied over all 15 pairwise comparisons among the six transfer functions, of which Fig. 1c displays the five contrasts with the rate equation.

#### S10 MNIST classification with Current-Balance Implicit Training

##### S10.1 Dataset and training protocol

Each MNIST image [5] was represented as $I_n \in [0,1]^{1\times28\times28}$ and vectorized to $\boldsymbol{x}_n = \mathrm{vec}(I_n) \in [0,1]^{784}$. The comparison used the full 60,000-example training set and a 4,096-example test subset. Training used minibatches of 256 samples with shuffling, and evaluation used minibatches of 256 samples without shuffling. Each method was trained for exactly 3,000 optimizer steps, without early stopping. The same ten random seeds, 0–9, were used for all methods. All LIF populations used $\beta = 0.1813$: because the LIF update fixes only the ratio $\rho = \Delta t/\tau_m = -\ln(1-\beta) = 0.200$, a membrane time constant of $\tau_m = 20$ ms corresponds to a timestep of $\Delta t = 4.00$ ms, and the 100-timestep stimulus therefore lasts 400 ms and a per-timestep rate $r$ corresponds to $r/\Delta t$ spikes per second. Rasters and firing rates in the main figure use these physical units.

##### S10.2 RSNN architecture

All methods used the same LIF recurrent classifier: Bernoulli pixel spikes were passed through a single trainable linear projection into an input LIF population, input spikes drove a recurrent LIF population, and recurrent spikes drove a ten-neuron LIF output layer. The architecture used $N_{\mathrm{in}} = 192$ input, $N_{\mathrm{rec}} = 96$ recurrent and $N_{\mathrm{out}} = 10$ output LIF neurons. Let $A \in \mathbb{R}^{192\times784}$ be the sensory projection, $U \in \mathbb{R}^{96\times192}$ the input-to-recurrent matrix, $W \in \mathbb{R}^{96\times96}$ the recurrent matrix and $V \in \mathbb{R}^{10\times96}$ the recurrent-to-output matrix. Recurrent self-connections were set to zero after each optimizer step, and all trainable weights and biases were clipped to $[-3,3]$ after each update.

**Supplementary Table 2 | Architecture and initialization for the computational MNIST comparison.**

| Parameter | Value |
|---|---|
| Input LIF neurons | 192 |
| Recurrent LIF neurons | 96 |
| Output LIF neurons | 10 |
| Sensory front end | single trainable linear projection to the input LIF population |
| Simulation length | $T = 100$ |
| Burn-in | 10 timesteps |
| Input Bernoulli probability scale | 0.65 |
| Input current scale | 0.30 |
| Sensory projection weight scale | 0.70 |
| Sensory projection bias initialization | 0.240717 |
| Input-LIF bias initialization | 0.00 |
| Input-to-recurrent weight scale | 0.60 |
| Recurrent weight scale | 0.50 |
| Recurrent-to-output weight scale | 0.05 |
| Recurrent bias initialization | 0.122215 |
| Output bias initialization | 0.172215 |
| LIF threshold | 1.10 |
| LIF leak amount $\beta$ | 0.1813 (4.00-ms timestep at $\tau_m = 20$ ms) |
| Reset | subtractive |
| Recurrent-rate / output-rate $L_1$ penalty | 0.1 / 0.1 |

#### S10.3 Image spikes and spiking forward pass

For the spike-forward methods (BPTT, e-prop, ID-SNN and Rate-BP), for raster examples, empirical firing-rate measurements and all final spike evaluations, pixel values were converted into Bernoulli spike frames. At internal timestep $t$, pixel $j$ emitted

$$z_{n,t,j} \sim \text{Bernoulli}(\rho_{n,j}), \qquad \rho_{n,j} = \text{clip}(0.65\, x_{n,j}, 0, 1),$$

and the input-population current was $\boldsymbol{h}^{\text{in}}_{n,t} = 0.30\,(A\boldsymbol{z}_{n,t} + \boldsymbol{b}^{\text{in}})$, where $\boldsymbol{b}^{\text{in}}$ is the trainable bias of the sensory projection. All populations used subtractive-reset LIF neurons with $\beta = 0.1813$ and $\theta = 1.10$ (Methods). The recurrent and output currents were $\boldsymbol{h}^h_t = U\boldsymbol{s}^{\text{in}}_t + W\boldsymbol{s}^h_{t-1} + \boldsymbol{b}^h$ and $\boldsymbol{h}^o_t = V\boldsymbol{s}^h_t + \boldsymbol{b}^o$. The network was simulated for $T = 100$ timesteps and rates were computed after discarding the first ten timesteps. Class logits were $20\, r^o_k$, training minimized the cross-entropy loss, and spike predictions were $\hat{y} = \arg\max_k r^o_k$.

#### S10.4 CBIT implementation

CBIT replaced the spike trajectories by rate-equation states computed from the same synaptic currents (Sections S1 and S2); no recurrent spike trajectory was simulated during its training forward pass. The input rate used a two-sample Monte Carlo estimate of the Bernoulli sensory drive,

$$\boldsymbol{r}^{\text{in}}_n = \frac{1}{2}\sum_{m=1}^{2} \Psi\left(0.30\,(A\boldsymbol{z}^{(m)}_n + \boldsymbol{b}^{\text{in}})\right),$$

the recurrent fixed point $\boldsymbol{r}^{h,*} = \Psi(W\boldsymbol{r}^{h,*} + U\boldsymbol{r}^{\text{in}} + \boldsymbol{b}^h)$ was initialized at zero and solved with $K = 10$ damped iterations ($\eta = 0.5$, clipping to $[0, 1]$ after each iteration), and the output rate was $\boldsymbol{r}^{o,*} = \Psi(V\boldsymbol{r}^{h,*} + \boldsymbol{b}^o)$. Gradients were computed by implicit differentiation with 12 Neumann iterations without damping (Section S2).

##### S10.5 Comparator methods

**ID-SNN.** The spike-forward implicit-differentiation comparator first ran the stochastic spike simulation without surrogate gradients to obtain empirical input, recurrent and output rates. A rate-equation pass was then computed from the empirical input-LIF rates, with the recurrent state solved from zero, and the output used in the loss was

$$\boldsymbol{r}^{o}_{\text{used}} = \text{stopgrad}(\hat{\boldsymbol{r}}^{o}) + \boldsymbol{r}^{o,*} - \text{stopgrad}(\boldsymbol{r}^{o,*}),$$

with analogous straight-through constructions for internal rate quantities where required. ID-SNN therefore used the spike forward value while retaining the implicit rate-equation gradient. Its learning rate was $2.5 \times 10^{-3}$.

**BPTT.** BPTT simulated the full spiking network for $T = 100$ timesteps and differentiated through the unrolled computation graph using a fast-sigmoid surrogate derivative for the threshold [6],

$$\frac{\partial H(a)}{\partial a} \approx \frac{1}{(1+\kappa|a|)^2}, \qquad \kappa = 1,$$

with learning rate $5 \times 10^{-3}$.

**E-prop.** The e-prop comparator [7] kept a spiking LIF output layer rather than a non-spiking linear readout. Its gradient was computed by reverse-mode automatic differentiation of the spiking forward pass with the recurrent spike input detached where it enters the recurrent weights and with the reset not differentiated, the formulation that Bellec et al. used for their GPU experiments. This gradient is identical to the eligibility-trace form $\partial E/\partial W_{ji} = \sum_t L_j^t \, \bar{e}_{ji}^t$ (verified numerically to double precision against an explicit-trace implementation), but is obtained at a cost similar to BPTT. The pseudo-derivative was triangular,

$$\psi_{\text{ep}}(u-\theta) = \frac{\gamma_{\text{ep}}}{\theta} \max\left(0, 1 - \left|\frac{u-\theta}{w_{\text{ep}}\theta}\right|\right),$$

with $\gamma_{\text{ep}} = 0.3$ and $w_{\text{ep}} = 1.0$. Output spikes were decoded by an exponential moving average with decay 0.95, cross-entropy was applied at every post-burn-in timestep to 20 times the decoded trace, and the output error was broadcast to recurrent neurons through the current recurrent-to-output matrix $V$ (symmetric feedback). The sensory projection received the same automatic-differentiation gradient. The learning rate was $2 \times 10^{-3}$.

**Rate-BP.** Rate-based backpropagation was adapted from Yu et al. [8]. The spiking network was first run without gradient tracking to obtain post-burn-in spike rates and neuron-wise rate-gradient factors. For each LIF population, a sigmoid derivative around threshold,

$$\psi_{\text{RBP}}(u-\theta) = \alpha\,\sigma(\alpha(u-\theta))\,[1-\sigma(\alpha(u-\theta))], \qquad \alpha = 4.0,$$

was multiplied by a scalar eligibility factor accumulated through time as the membrane response to a current perturbation (detached reset), and these factors were averaged over time after burn-in. The backward graph was then compressed into a single rate-level spatial graph: if $\hat{\boldsymbol{r}}^{\ell}$ is the empirical spike rate of layer $\ell$, $\boldsymbol{c}^{\ell}$ the mean current computed from upstream rates and trainable weights, and $\bar{\boldsymbol{\psi}}^{\ell}_{\text{RBP}}$ the averaged factor, the graph used

$$\boldsymbol{r}^{\ell}_{\text{RBP}} = \text{stopgrad}(\hat{\boldsymbol{r}}^{\ell}) + \boldsymbol{c}^{\ell} \odot \bar{\boldsymbol{\psi}}^{\ell}_{\text{RBP}} - \text{stopgrad}(\boldsymbol{c}^{\ell} \odot \bar{\boldsymbol{\psi}}^{\ell}_{\text{RBP}}).$$

For the sensory projection, the rate graph used the Bernoulli image probabilities as the input-rate estimate. No additional $1/T$ scaling of the loss gradient was applied. The learning rate was $2 \times 10^{-3}$.

##### S10.6 Optimization, evaluation and statistical analysis

Every method used Adam [9] with the learning rates given above (CBIT, $2.5 \times 10^{-3}$), weight decay $10^{-5}$, global gradient clipping at 1.0, zero recurrent diagonal and post-update clipping of trainable weights and biases to $[-3, 3]$. Recurrent-rate and output-rate $L_1$ regularization coefficients were both 0.1, applied to the recurrent and output firing rates of each method's training forward pass (rate-equation values for CBIT, empirical spike values for the spike-forward methods).

All reported accuracies were measured by spike simulation for $T = 100$ timesteps with Bernoulli image spikes and hard thresholds. The final evaluation used 16 minibatches of 256 samples (4,096 examples); the same subset was used for periodic and final evaluation. Training-only time included optimizer-step computation and excluded evaluation, plotting and output writing. Peak CUDA memory was the maximum allocated CUDA memory during training.

The example rasters (Fig. 1e) used the CBIT model of seed 5 and are plotted in physical time (4 ms per timestep) with output rates in spikes per second; they show all 192 input, 96 recurrent and 10 output neurons over all 100 timesteps without cropping, removal of silent rows or sorting.

The statistical unit was the random seed, and all comparisons were paired by seed. For each metric (best evaluated accuracy, final evaluated accuracy, training-only time and mean recurrent firing rate at evaluation), CBIT was compared with each of the four alternatives by an exact Wilcoxon signed-rank test, with Holm correction across these four comparisons. Accuracy differences are reported in percentage points.

A separate simulation-length analysis used seed 7 and varied $T \in \{5, 10, 30, 50, 100, 150, 300, 500\}$ timesteps (20 to 2,000 ms of simulated time) while keeping the architecture and training configuration fixed, except that the burn-in was $\min(10, \lfloor T/10 \rfloor)$ timesteps, recording training-only time and peak CUDA memory for every method. A second single-seed sweep (seed 7) varied the number of recurrent neurons over 10, 30, 100, 300, 1,000, 3,000, 10,000 and 30,000 with $T = 100$ and all other settings, including the learning rates, held at the values of the 96-neuron configuration; only the training-only time of each run is reported (Fig. 1i).

#### S11 Modular addition with State-Carry Initialization

##### S11.1 Sequential modular-addition task

A sequential version of the modular-addition task [10,11] was used. Each trial consisted of four sequential input phases: an inter-cue signal (ICS), a first digit, a second ICS and a second digit. The ICS was carried by a single cue channel and signaled addition in both ICS phases. The digits were $x, y \in \{0, \ldots, 9\}$, and the supervised target after the second digit was $(x + y) \bmod 10$. The training loss was applied to the second digit phase only, the first phase at which the sum is defined. The input space contained 11 channels: the cue channel and ten digit channels. Training batches were balanced across ordered digit pairs.

##### S11.2 Input-rate representation and spike generation

For each phase $m$, the symbolic event was represented as a rate vector $\boldsymbol{q}_m \in [0,1]^{11}$. Digit channels used a rate of 0.70 for the active digit and 0.03 for inactive digit channels; the cue channel used 0.70 during ICS phases and 0 otherwise. During training, Gaussian noise with standard deviation 0.02 was added to the phase-rate vector before clipping to $[0,1]$. For explicit-spike evaluation, the phase-rate vector was converted to Bernoulli input spikes at each internal timestep, $z_{m,t,j} \sim \text{Bernoulli}(q_{m,j})$.

##### S11.3 RSNN architecture

The model was a spiking recurrent network with one recurrent LIF layer of 1,024 neurons and one LIF output layer of 10 neurons, with input-to-recurrent matrix $U \in \mathbb{R}^{1024\times 11}$, recurrent matrix $W \in \mathbb{R}^{1024\times 1024}$ and recurrent-to-output matrix $V \in \mathbb{R}^{10\times 1024}$. Both layers used subtractive-reset LIF dynamics (Methods) with threshold $\theta = 1.0$, $\beta_r = 0.005$ for the recurrent layer and $\beta_o = 0.02$ for the output layer. In explicit-spike simulation, the recurrent and output membrane states were carried across phases, so that the spiking evaluation preserved temporal continuity across the full trial.

**Supplementary Table 3 | Architecture and initialization for sequential modular addition.**

| Parameter | Value |
|---|---|
| Input channels | 11 |
| Recurrent LIF neurons | 1024 |
| Output LIF neurons | 10 |
| Recurrent leak $\beta_r$ | 0.005 |
| Output leak $\beta_o$ | 0.02 |
| Threshold | 1.0 |
| Reset | subtractive |
| Input weight scale | 0.60 |
| Recurrent weight scale before spectral normalization | 0.45 |
| Output weight scale | 0.45 |
| Recurrent spectral radius at initialization | 0.95 |
| Recurrent bias initialization | 0.01 |
| Output bias initialization | 0.20 |
| Output state carried across phases | yes |

##### S11.4 Training with SCI: phase durations and solver schedule

Training used CBIT with SCI (Sections S2 and S3): the recurrent rate of each phase was solved from the fixed point $\boldsymbol{r}_m = \Psi(W\boldsymbol{r}_m + U\boldsymbol{q}_m + \boldsymbol{b}^r)$ initialized at the preceding phase's solved rate, gradients were not propagated through the carried initializer, and the output rate $\boldsymbol{r}^o_m = \Psi(V\boldsymbol{r}_m + \boldsymbol{b}^o)$ gave the logits $20\,\boldsymbol{r}^o_m$.

During training, the ICS phases lasted 30 timesteps (8-step burn-in for explicit-spike diagnostics), the first digit phase 240 timesteps (80-step burn-in) and the second digit phase 300 timesteps (120-step burn-in). For the explicit-spike evaluation and the rate-alignment analysis reported in the Results and Extended Data Fig. 1d, the ICS phases lasted 40 timesteps with a 10-step burn-in, and the digit phases were unchanged. The principal-component visualization used the training phase durations.

The fixed-point solver used phase-specific iteration counts. During the first training stage, ICS phases used 2 iterations, the first digit phase 40 and the second digit phase 20. After step 400, training entered a lower-learning-rate stage in which the digit phases used 60 and 40 iterations;

ICS phases continued to use 2. The relaxation factor was 0.05 for ICS phases and 0.20 for digit phases. Gradients through the recurrent fixed point used a 16-step Neumann approximation with the hard clipped transfer function.

##### S11.5 Objective and optimization

The primary loss was the cross-entropy of the second-digit logits $20\,\boldsymbol{r}^o$ with target class $(x + y) \bmod 10$. Two output-shaping terms were added: a rate mean-squared-error term with weight 0.30, using target rate 0.80 for the correct output neuron and 0.005 for the others, and a margin loss with weight 0.70 that encouraged the target output rate to exceed the runner-up by at least 0.45. A recurrent firing-rate $L_1$ penalty with weight 0.03 and weight decay $10^{-5}$ were used.

The optimizer was Adam with initial learning rate $2.5 \times 10^{-4}$, gradient-norm clipping at 1.0 and a second stage from training step 400 with learning rate $3 \times 10^{-6}$. Each seed was trained for 900 optimizer steps with batch size 512. The same ten seeds, 0–9, were used for all endpoint summaries, and explicit-spike performance was evaluated with the final trained models.

##### S11.6 Explicit-spike evaluation and rate-alignment analysis

Explicit-spike performance was measured with Bernoulli input spikes and hard LIF recurrent and output dynamics. Each of the ten trained models was evaluated on eight batches of 1,024 examples (8,192 per seed) with 40-timestep ICS phases (10-step burn-in); membrane and spike states evolved continuously through all four phases, and the predicted class was the output neuron with the largest post-burn-in firing rate during the second digit.

To test whether SCI rate states predicted the spiking trajectory, theoretical recurrent and output firing rates were compared with empirical spike rates from explicit simulations, using the final model of seed 1, all 100 ordered digit pairs and six spiking trials per pair with independent input-rate noise, simulated with the phase durations of the accuracy evaluation. The theoretical recurrent rate was the rate-equation state of the current phase initialized from the previous phase state; empirical rates were the mean spikes after burn-in. Alignment was summarized with $R^2$ separately for the first and second digit phases and for the recurrent and output populations. Pair-grid evaluation tested all 100 ordered digit pairs: explicit-spike accuracy was averaged over repeated stochastic trials for each pair, and the resulting $10 \times 10$ grid was averaged across seeds.

##### S11.7 Principal-component visualization

Recurrent spiking activity was projected onto the first two principal components of recurrent activity. For each shown digit pair, 20 explicit-spike trials with independent input-rate noise were simulated, and the cumulative mean spike rate within each digit phase (binned in non-overlapping 10-step windows) was projected into the component space as a trajectory. The trial-averaged cumulative-rate trajectories of the first- and second-digit phases were plotted together with the projections of the theoretical rate states, namely the state solved after the first digit and the state solved with SCI after the second digit. The principal axes were fitted to a separate simulation with six spiking trials per ordered pair, in which the recurrent spikes of the whole trial were binned in the same 10-step windows, and were held fixed.

##### S11.8 Solver comparison for the carried state

The solver comparison used the same ten trained models. Each theoretical solver was evaluated on all 100 ordered digit pairs with six input-rate-noise trials per pair (600 examples per model);

the predicted class was the second-digit output-rate argmax; weights were fixed and no solver received labels. ICS phases used two iterations with relaxation 0.05, the first digit 60 iterations and the second digit 40, both with relaxation 0.20.

Four solvers were compared. SCI started the first ICS phase from zero and each subsequent phase from the previous theoretical recurrent rate. Zero initialization started every phase from zero. Random initialization started every phase from an independent Uniform(0,1) vector; each model-level accuracy pooled 16 restarts, with the same restart seeds paired across model seeds. The clipped closed-form solution solved the unclipped linear recurrent system in double precision, clipped the resulting recurrent vector to $[0, 1]$ and computed the clipped output rate from that vector; post-solve clipping does not in general solve the clipped recurrent equation, because clipping changes the recurrent input. Model-level accuracies are summarized across the ten seeds by their mean and population standard deviation.

### S12 Continuous-control reinforcement learning with CBIT

#### S12.1 Tasks and comparison design

Continuous-control experiments used the Gymnasium MuJoCo locomotion environments Ant-v5 and Walker2d-v5 [12,13]. Agents were trained with proximal policy optimization (PPO) [14] and generalized advantage estimation (GAE) [15]. All methods used the same recurrent spiking actor–critic architecture, observation preprocessing, environment timing, PPO rollout structure, reward normalization, evaluation protocol and seed set; the experimental variable was the training rule used to obtain gradients through the recurrent spiking network:

1. **CBIT with State-Carry Initialization (CBIT+SCI)**: the recurrent state was represented by a current-balance fixed point during training, initialized from the previous solved recurrent state, and gradients through that fixed point were computed by implicit differentiation.
2. **ID-SNN**: the numerical forward loss used the actual spiking trajectory, and gradients were routed through the current-balance equilibrium model by a straight-through construction.
3. **BPTT**: the forward pass used the actual spiking trajectory and gradients were computed by surrogate-gradient backpropagation through the internal spike timesteps.

Learning rates and the other task-specific parameters are listed in Supplementary Table 4.

#### S12.2 Observation preprocessing and spike encoding

At each environment step, the raw observation vector $\boldsymbol{o}_t$ was normalized online using a running mean and variance, then clipped:

$$\tilde{\boldsymbol{o}}_t = \mathrm{clip}\left(\frac{\boldsymbol{o}_t - \boldsymbol{\mu}_o}{\sqrt{\boldsymbol{\sigma}_o^2 + \epsilon}}, -10, 10\right).$$

Because MuJoCo observations can be positive or negative, each normalized observation dimension was split into positive and negative nonnegative channels,

$$\boldsymbol{x}_t^+ = \mathrm{ReLU}\left(\mathrm{clip}(\tilde{\boldsymbol{o}}_t/3, -1, 1)\right), \qquad \boldsymbol{x}_t^- = \mathrm{ReLU}\left(-\,\mathrm{clip}(\tilde{\boldsymbol{o}}_t/3, -1, 1)\right), \qquad \boldsymbol{x}_t = [\boldsymbol{x}_t^+, \boldsymbol{x}_t^-],$$

so that an observation of dimension $N_o$ was encoded as $2N_o$ nonnegative input channels. During spiking forward passes, each channel was sampled independently at each internal timestep, $z_{t,\tau,k} \sim \mathrm{Bernoulli}\,(\mathrm{clip}(x_{t,k}, 0, 1))$. BPTT and ID-SNN used these sampled input spikes; CBIT+SCI used the expected input rate. When empirical input spikes had already been generated for the spike-forward control, their mean rate was used in the equilibrium backward path.

##### S12.3 Alignment of RSNN time with environment time

If one environment action covered duration $\Delta t_{\mathrm{env}}$ and the RSNN timestep was $\Delta t_{\mathrm{SNN}}$, the number of internal timesteps per decision was $T_{\mathrm{SNN}} = \mathrm{round}(\Delta t_{\mathrm{env}}/\Delta t_{\mathrm{SNN}})$, and the LIF decay followed from the membrane time constant, $\alpha = \exp(-\Delta t_{\mathrm{SNN}}/\tau_m)$, $\beta = 1 - \alpha$. Recurrent and readout LIF neurons used $\tau_m = 20\,\mathrm{ms}$. Ant-v5 used frame skip 5, $\Delta t_{\mathrm{SNN}} = 0.32\,\mathrm{ms}$ and 156 timesteps per decision; Walker2d-v5 used frame skip 8, $\Delta t_{\mathrm{SNN}} = 0.08\,\mathrm{ms}$ and 200 timesteps per decision.

##### S12.4 Recurrent spiking actor–critic model

The policy and value function shared a recurrent LIF encoder with 128 neurons, which received the input projection $U\boldsymbol{z}_{t,\tau}$, the recurrent input $W\boldsymbol{s}^h_{t,\tau}$, a trainable bias $\boldsymbol{b}^h$ and a tonic current $I_0 = 0.05$. Neurons used threshold $\theta = 1$ and subtractive reset (Methods). The recurrent rate feature of one environment step was the mean recurrent spike rate over the internal window. For spiking methods, recurrent membrane and spike states were carried across environment steps within an episode, detached between steps and reset at episode boundaries.

Separate actor and critic heads read out from the shared recurrent population through spiking population readouts of 16 positive-coding and 16 negative-coding LIF neurons per scalar output, decoded as

$$y_j = a_j(r_j^+ - r_j^-),$$

where $a_j$ is a learnable positive scale initialized at 3.0 (Ant-v5 actor), 2.5 (Walker2d-v5 actor) or 10.0 (critic). The actor output parameterized the mean of a tanh-squashed diagonal Gaussian policy; evaluation used the policy mean.

##### S12.5 Current-balance equilibrium forward pass

CBIT+SCI replaced the recurrent spike trajectory during training by the current-balance equilibrium $\boldsymbol{r}_t^\star = \Psi(W\boldsymbol{r}_t^\star + U\boldsymbol{x}_t + \boldsymbol{b}^h + I_0)$, where $\boldsymbol{x}_t$ is the observation-rate code or the empirical input-spike rate, and applied the same transfer function to the actor and critic readouts. The rate state was computed with 10 damped iterations, relaxation $\eta = 0.25$ and clipping to $[0, 1]$. The main condition initialized the solve from the previous solved state, $\boldsymbol{r}_t^{(0)} = \boldsymbol{r}_{t-1}^\star$, separately for each parallel environment; the stored state was detached after each environment step and reset when that environment terminated. Two initialization controls started the fixed point at every environment step from zero or from a clipped closed-form solution (unclipped linear solve with diagonal damping $10^{-4}$, then clipping), followed by the same ten damped iterations. The implicit gradient used 5 Neumann iterations (Section S2).

##### S12.6 ID-SNN and BPTT controls

During the PPO update, ID-SNN first ran the full spiking encoder and readout forward path without gradient tracking to compute the numerical action log-probabilities and values. The empirical recurrent spike rate was supplied directly as the recurrent state for implicit differentiation, without a second fixed-point solve; the backward path used the mean of the sampled input spikes and the rate-model readouts, joined by the straight-through expression

$$\boldsymbol{y} = \boldsymbol{y}_{\text{spike}}^{\text{detach}} + \boldsymbol{y}_{\text{eq}} - \boldsymbol{y}_{\text{eq}}^{\text{detach}},$$

so that the forward value came from spikes and the gradient from the equilibrium model. BPTT used the same spiking forward path and differentiated through the internal spike-time trajectory with a surrogate threshold derivative [6].

##### S12.7 PPO optimization, evaluation and statistical analysis

Each PPO update collected 64 environment steps from 64 parallel environments (4,096 transitions) and used 4 minibatches and 2 epochs, policy clipping coefficient 0.2, discount factor $\gamma = 0.99$, GAE parameter $\lambda = 0.95$, target KL 0.03, maximum gradient norm 1, reward normalization with reward clipping at 10, spike-rate regularization coefficient 0.001 and entropy coefficient 0. The value-loss coefficient was 1.3 for Ant-v5 and 1.0 for Walker2d-v5. Each condition was trained on ten matched seeds (0–9) for 2,441 full rollouts (9,998,336 transitions), the largest whole number of rollouts within 10 million environment steps, with linear learning-rate annealing. Evaluation occurred every ten updates and at the final update. Recurrent state was maintained across rollout boundaries and reset at episode termination.

All evaluation rewards (total reward per episode) were measured with the spiking RSNN forward path, including for CBIT+SCI, over 64 completed deterministic episodes. Learning curves show the seed mean with SEM shading. Because the same seeds were used for every condition, comparisons were paired by seed. For each environment, best and final evaluation rewards were separate families, each containing the four comparisons of CBIT+SCI with the controls (ID-SNN, BPTT, zero initialization and closed-form initialization), with Holm correction.

##### S12.8 Conditional gradient variance under input-spike resampling

For Extended Data Fig. 2f, BPTT, ID-SNN and CBIT+SCI were compared at the same initial parameters, using the initial networks of seeds 0–9 in Ant-v5 and Walker2d-v5. Each network supplied one fixed minibatch of 1,024 transitions from its first spiking-policy rollout (64 environments $\times$ 64 decisions); the normalized observations, actions, old log probabilities, values, advantages (normalized once within the minibatch), return targets and incoming network states were held fixed. We generated 32 independent Bernoulli input-spike realizations per network from the fixed observation-dependent probabilities. BPTT and ID-SNN received the same input spikes and incoming spiking states within each realization; CBIT+SCI used the expected input probabilities and a fixed incoming equilibrium state from the same observation history. All incoming states were detached. Ant used 156 internal timesteps and Walker2d 200, with 128 recurrent neurons; CBIT used ten damped forward iterations with relaxation 0.25 and five Neumann iterations; ID-SNN used the empirical recurrent spike rate as its backward state with rate-model readout derivatives.

For each realization, gradients of the clipped PPO policy loss alone and of the full PPO objective (value loss and configured regularization included, with the training coefficients) were extracted before gradient-norm clipping and without an optimizer step, concatenating all model parameters except the Gaussian log standard deviation in a fixed order. For gradient vectors $\boldsymbol{g}_r$ over $R = 32$ repetitions,

$$\bar{\boldsymbol{g}} = \frac{1}{R}\sum_{r=1}^{R}\boldsymbol{g}_r, \qquad V = \frac{1}{R-1}\sum_{r=1}^{R}\|\boldsymbol{g}_r - \bar{\boldsymbol{g}}\|_2^2, \qquad V_{\mathrm{rel}} = \frac{V}{\|\bar{\boldsymbol{g}}\|_2^2}.$$

$V$ is the trace of the sample gradient covariance, and the relative variance normalizes it by the squared mean-gradient norm. Each initial-network seed supplied one estimate from its 32 repetitions; CBIT+SCI's gradients were identical across repetitions because its expected inputs and incoming states were fixed, giving zero conditional variance. Panel f shows policy-only relative variance with one point per seed and arithmetic means with 95% percentile confidence intervals from 10,000 bootstrap resamples of the ten seeds. The percentage reductions in the Results are $100(1 - \text{ratio})$, where the ratio is the geometric mean over seeds of the within-seed ratio of ID-SNN to BPTT relative variance.

Exact Wilcoxon signed-rank tests compared ID-SNN and CBIT+SCI with BPTT on seed-level relative variance, with Holm correction over the eight comparisons (two methods $\times$ two tasks $\times$ two objectives). The 32 spike draws estimate the variance within a seed and were not treated as independent networks; bootstrap intervals are descriptive. This comparison measures sensitivity to the input-spike realization of the current decision at initialization; it does not estimate gradient variability across minibatches, rollouts or training checkpoints.

##### S12.9 Tracking the rate state during spiking control

For Fig. 2d and Extended Data Fig. 2c, the final CBIT+SCI checkpoint of each of the ten seeds was evaluated over eight episodes per environment (up to 1,000 steps) with reproducibly seeded Bernoulli input spikes and deterministic policy-mean actions. The controller carried its spiking state, and three rate estimates received the same observation and weights at every step: SCI initialized from its own preceding rate estimate, zero initialization, and the clipped closed-form solution without iterative refinement. The first two used the training solver with ten iterations and relaxation 0.25, with expected observation rates as input. No labels, future observations or measured recurrent rates were supplied to SCI. Recurrent MAE was computed across the 128 neurons at each environment step and averaged across the episodes still active at that step within each seed; seed curves were smoothed with a centered 21-step running mean (partial windows at the edges) and summarized by mean and SEM.

##### S12.10 Development of touchdown-aligned activity during locomotor learning

For Fig. 2e,f and Extended Data Fig. 2d,e, the ten CBIT+SCI training seeds of each task were replayed at checkpoints corresponding to 0%, 1%, 2%, 5%, 10%, 15%, 20%, 30%, 40%, 50%, 60%, 70%, 80%, 90% and 100% of training (nearest retained checkpoint), with eight explicit-spike episodes per checkpoint. The sequentiality time course (Fig. 2f and Extended Data Fig. 2e) uses all of these checkpoints, whereas the activity heatmaps, limb positions and controls (Fig. 2e and Extended Data Fig. 2d) display the 5%, 30% and 100% checkpoints, which contained 503,808, 3,002,368 and 9,998,336 environment transitions. These replays are separate from the 64-episode evaluations used for the performance comparisons.

Replay used each checkpoint's frozen observation normalizer, policy-mean actions and stochastic sensory spikes, with recurrent spiking state carried between decisions and reset on episode termination. The internal neural exposure per decision was $156 \times 0.32$ ms $= 49.92$ ms for Ant and

$200 \times 0.08$ ms = 16 ms for Walker2d, and the physical decision intervals were 50 ms and 16 ms. The same environment-reset and spike-generator random seeds were reused across checkpoints.

**Contact events and episode sets.** A limb was in contact when any of its geometries contacted the ground, detected at the MuJoCo physics resolution (10 ms for Ant, 2 ms for Walker2d). A contact-state change had to persist for at least 20 ms and two physics samples; its timestamp was the beginning of the qualifying run. Touchdowns in the first second were excluded. Accepted touchdown-to-touchdown intervals lasted at least three decision steps and at most 3 s, and a central touchdown required accepted preceding and following intervals and at least 0.7 s of trajectory on both sides. At each checkpoint, episodes 0–3 formed the sorting set and episodes 4–7 the held-out set; sets were not pooled across checkpoints, and episodes that failed the inclusion criteria were excluded rather than reassigned. For each training seed, the limb with the most accepted events in the final-checkpoint sorting episodes was the reference limb and was retained across checkpoints. An episode required at least two matched events and nonzero kernel exposure throughout the analysis window; a seed/checkpoint required at least two eligible episodes and at least ten matched events in each episode set. Checkpoints failing these criteria were missing, not zero-valued, observations.

**Event-aligned templates.** Activity was analyzed from $-0.5$ to $+0.5$ s around touchdown on a grid of 200 cells with 5-ms spacing. Decision-level spike counts were assigned their decision-start timestamps and smoothed with a Gaussian kernel of standard deviation 25 ms, truncated at four standard deviations. With $C_{e,d,i}$ the spike count of neuron $i$ in decision $d$ of episode $e$, $u_{e,d}$ the decision-start time, $b_{e,k}$ the retained touchdown times and $h$ the internal neural exposure per decision in seconds,

$$w_{e,t,d} = \sum_k K_{25\,\mathrm{ms}}(u_{e,d} - b_{e,k} - t), \qquad R_{e,i}(t) = \frac{\sum_d w_{e,t,d} C_{e,d,i}}{h \sum_d w_{e,t,d}},$$

so that rates are weighted spike counts divided by weighted internal neural exposure, in internal spikes per second; event weights add when windows overlap, and eligible episode templates were averaged with equal weight. For the heatmaps, all 128 neurons were ordered at each checkpoint by the time of their maximum in that checkpoint's mean sorting template (first maximum, stable ordering); the order was fitted independently for each displayed checkpoint. A neuron's reference mean and population SD were calculated across time and eligible sorting-episode templates of the same checkpoint; held-out rates minus this mean were divided by the larger of the reference SD and 1 internal spike per second, averaged over eligible held-out episodes, and clipped to $\pm 3$ for display only.

Limb position was the distal capsule endpoint relative to the torso origin, rotated into torso coordinates; the displayed coordinate is forward position in centimeters. Positions and applied controls used the same decision timestamps and event weights as firing rates, without the exposure divisor; position lines and bands show the mean and sample SD across held-out episode mean trajectories, and control lines show the mean normalized applied action. The displayed Ant model (training seed 2) used right hindlimb contact and four held-out episodes at each displayed checkpoint (165, 295 and 317 matched events); the Walker2d model (training seed 3) used left foot contact and four held-out episodes (25, 123 and 163 events).

**Sequentiality index.** The entropy-based sequentiality index (SqI) of Zhou, Masmanidis and Buonomano [16] was computed from the held-out episode rate templates averaged into a non-negative raw-rate matrix $A_i(t)$ ($N = 128$ neurons, $T = 200$ time samples), without z-scoring or clipping. If $p_b$ is the proportion of neurons whose peak occurs in histogram bin $b$ (ten bin centers equally spaced from 0 to $T$, midpoint boundaries, first maximum for ties, as in ref. [16]), the normalized peak entropy is $\mathrm{PE} = -\sum_{b=1}^{10} p_b \log_2 p_b / \log_2 10$. With $q_i(t) = A_i(t) / \sum_j A_j(t)$,

$$\mathrm{TS} = 1 - \frac{1}{T\log_2 N}\sum_t\left[-\sum_i q_i(t)\log_2 q_i(t)\right], \qquad \mathrm{SqI} = \sqrt{\mathrm{PE}\,\mathrm{TS}}.$$

Terms with zero probability contribute zero to the entropy, and a time sample with zero population activity makes the index undefined. SqI was computed separately for each eligible seed/checkpoint and summarized by the seed mean and SEM; no hypothesis test was used for these descriptive trajectories. Eligible seed counts were:

| Training progress (%) | 0 | 1 | 2 | 5 | 10 | 15 | 20 | 30 | 40 | 50 | 60 | 70 | 80 | 90 | 100 |
|---|---|---|---|---|---|---|---|---|---|---|---|---|---|---|---|
| Ant | 5 | 9 | 10 | 10 | 10 | 10 | 10 | 10 | 10 | 10 | 10 | 10 | 10 | 10 | 10 |
| Walker2d | 0 | 0 | 0 | 7 | 9 | 7 | 9 | 9 | 8 | 8 | 7 | 7 | 9 | 9 | 10 |

##### S12.11 Hyperparameters

**Supplementary Table 4 | Task-specific parameters for computational continuous-control reinforcement learning.**

| Parameter | Ant-v5 | Walker2d-v5 |
|---|---|---|
| CBIT+SCI / ID-SNN learning rate | 0.001 | 0.00075 |
| BPTT learning rate | 0.005 | 0.003 |
| Frame skip | 5 | 8 |
| SNN timestep | 0.32 ms | 0.08 ms |
| SNN/readout steps | 156 / 156 | 200 / 200 |
| LIF $\tau_m$ | 20 ms | 20 ms |
| Recurrent neurons | 128 | 128 |
| Readout population | 16 positive + 16 negative per scalar | 16 positive + 16 negative per scalar |
| PPO rollout | 64 environments × 64 steps | 64 environments × 64 steps |
| Value-loss coefficient | 1.3 | 1.0 |

#### S13 Biologically local MNIST learning with Feedback-Spike Nudging

##### S13.1 MNIST dataset and spike input

The full 60,000-image MNIST training set [5] was used for training and the 10,000-image test set for evaluation. Each image was normalized to pixel intensities in $[0, 1]$ and represented as 784 sensory channels. For each image and internal timestep $t$, pixel $k$ emitted a Bernoulli input spike $x_{t,k} \sim \mathrm{Bernoulli}(I_k)$, where $I_k$ is the normalized intensity. For the methods with a feedback phase, the same input spike sequence was reused in that phase, so that the measured difference between phases reflects the feedback perturbation rather than resampling noise.

##### S13.2 RSNN architecture and parameters

All methods trained with local rules used the same spiking architecture: 784 input channels, a 256-neuron recurrent LIF layer and a 10-neuron LIF output layer. The trainable forward parameters were the input-to-recurrent matrix $U$, recurrent matrix $W$, recurrent-to-output matrix $V$, recurrent bias $\boldsymbol{b}$ and output bias $\boldsymbol{b}^o$; the output-to-recurrent feedback matrix $B_{\mathrm{out}}$ transmitted output-derived feedback spikes to the recurrent population. The recurrent current was

$\boldsymbol{h}_t = U\boldsymbol{x}_t + W\boldsymbol{s}_{t-1} + \boldsymbol{b} + \boldsymbol{f}_t^h$ and the output current $\boldsymbol{h}_t^o = V\boldsymbol{s}_t + \boldsymbol{b}^o + \boldsymbol{f}_t^o$, where the feedback currents $\boldsymbol{f}_t^h$ and $\boldsymbol{f}_t^o$ are zero in the free phase. Recurrent and output neurons used subtractive-reset LIF dynamics (Methods) with $\beta = 0.05$ and $\theta = 1.0$. The recurrent and output biases were initialized to 0.22 and 0.16; input, recurrent, output and output-feedback weights were initialized with scales 0.70, 0.08, 0.70 and 0.70. Recurrent self-connections were removed, and all trainable weights were clipped to maximum absolute value 3.0.

##### S13.3 Free phase, output error and feedback phase

Each training step used a free phase of $T_{\text{free}} = 100$ timesteps followed by a feedback phase of $T_{\text{nudge}} = 100$ timesteps; the first 10 timesteps of each phase were discarded as burn-in when computing rates. Classification logits were $\boldsymbol{z} = 20\,\boldsymbol{r}^{o,-}$, the loss was the cross-entropy, and for class label $y$ the output teaching signal was

$$\boldsymbol{\lambda}^o = \frac{\partial \mathcal{L}}{\partial \boldsymbol{r}^{o,-}} = 20\left[\text{softmax}(20\boldsymbol{r}^{o,-}) - \text{onehot}(y)\right].$$

This signal was converted into signed Bernoulli feedback events (Section S4) that perturbed the output layer first; the resulting output displacement was converted into output-credit feedback spikes that projected to the recurrent layer through $B_{\text{out}}$. The nudge strength was $\epsilon = 0.15$, with output feedback gain 2.75 and feedback gain 0.5; the feedback spike probability was capped at 1.0.

##### S13.4 Learning rules and controls

FSN and cFSN used the updates of Sections S4 and S5: the directed input and recurrent updates and the delta-readout update from the measured displacements, with $B_{\text{out}}$ learned by Kolen–Pollack mirroring (learning rate 0.012); cFSN centered the Kolen–Pollack activity product across the minibatch. The controls were:

**Supplementary Table 5 | Control methods for biologically local MNIST learning.**

| Method | Main difference from FSN |
|---|---|
| rFSN | $B_{\text{out}}$ fixed after random initialization [17] |
| CHFSN | Kolen–Pollack mirroring with the contrastive-Hebbian recurrent update of Section S6 [18,19] |
| Frozen recurrent | Readout learning only; input, recurrent and feedback parameters fixed |
| BPTT | Non-local surrogate-gradient reference |
| e-prop | Non-local online comparator with an adaptive feedback matrix and a LIF output layer |

##### S13.5 Frozen-recurrent control

The frozen-recurrent control trained only the output weights $V$ and output bias with the FSN delta-readout rule at learning rate 0.006, holding $U$, $W$, the recurrent bias and the feedback matrix at their initial values; both phases remained active. Architecture, seeds, data order, training duration and evaluation were matched to the FSN comparison, and the FSN-versus-frozen contrast therefore measures the joint contribution of input, recurrent and feedback plasticity beyond readout learning.

##### S13.6 BPTT comparator

BPTT used the same LIF forward model and spike-simulation duration, with surrogate threshold derivatives in the backward pass [6], and plain gradient-descent updates at learning rate 0.02 without weight decay or gradient clipping.

##### S13.7 E-prop comparator

The e-prop comparator [7] used the same architecture, $T = 100$, batch size, training length and spike evaluation protocol, with a spiking LIF output layer. Its gradient was computed by automatic differentiation with the recurrent spike input detached and the reset not differentiated (the GPU formulation of Bellec et al.), which gives the same gradient as the explicit eligibility-trace form. The pseudo-derivative was triangular with scale 0.3 and width 1.0, output traces used exponential moving averaging tied to the LIF output time scale, and the feedback matrix was adaptive, updated from the output learning signals and recurrent activity with learning rate 0.002. The learning rate was $2 \times 10^{-3}$; a check with learning rates of $10^{-3}$ and $3 \times 10^{-3}$ on the three plateau seeds and two other seeds did not improve the plateau seeds.

##### S13.8 Training protocol and hyperparameters

All methods used random seeds 0–9 with matched data order, without global gradient-norm clipping or an adaptive optimizer. Test accuracy was evaluated on all 10,000 test images in 79 batches of 128 images (the last containing 16 images) and reported as the unweighted mean of the 79 batch accuracies.

**Supplementary Table 6 | Hyperparameters for biologically local MNIST learning.**

| Hyperparameter | Value |
|---|---:|
| Training images | 60,000 |
| Test images | 10,000 |
| Seeds | 0–9 |
| Batch size | 128 |
| Recurrent neurons | 256 |
| Output neurons | 10 |
| Free-phase timesteps | 100 |
| Feedback-phase timesteps | 100 |
| Burn-in | 10 timesteps |
| Reset mode | subtractive |
| Input encoding | Bernoulli pixel spikes |
| Recurrent/output leak $\beta$ | 0.05 |
| Recurrent/output threshold $\theta$ | 1.0 |
| Logit scale | 20 |
| Local-rule learning rate | 0.006 |
| Kolen–Pollack learning rate | 0.012 |
| Nudge strength | 0.15 |
| Output feedback gain | 2.75 |
| Feedback gain | 0.5 |
| Maximum absolute weight | 3.0 |
| Local-rule training steps | 24,000 |
| Evaluation interval | 100 steps |
| BPTT learning rate | 0.02 |
| e-prop learning rate | 0.002 |

#### S13.9 Alignment and statistical analyses

All classification metrics were measured with spike simulation. The performance panels use test accuracy over training time, the best test accuracy across periodic evaluations and the number of training steps needed to reach test accuracies of 0.80, 0.90 and 0.95; final accuracy is a separate final spiking evaluation, whereas Fig. 3b shows the maximum periodic evaluation accuracy of each seed. If a run did not reach a threshold, the threshold step was set to the final training step rather than excluded.

For update-alignment diagnostics, each local-rule or e-prop update was compared with the raw surrogate-BPTT negative gradient computed on the same minibatch and the same stochastic spike realization (the random number generator state was restored after the diagnostic). Alignment was the cosine similarity between the actual parameter step, including any weight decay, and the negative gradient; the recurrent-layer update concatenated input weights, recurrent weights and recurrent biases, and the readout update concatenated output weights and biases. Mean cosines were averaged over all diagnostic evaluations across training within each seed. The frozen-recurrent control has no recurrent-layer update and is omitted from that comparison. Feedback–readout alignment was the cosine similarity between $B_{\text{out}}$ and $V$, and recurrent asymmetry was $\|W - W^\top\|_F / \|W\|_F$.

Comparisons were paired by seed, and each comparison against FSN used an exact Wilcoxon signed-rank test, with Holm correction within each metric. Final accuracy, best accuracy, asymmetry, training time and the steps to each threshold used six FSN-reference comparisons; recurrent-update alignment used five, because the frozen-recurrent control has no recurrent update, and feedback–readout alignment used five, because BPTT has no learned feedback matrix.

### S14 Supervised Reacher imitation with feedback-state carry

#### S14.1 Task and teacher–student design

The supervised Reacher experiment used Gymnasium Reacher-v5 with MuJoCo dynamics [12,13] in an imitation setting. A non-spiking teacher policy was first trained on a randomized-start Reacher task with Soft Actor–Critic (SAC) [20] and reached a deterministic evaluation reward of $-2.878$ over 128 episodes under the same reset condition. During RSNN training, the teacher supplied only a target action for the current observation. At each environment step $t$, the student observed $\boldsymbol{o}_t$, ran a free spiking phase to produce the action $\boldsymbol{a}_t$, stepped the environment with $\boldsymbol{a}_t$, and then ran a feedback phase driven by the difference between the teacher action $\boldsymbol{a}_t^{\mathrm{targ}}$ and $\boldsymbol{a}_t$. The feedback phase produced the local weight updates and the recurrent state carried to the next observation.

#### S14.2 Randomized Reacher environment

At reset, the standard Reacher-v5 reset was called and the MuJoCo state was then modified: the two arm joint positions were sampled from $U(-2, 2)$ radians and the two joint velocities from $U(-0.05, 0.05)$; the default target sampling was retained. Actions were clipped to the environment bounds. Training used 256 parallel environments and 64 environment steps per update (16,384 transitions per update). Evaluation used 64 parallel environments and 64 completed deterministic episodes per checkpoint. Each configuration was trained on ten matched seeds.

#### S14.3 Teacher policy

The teacher was the frozen actor of the SAC policy: two fully connected ReLU hidden layers of 256 units and a tanh action head, receiving the raw 10-dimensional observation and producing a 2-dimensional action in normalized coordinates. The teacher action never stepped the environment; it supplied the output teaching signal during training and in the feedback-control evaluation, and served only to calculate the imitation error in the evaluation without teaching feedback.

#### S14.4 RSNN student architecture

Observations were clipped to $[-10, 10]$, divided by an observation-rate scale of 2 and split into positive and negative channels,

$$\boldsymbol{q}(\boldsymbol{o}) = [\mathrm{ReLU}\left(\mathrm{clip}(\boldsymbol{o}/2, -1, 1)\right), \mathrm{ReLU}\left(-\,\mathrm{clip}(\boldsymbol{o}/2, -1, 1)\right)],$$

giving 20 signed-rate channels, each represented by 96 redundant Bernoulli sensory axons (effective fan-in 1,920). The recurrent layer contained 256 LIF neurons, and the action readout used 16 positive and 16 negative LIF neurons per action dimension (64 output neurons). The output rates were decoded as $y_k = \bar{r}_k^+ - \bar{r}_k^-$, where $\bar{r}_k^{\pm}$ are the mean rates of the positive and negative populations of action dimension $k$, and the normalized action was $a_k = \tanh(2.5 y_k)$.

**Supplementary Table 7 | Architecture and LIF parameters for supervised Reacher imitation.**

| Parameter | Value |
|---|---|
| Raw observation dimension | 10 |
| Signed encoded channels | 20 |
| Redundant sensory axons per channel | 96 |
| Effective sensory axons | 1,920 |
| Recurrent LIF neurons | 256 |
| Action dimensions | 2 |
| Readout population size | 16 positive + 16 negative per action |
| Output LIF neurons | 64 |
| Free-phase timesteps | 150 |
| Feedback-phase timesteps | 150 |
| Burn-in timesteps | 15 |
| Reset mode | zero reset |
| Recurrent/output leak $\beta$ | 0.05 / 0.05 |
| Recurrent/output threshold | 1.0 / 1.0 |
| Recurrent/output tonic current | 0.05 / 0.16 |
| Input/recurrent/output weight scale | 0.70 / 0.08 / 0.70 |
| Feedback-output/feedback-recurrent scale | 0.70 / 0.08 |
| Maximum absolute weight | 3.0 |

#### S14.5 Free phase and feedback phase

At each environment step, the student first ran a free phase without feedback current, which produced the free recurrent and output rates $\boldsymbol{r}^{h,-}$ and $\boldsymbol{r}^{o,-}$ and the action used to step the environment. The supervised loss compared the decoded student action with the teacher action,

$$L = 4\ \mathrm{MSE}(\boldsymbol{a}_t, \boldsymbol{a}_t^{\mathrm{targ}}) + 3 \times 10^{-4}\ \mathrm{mean}\left[(\boldsymbol{r}^{o,-} - 0.25)^2\right],$$

where the second term weakly regularizes the output rates toward a homeostatic target. Automatic differentiation of this loss was used only to obtain the output teaching signal $\boldsymbol{\lambda}^o = \partial L / \partial \boldsymbol{r}^{o,-}$, which was not backpropagated through the recurrent network but converted into signed feedback spikes. During the feedback phase, signed Bernoulli events perturbed the output population (feedback spike scale 0.04, nudge strength $\epsilon = 0.04$, feedback gain 0.5, output feedback gain 1.0, 384 redundant output-feedback axons), the output displacement was converted into output-feedback events and projected through $B_{\mathrm{out}}$ with a one-step causal delay, and the recurrent displacement was measured as in Section S4. The feedback phase reused the sensory input spike sequence of the free phase.

#### S14.6 Feedback-state carry

The environment action was always decoded from the free phase, but the membrane and spike state at the end of the feedback phase initialized the next environment step. Training-time evaluations used the same carry with fixed weights; the checkpoint evaluations described below omitted the feedback phase and carried the free-phase state.

#### S14.7 Local updates and Kolen–Pollack mirroring

Input, recurrent and readout weights used the FSN updates of Section S4 with nudge strength $\epsilon = 0.04$, and cFSN learned $B_{\mathrm{out}}$ with the centered Kolen–Pollack rule of Section S5. Table 8 lists the configurations.

#### S14.8 Compared methods

Six configurations shared the task, teacher, architecture, phase lengths, feedback-spike scale, learning rate and evaluation protocol, and differed in the feedback pathway or recurrent update rule.

**Supplementary Table 8 | Reacher methods.**

| Method | Difference from cFSN |
|---|---|
| Centered FSN (cFSN) | Proposed method: directed recurrent update, delta readout, centered Kolen–Pollack mirroring |
| FSN | Kolen–Pollack mirroring without centering |
| Random-feedback FSN (rFSN) | Fixed random feedback matrix; no Kolen–Pollack mirroring [17] |
| Frozen recurrent | Input and recurrent weights held fixed; readout and feedback learning retained |
| CHFSN | Contrastive-Hebbian recurrent update (Section S6) instead of the directed displacement update |
| cCHFSN | Centered contrastive-Hebbian recurrent update |

The Reacher CHFSN and cCHFSN controls changed the recurrent rule while retaining the delta readout; cCHFSN centered its recurrent contrastive products and Kolen–Pollack update.

#### S14.9 Optimization and diagnostics

Local increments were accumulated over the 64-step parallel rollout and applied once. The forward and output learning rates were $1.5 \times 10^{-4}$, the rollout gradient scale 16, weight decay $3 \times 10^{-6}$ and global clipping of the accumulated local update 1; Adam preconditioning [9] was applied to the measured local updates. The Kolen–Pollack update used twice the forward learning rate with the same preconditioning and feedback weight decay $3 \times 10^{-6}$. Training ran for 80,000,000 environment transitions (4,883 updates of 256 environments × 64 steps, the last rollout shortened). Training-time evaluation occurred every 20 updates over 64 episodes with fixed weights and feedback-state carry (246 evaluations); the checkpoint with the highest logged reward was saved as the best checkpoint, and the final checkpoint contained the weights after update 4,883.

For diagnostics, a surrogate-BPTT reference gradient [6] was computed every 20 updates on 64 stored examples without training the model. The recurrent-layer cosine in Extended Data Fig. 4 concatenates the updates of the input weights, recurrent weights and recurrent biases. Feedback–readout alignment was the centered cosine similarity between $B_{\text{out}}$ and the readout matrix.

#### S14.10 Evaluation without teaching feedback

The best and final checkpoints of all six methods and ten seeds were evaluated on 512 fresh episodes per checkpoint (256 parallel environments, two episodes each) with randomized resets from the training distribution and 50 environment steps per episode. The reward plotted in Fig. 4c is the selected checkpoint's mean total reward per episode over these episodes; logged maxima were used only to select the checkpoints, and the training-time curves in Fig. 4b retain the feedback-state-carry protocol. In this evaluation, each action and the carried recurrent state came from the 150-timestep free phase, the feedback phase was never run, weights were fixed and the recurrent state was reset at episode boundaries. A control evaluation ran the teacher-driven feedback phase and carried its final state without weight updates; both modes applied the student's free-phase action. All methods, seeds, checkpoints and evaluation modes used the same 512 initial conditions, with reset seeds unused in training and the same input-spike random seed at the start of each evaluation.

##### S14.11 Statistical analysis

Checkpoint rewards were averaged over the 512 episodes for each trained seed, which was the unit of analysis. Exact Wilcoxon signed-rank tests across matched seeds were Holm-corrected over all 15 method pairs, separately for the best and final checkpoints. The comparison with versus without teaching feedback used paired seed-level rewards with Holm correction across the six methods for each checkpoint. The gradient-alignment diagnostics used Holm correction over the ten pairs among the five plotted methods; the recurrent-update and centered feedback–readout cosines are final-checkpoint diagnostics. For runs in which the diagnostic became undefined during training, the last finite value was used.

#### S15 Biologically local Ant reinforcement learning

##### S15.1 Task and training objective

The reinforcement-learning experiment with FSN trained a spiking recurrent actor–critic on Gymnasium MuJoCo Ant-v5 (105-dimensional observation, 8-dimensional bounded action) [12,13] with PPO and GAE [14,15]. The actor–critic RSNN selected actions during a free spiking phase; during training, PPO supplied an output-rate teaching signal that was delivered into the network as signed feedback events, and the resulting rate displacements drove the local updates of Section S4. cFSN, in which the feedback matrix is learned by the centered Kolen–Pollack rule, was the primary method and was compared with FSN, rFSN, CHFSN and cCHFSN under the same task, rollout geometry, observation encoding, architecture and evaluation procedure.

##### S15.2 PPO rollout and advantage computation

Training used 256 parallel environments and 64 environment steps per rollout (16,384 transitions per PPO update), divided into 32 minibatches of 512 transitions and optimized for 2 epochs. Runs used seeds 0–9 and were trained for 1,220 full rollouts (19,988,480 transitions), the largest whole number of rollouts within 20 million environment steps. For each transition, the normalized observation, sampled action, old log probability, value estimate, reward, termination indicator, initial recurrent state and free-phase input spike sequence were stored. Advantages used GAE with $\gamma = 0.99$ and $\lambda = 0.95$; PPO used clip coefficient 0.2, value coefficient 1.5, entropy coefficient 0, target KL 0.03 and maximum gradient norm 1.0. Rewards were normalized and clipped to $[-10, 10]$, and observations were normalized with a running mean and variance and clipped to $[-10, 10]$.

The PPO loss was not backpropagated through the recurrent spiking dynamics. During each minibatch update, the stored input spike sequence and initial recurrent state were replayed to reproduce the free output rate $\boldsymbol{r}_t^{o,-}$, and the loss was differentiated only with respect to this vector,

$$\boldsymbol{\lambda}_t^o = \frac{\partial L_{\mathrm{PPO},t}}{\partial \boldsymbol{r}_t^{o,-}},$$

which was the interface between PPO and the spiking circuit.

##### S15.3 Observation-to-spike encoding

After normalization, each observation coordinate was divided by 3, clipped to $[-1, 1]$ and split into positive and negative channels,

$$\boldsymbol{q}_t = [\mathrm{ReLU}(\mathrm{clip}(\tilde{\boldsymbol{o}}_t/3, -1, 1)), \mathrm{ReLU}(-\,\mathrm{clip}(\tilde{\boldsymbol{o}}_t/3, -1, 1))],$$

giving 210 signed channels, each represented by 32 redundant Bernoulli sensory axons (6,720 input axons, input gain 1.0).

##### S15.4 Spiking recurrent actor–critic

The actor and critic shared a recurrent LIF population of 256 neurons, and the output layer used a positive/negative population code with 16 LIF neurons per sign for each of the 8 action coordinates and the value, that is, 288 output LIF neurons. At internal timestep $\tau$, the recurrent current was $\boldsymbol{h}^h_{t,\tau} = U\boldsymbol{z}_{t,\tau} + W\boldsymbol{s}^h_{t,\tau-1} + \boldsymbol{b}^h + \boldsymbol{f}^h_{t,\tau}$ and the output current $\boldsymbol{h}^o_{t,\tau} = V\boldsymbol{s}^h_{t,\tau} + \boldsymbol{b}^o + \boldsymbol{f}^o_{t,\tau}$, with feedback currents $\boldsymbol{f}$ equal to zero in the free phase. Recurrent and output neurons used zero-reset LIF dynamics (Methods) with $\beta = 0.05$ and $\theta = 1.0$. The recurrent and output tonic currents were initialized to 0.05 and 0.16; input, recurrent, output, recurrent-feedback and output-feedback weight scales were 0.70, 0.08, 0.70, 0.08 and 0.70; self-recurrent weights were zeroed, weights were clipped to maximum absolute value 3, and forward weight decay was $10^{-6}$.

##### S15.5 Free phase, feedback-state carry and environment interaction

At each environment step, the network ran a 225-timestep free phase; the first 22 timesteps were discarded as burn-in, and the free phase alone determined the action. For each action coordinate, the signed action feature was the positive-population rate minus the negative-population rate, converted into the pre-tanh Gaussian policy mean by a learnable positive scale initialized at 4.0; the critic value used a scale initialized at 15.0, and the policy used a diagonal Gaussian in pre-tanh space with learnable log standard deviation initialized at $-0.5$.

After the action was computed, the model ran a 225-timestep feedback-phase window and carried its final membrane and spike state into the next free phase (feedback-state carry), so that the recurrent state at step $t+1$ reflected both the sensory-driven free-phase activity at step $t$ and the feedback-phase dynamics that followed it. During rollout and evaluation, this window received no teaching signal, because the PPO loss is available only after the rollout. During PPO updates, the same feedback-phase machinery was driven by the PPO-derived teaching signal and produced the local updates; each minibatch replayed individual stored transitions with their stored initial states and input spikes, and interaction then resumed from the retained final state of the actual rollout with the updated weights. Recurrent states were detached between environment steps and reset to zero at episode boundaries.

##### S15.6 Feedback-spike conversion and local plasticity

Before feedback delivery, the actor and critic components of $\boldsymbol{\lambda}^o_t$ were each scaled by 1.5, the vector was normalized to a global root mean square of 0.75 and clipped to $[-4, 4]$. The teaching signal was converted into signed Bernoulli feedback events with event scale 0.04 and probability cap 1.0, using 384 redundant signed feedback axons per output channel. The output displacement was converted into output-delta feedback spikes and projected through $B_{\mathrm{out}}$ to the recurrent population with one axon per feedback channel and a causal one-timestep delay. The measured displacements and the FSN, delta-readout and centered Kolen–Pollack updates were those of

Sections S4 and S5 with nudge strength $\epsilon = 0.04$; FSN removed the centering, rFSN kept the feedback matrix fixed after random initialization, and CHFSN and cCHFSN replaced the directed displacement and delta-readout updates with the contrastive updates of Section S6. Adam preconditioning [9] was applied to the forward-weight and feedback updates. The forward-weight learning rate was $7 \times 10^{-5}$, the Kolen–Pollack learning-rate scale 1 and gradient scale 0.8, the recurrent and output forward-weight gradient scales 0.8, and forward and feedback weight decay $10^{-6}$.

#### S15.7 Evaluation and statistical analysis

Evaluation was performed every 20 PPO updates with 64 parallel environments and 64 completed deterministic episodes, using the tanh-squashed Gaussian mean action. The primary outcome was the deterministic evaluation reward (total reward per episode); the forward-reward component was also recorded. Time-course plots show the seed mean and SEM. Comparisons used exact Wilcoxon signed-rank tests across matched seeds, with Holm correction across all ten pairs among the five methods separately for each metric.

#### S15.8 Hyperparameters

**Supplementary Table 9 | Hyperparameters for biologically local Ant reinforcement learning.**

| Parameter | Value |
|---|---|
| Environment | Ant-v5 |
| Seeds | 0–9 |
| Environment steps | 19,988,480 (1,220 rollouts) |
| Parallel environments | 256 |
| Rollout length | 64 |
| PPO batch size | 16,384 |
| Minibatches / minibatch size | 32 / 512 |
| PPO epochs | 2 |
| PPO learning rate | $7 \times 10^{-5}$ |
| Clip coefficient | 0.2 |
| GAE $\gamma$, $\lambda$ | 0.99, 0.95 |
| Value coefficient / entropy coefficient | 1.5 / 0 |
| Target KL / max gradient norm | 0.03 / 1.0 |
| Observation normalization | running mean/variance, clip 10 |
| Reward normalization | enabled, clip 10 |
| Observation rate scale / input gain | 3 / 1 |
| Input mode | signed Bernoulli spikes |
| Redundant sensory axons | 32 per signed channel |
| Recurrent LIF neurons | 256 |
| Output LIF neurons | 288 |
| Free phase / feedback phase | 225 / 225 timesteps |
| Burn-in | 22 timesteps |
| Recurrent/output leak $\beta$ | 0.05 / 0.05 |
| Recurrent/output threshold $\theta$ | 1.0 / 1.0 |
| Reset mode | zero reset |
| Readout scheme | positive/negative population code |
| Population size | 16 neurons per sign |
| State carried between environment steps | feedback-phase final state |
| Initial action/value scale | 4 / 15 |
| Learnable log standard deviation | initialized at $-0.5$ |
| Nudge strength | 0.04 |
| Feedback mode | signed Bernoulli |
| Feedback spike scale / max probability | 0.04 / 1.0 |
| Output feedback axons | 384 per output channel |

| Parameter | Value |
|---|---|
| Recurrent feedback axons | 1 per feedback channel |
| Output feedback signal | output-rate error |
| Recurrent feedback signal | output-displacement spikes |
| Main forward-weight rule | directed recurrent update, delta readout |
| Main feedback rule | centered Kolen–Pollack rule |
| Forward-weight optimizer | Adam |
| Feedback optimizer | same as forward-weight optimizer |
| Forward / feedback weight decay | $10^{-6}$ / $10^{-6}$ |
| Teaching-signal normalization / clipping | global RMS 0.75 / $[-4, 4]$ |
| Actor / critic teaching-signal scale | 1.5 / 1.5 |
| Evaluation interval | every 20 PPO updates |
| Evaluation episodes | 64 deterministic episodes |

### S16 Biological spine-data validation of the local-product rule

#### S16.1 Dataset and measured variables

We analyzed the open motor-learning dataset of Hedrick, Wright and Komiyama [21], which contains trial-aligned synaptic activity, dendritic activity, structural spine measurements and later spine fate in the same preparation. Mice underwent daily rewarded lever-press training for 14 days, with imaging on days 1–3, 6–8 and 11–13; each field was imaged at five-day intervals, and the analysis paired early with middle and middle with late sessions. Spine-local iGluSnFR3 activity, a glutamate indicator signal [22], was used as a proxy for presynaptic activity. RCaMP2 activity, a calcium indicator signal [23] pooled from shaft regions of interest along the parent dendritic branch in the source study, was used as a proxy for postsynaptic activity; the same branch-level trace was paired with each spine on that branch. The apparent spine area, obtained in the source study from integrated spine fluorescence normalized by nearby shaft fluorescence within 20 $\mu$m and corrected for pixel size, was used as a structural proxy for synaptic strength, and spine retention or elimination at the next session was a separate outcome.

#### S16.2 Reward-aligned variables

For each spine $j$, session $s$ and rewarded trial $k$, four event-onset rates were extracted: the pre-reward ($a_{jsk}$) and post-reward ($b_{jsk}$) iGluSnFR3 spine event-onset rates and the pre-reward ($x_{jsk}$) and post-reward ($y_{jsk}$) parent-shaft RCaMP2 event-onset rates. Activity was sampled at 58.3 Hz, and the binary active/inactive classifications published with the source study were used: its pipeline estimated the fluorescence noise from the standard deviation of negative $\Delta F/F_0$ values mirrored about zero, detected an active interval when the 1-s LOESS-smoothed trace crossed twice the noise estimate for iGluSnFR3 or three times for RCaMP2, and extended the onset backward to the crossing of a baseline threshold equal to the noise estimate in the unsmoothed trace [21]. An event onset was a transition from inactive to active in this binary trace; the first frame was assigned no onset, and every detected event counted once, without amplitude weighting. The event-onset rate was the number of onsets in the interval divided by its duration.

The primary analysis used matched 2.5-s pre- and post-reward intervals with a 0.1-s guard on each side of reward, $[t_{\text{reward}} - 2.6\,\text{s}, t_{\text{reward}} - 0.1\,\text{s})$ and $[t_{\text{reward}} + 0.1\,\text{s}, t_{\text{reward}} + 2.6\,\text{s})$, for both sensors. Only rewarded trials in which the mouse obtained the reward by a cue-initiated lever movement were analyzed: a reward was included if a cue onset occurred within 11 s and a lever-movement onset within 10 s before reward delivery. Rewards within 4 s of the start or 2.1 s of the end of the recording, and rewards whose pre- or post-reward interval extended outside the recording, were excluded. In the 16 sessions analyzed, 694 of 722 rewards met these criteria.

Before calculating an update product, rates were standardized within each dendritic-branch × learning-stage stratum using the median and median absolute deviation (MAD): the iGluSnFR3 pre- and post-reward rates were pooled and transformed as

$$z = \frac{r - \mathrm{median}(r)}{1.4826\ \mathrm{median}(|r - \mathrm{median}(r)|)},$$

and the RCaMP2 rates were standardized in the same way with their own median and scale, so that the pre–post difference of each sensor is preserved on a common scale. Median/MAD standardization was used because event-onset rates were sparse with occasional high values. If the denominator was zero or undefined, it was replaced by the standard deviation of the pooled values, and if that was also zero or undefined, by 1. The symbols $a$, $b$, $x$ and $y$ below refer to these standardized rates. For each rule, the product was calculated separately for every rewarded trial with finite values for all four rates and then averaged within each spine transition.

The FSN product, $U^{\mathrm{FSN}}_{jsk} = a_{jsk}(y_{jsk} - x_{jsk})$, is presynaptic input before reward multiplied by the reward-aligned displacement of the postsynaptic signal, the biological-data counterpart of the feedback-spike update.

##### S16.3 Comparison update rules

The FSN product was compared with Hebbian-like and contrastive alternatives computed from the same variables; the rule names are those used in Fig. 6.

**Supplementary Table 10 | Local-product and comparator rules evaluated in the spine-data analysis.**

| Rule (label in Fig. 6) | Formula | Interpretation |
|---|---|---|
| FSN product | $a(y - x)$ | pre-reward iGluSnFR3 rate × reward-locked RCaMP2 displacement |
| CHL | $by - ax$ | contrastive Hebbian learning: post-reward minus pre-reward coactivity |
| Plus Hebb | $by$ | post-reward iGluSnFR3 × RCaMP2 coactivity |
| Minus Hebb | $ax$ | pre-reward iGluSnFR3 × RCaMP2 coactivity |
| Delta-pre × delta-post | $(b - a)(y - x)$ | iGluSnFR3 change × RCaMP2 change |
| CHL extra | $x(b - a)$ | pre-reward RCaMP2 rate × iGluSnFR3 change; the term that, added to the FSN product and to delta-pre × delta-post, completes CHL |
| Post-pre timing control | $b(y - x)$ | post-reward iGluSnFR3 rate × RCaMP2 change |

##### S16.4 Spine-area outcome analysis

The primary structural outcome was the signed spine-area change between successive imaging sessions among spines present at both, $\Delta A_{j,s} = A_{j,s+1} - A_{j,s}$, where $A_{j,s}$ is the area of spine $j$ in session $s$; positive values indicate enlargement. Spine area was reconstructed from the integrated spine-fluorescence measure and the imaging zoom published with the source study, following its field- and session-matching procedure [21]: for each field and session, the area was the integrated measure divided by $(0.5\,z)^2$, where $z$ is the imaging zoom of that session. The primary 2.5-s analysis contained $n = 517$ surviving spine transitions from seven mice.

For each rule, Spearman and Pearson correlations were calculated between the transition score and the future spine-area change across the same 517 transitions; the Spearman correlation was the primary unadjusted statistic, and Fig. 6c displays its raw $p$ values. For the grouped visualization, scores were classified as negative, exactly zero or positive independently of the outcome; the plotted point of each group is the pooled mean future area change at the group's median score, with 95% percentile confidence intervals from 10,000 mouse-cluster bootstrap resamples (seven mouse identities sampled with replacement, all transitions of each selected mouse retained). The least-squares line used all 517 transitions, with a pointwise mean-response confidence band from the mouse-clustered covariance and Student's $t$ distribution with six degrees of freedom. The pooled correlation tests do not use mouse-clustered inference.

##### S16.5 Controlled linear models

To test whether each rule predicted future spine-area change after accounting for measured covariates, separate ordinary-least-squares models of the form

$$\Delta A_{j,s} = \beta_0 + \beta_m \widetilde{U}_{m,j,s} + \beta_2 A_{j,s} + \beta_3 C_{j,s} + \beta_4 M_{j,s} + \beta_5 P_{j,s} + \epsilon_{j,s}$$

were fitted, where $\widetilde{U}_{m,j,s}$ is the transition score of rule $m$ standardized by its mean and sample standard deviation across all 563 transitions, $A_{j,s}$ the natural logarithm of the current spine area, $C_{j,s}$ the nearby-spine coactivity within $10\,\mu$m (mean normalized coactivity with simultaneously imaged neighboring spines, with coactivity onsets normalized by the geometric mean of the two spines' event rates), $M_{j,s}$ the binary movement-related-spine classification of the source study [21] and $P_{j,s}$ an indicator of the middle-to-late rather than early-to-middle transition. Current log area and nearby coactivity were standardized by their medians and 1.4826 times their MADs. Each model contained the 517 transitions from seven mice; the covariance matrix used a finite-sample-corrected sandwich estimator clustered by mouse, and each coefficient was tested against zero with a mouse-clustered Wald $t$ test on six degrees of freedom. The rule coefficient plotted in Fig. 6d is the adjusted difference in future area change per standard deviation of the transition score; coefficient $p$ values were not adjusted for multiple testing.

##### S16.6 Spine survival and elimination analysis

All $n = 563$ transitions with a finite score were included: 517 ending in survival and 46 in disappearance. Transition scores were compared with a one-sided Wilcoxon rank-sum test with the directional alternative that scores were greater for surviving spines (asymptotic $p$ value with continuity correction). This test is transition-pooled and does not adjust for clustering by mouse.
